# Handhold-mediated strand displacement: a nucleic acid-based mechanism for generating far-from-equilibrium assemblies through templated reactions

**DOI:** 10.1101/2020.05.22.108571

**Authors:** Javier Cabello-Garcia, Wooli Bae, Guy-Bart V. Stan, Thomas E. Ouldridge

**Affiliations:** Department of Bioengineering and Centre for Synthetic Biology, Imperial College London, London, U.K.

## Abstract

Toehold-mediated strand displacement (TMSD) is a nucleic acid-based reaction wherein an invader strand (*I*) replaces an incumbent strand (*N*) in a duplex with a target strand (*T*). TMSD is driven by toeholds, overhanging single-stranded domains in *T* recognised by *I*. Although TMSD is responsible for the outstanding potential of dynamic DNA nanotechnology^1, 2^, TMSD cannot implement templating, the central mechanism by which biological systems generate complex, far-from equilibrium assemblies like RNA or proteins^3, 4^. Therefore, we introduce handhold-mediated strand displacement (HMSD). Handholds are toehold analogues located in *N* and capable of implementing templating. We measure the kinetics of 98 different HMSD systems to demonstrate that handholds can accelerate the rate of invader-target (*IT*) binding by more than 4 orders of magnitude. Furthermore, handholds of moderate length accelerate reactions whilst allowing detachment of the product *IT* from *N*. We are thus able to experimentally demonstrate the use of HMSD-based templating to produce highly-specific far-from-equilibrium DNA duplexes.

---

Since their first description by Yurke et al.^5^, toeholds have been the main regulatory motif for DNA strand displacement (**Fig. 1a**). The versatility and ease of design of toeholds have made TMSD a great building block for constructing all sort of chemical reaction networks^6, 7^. TMSD reaction rates increase exponentially with toehold length until saturating at approximately six orders of magnitude faster than toehold-free displacement, at a toehold length of 6-7 nucleotides (nt)^8^. Toeholds increase TMSD reaction rates by inhibiting the detachment of the *I*-strand from *T* during the competition with *N*^9^.

**Fig. 1:**
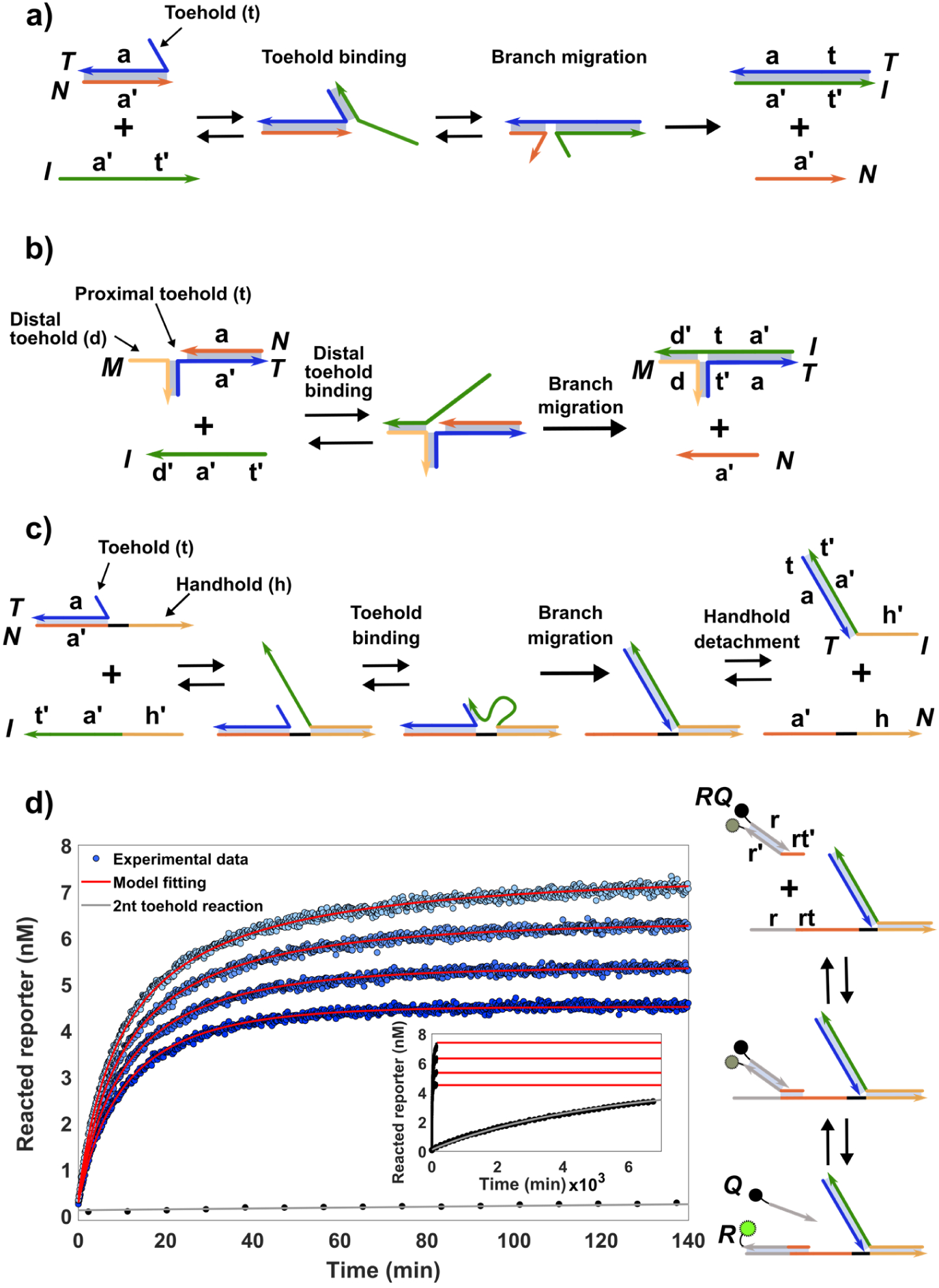
Strand displacement topologies. **A) Toehold-mediated strand displacement.** The presence of a toehold (t) in the target strand (*T*) mediates the dsplacement of the incumbent strand (*N*) by the invader strand (*I*). **B) Associative toehold.** A reaction with a small proximal toehold (t) is accelerated by an auxiliary strand (*M*) complementary to the strands *I* and *T*. This mechanism effectively increases the length of t. However, it results in an *IT* product with the *M* strand strongly attached. **C) Handhold-mediated strand displacement (HMSD).** The incorporation of a handhold (h) – an independent overhang in *N* – spatially constrains *I* to the vicinity of *T*, enhancing the efficiency of binding to short proximal toeholds in the target. This binding results in faster reaction rates while, for short handholds, allowing the complete detachment of *IT* after the reaction. **D) Handhold-complementarity substantially enhances displacement rates for short toeholds.** A reporter (*RQ*) monitors the progress of HMSD reactions by interacting with the displaced domain in *N* and fluorescing. We plot representative results for a reaction with a 2 nucleotide (nt) proximal toehold and a 7 nt handhold. All traces are fitted with a single set of reaction parameters, as outlined in **Supplementary Note XVI**. Conditions: [*RQ*]_0_=15 nM, [*TN*]_0_=10 nM, [*I*]_0_=[7-4] nM, 1M NaCl in 1x TAE at 25°C. Inset: Kinetics for 6nM of the *I*-strand without handhold-complementarity (grey).

The rationale behind TMSD reactions is that the *I*-strand can specifically recognise a toehold in *T*, and then bind to further bases in *T* located adjacently to the toehold (the displacement domain). The toehold recognition interaction is retained intact in the *IT* product because the toehold and displacement domain act cooperatively to bind *I* and *T* together. More complex TMSD topologies have been demonstrated where toeholds are not adjacent to the displacement domain, such as the associative toehold^10^ (**Fig. 1b**) and remote toehold^11^ structures. However, the TMSD rationale remains: the toehold binding acts cooperatively with the displacement domain, so that the toehold remains sequestered in the product. Toehold sequestering prevents TMSD from implementing complex functionalities, like templating of far-from-equilibrium assemblies.

Templating is a key functionality in biology, responsible for sequence-based information transmission in the processes that are central to the central dogma of molecular biology^3, 4, 12^. In these processes, a pool of molecules assemble into a complex structure – such as a polynucleotide or polypeptide – with their sequence determined by that of a template^13^. For example, during RNA transcription, ribonucleotides use sequence-specific recognition interactions with a DNA template to polymerise into a sequence-complementary RNA strand^3^. The RNA sequence is thus exclusively determined by the template information transmitted by the recognition interactions, rather than the non-specific backbone bonds between RNA nucleotides. However, to be functional, the product RNA must eventually break all recognition interactions with the template and detach. As a result, the recognition interactions cannot shift the equilibrium state of the eventual product towards the template-complementary sequence,^14, 15^. Accurately assembled products are, therefore, necessarily produced far-from-equilibrium, i.e. far from the random sequence distribution obtained in equilibrium.

Without templating of far-from-equilibrium ensembles it would be impossible to achieve the chemical complexity of biology. For example, there simply cannot be enough information stored in the interactions between 20 basic building blocks (the standard amino acids) to code for the selective assembly of tens of thousands of specific proteins – and only those proteins – from a mixture of only these amino acids^16^. Far-from-equilibrium templating solves this problem by providing a reusable mechanism to assemble an arbitrary polypeptide chain.

Given the fundamental significance of far-from-equilibrium templating, it is desirable to implement similar functionality in synthetic DNA-based reactions. However, TMSD is ill-suited for operating far from equilibrium because toehold interactions, which are the source of specific recognition, cannot be transient; binding cooperativity keeps the toehold sequestered after the displacement. Hitherto, attempts at templating using TMSD have required external intervention or non-chemical mechanisms, such as heating cycles^17^, hindering their implementation in autonomous reaction networks. Other synthetic copying systems have faced similar challenges^18-20^.

Here, we introduce handhold-mediated strand displacement (HMSD, **Fig. 1c**), a new strand displacement topology capable of implementing templating in DNA reaction networks. HMSD accelerates the binding of *I* and *T* through a handhold domain in *N*, rather than a toehold in *T*. The proposed reaction mechanism works as follows: the *I-*strand with handhold complementarity (*I*_*hc*_) recognises and binds to the handhold, producing a transient 3-stranded complex that colocalises *I*_*hc*_ and the duplex *TN*. Then, *I*_*hc*_ displaces *N* from *T* producing the product *IT.* A short proximal toehold between *I*_*hc*_ and *T* further accelerates the displacement step. The formation of *IT* destabilises the interaction between *N* and *T*, eliminating cooperativity. Consequently, the product *IT* is now able to detach from *N*, breaking the handhold recognition interaction and thus enabling templating.

In this letter, we first demonstrate the displacement rate increment, by several orders of magnitude, when using *I*_*hc*_ over *I-*strands without handhold complementarity (*I*_*wc*_). We then show that it is possible to both increase the displacement rate and allow the *IT* product to detach from *N* if handholds of moderate length are used. Finally, we demonstrate production of specific, far-from-equilibrium *IT* complexes by handhold-based templating.

We first test the influence of different handhold and proximal toehold lengths on the strand displacement kinetics. We monitor strand displacement with the aid of an external reporter DNA duplex (*RQ*) labelled with a quenched fluorophore, minimising fluorophore interference^21^ in the reaction system. As shown in **Fig. 1d**, *N* triggers *RQ* by conventional TMSD, producing fluorescence only after *IT* formation. Typical HMSD reactions consist of adding *I*_*hc*_ to a solution containing *TN* and *RQ*. We also compare HMSD kinetics to those of a TMSD reaction triggered by *I*_*wc*_ in the same conditions.

We analyse the data by modelling HMSD as a two-step process, where *I*_*hc*_ reversibly attaches to the handhold and, once bound, triggers the displacement (**Fig. 1c**):

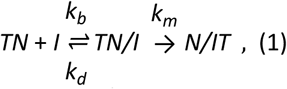

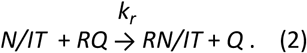

Here, “*X/Y*” indicates that species *“Y”* is bound to the handhold of *“X”. k*_*b*_ and *k*_*d*_ are rate constants for 2^nd^ order binding and 1^st^ order detachment of *I*_*hc*_ from *N*’s handhold, respectively. *k*_*m*_ is the 1^st^ order rate constant for the strand displacement once *I*_*hc*_ is bound to *N* by its handhold. *k*_*r*_ is the 2^nd^ order rate constant for reporter *RQ* activation. We treat all the displacement reactions as effectively irreversible, even in the absence of a proximal toehold because *RQ* acts as a sink for the displacement products.

We complement the reaction model by introducing TMSD as a 2^nd^ order competing reaction with rate *k*_*t*_. See **Supplementary Note XI** for the full implementation of the reaction model as an ODE system. We numerically solve the ODE system and use the results to fit a single set of *k*_*d*_, *k*_*m*_ and *k*_*b*_ to the fluorescent traces produced for each reaction system at a range of *I*_*hc*_ concentrations. *k*_*r*,_ the constraints for *k*_*b*_, and the competing TMSD rate *k*_*t*_ are obtained from specific experiments that probe their values in isolation (See *Materials and Methods*). The fitted results (**Supplementary Table 15**) reveal that HMSD operates in three different regimes: 1^st^ order, and 2^nd^ order with either reversible or irreversible handhold-binding. For the HMSD systems tested, the most common regime is the 2^nd^ order regime with reversible handhold-binding, occurring when *k*_*b*_[*I*]_0_<*k*_*d*_, *k*_*m*_, with [*I*]_0_ as the initial invader concentration. In this regime, *TN/I* is short-lived, resulting in a 2^nd^ order displacement rate (*k*) of the form:

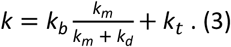

The 2^nd^ order regime with irreversible handhold-binding occurs for systems with handholds longer than 8 nt (ΔG°_*binding*_<-12.39 Kcal/mol^22^) and proximal toeholds of 1 nt or more (*k*_*d*_ <*k*_*b*_[*I*]_0_<*k*_*m*_). Here, all handhold-binding events result in rapid displacement. In this regime, *k* is insensitive to changes in *k*_*m*_ and *k*_*d*_, resulting in *k*≈*k*_*b*_. The 1^st^ order regime is identified in systems with handholds lengths above 8 nt and no proximal toehold (*k*_*b*_[*I*]_0_>*k*_*d*_,*k*_*m*_; *k*_*t*_≈0,). In this regime, the formation of *TN/I* is effectively irreversible, but the reaction rate is limited by *k*_*m*_, resulting in *k* being insensitive to *k*_*b*_ and *k*_*d*_.

Given the existence of 1^st^ and 2^nd^ order regimes, rate constants cannot be used directly to compare the speed of the different HMSD systems tested. Instead, we use the fitted rate constants to calculate, as described in *Materials and Methods*, each reaction’s half-life (*t*_*1/2*_) as a comparable measure of its reaction speed (**Fig. 2a**). Handhold complementarity decreases *t*_*1/2*_ by several orders of magnitude, analogously to toeholds in conventional TMSD. At saturating handhold lengths (>8 nt) and in the presence of a proximal toehold, the obtained *t*_*1/2*_ is consistent with conventional TMSD at saturating toehold lengths. In **Fig. 2b**, we report the half-life for systems triggered with 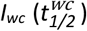. As expected, these systems only react by conventional TMSD, and 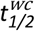 *i*s insensitive to handhold presence and exclusively determined by proximal toehold length. We plot the ratio 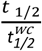 for every system in **Fig. 2c** to estimate the contribution of handholds to the kinetics. The handhold contribution is especially notable for proximal toeholds below 3 nt and handholds longer than 6 nt, where handhold-complementarity can reduce *t*_*1/2*_ by four orders of magnitude. this reduction of *t*_*1/2*_ in our experimental conditions can translate to *k* increments of more than 4 orders of magnitude (**Supplementary Fig. 19**).

**Fig. 2:**
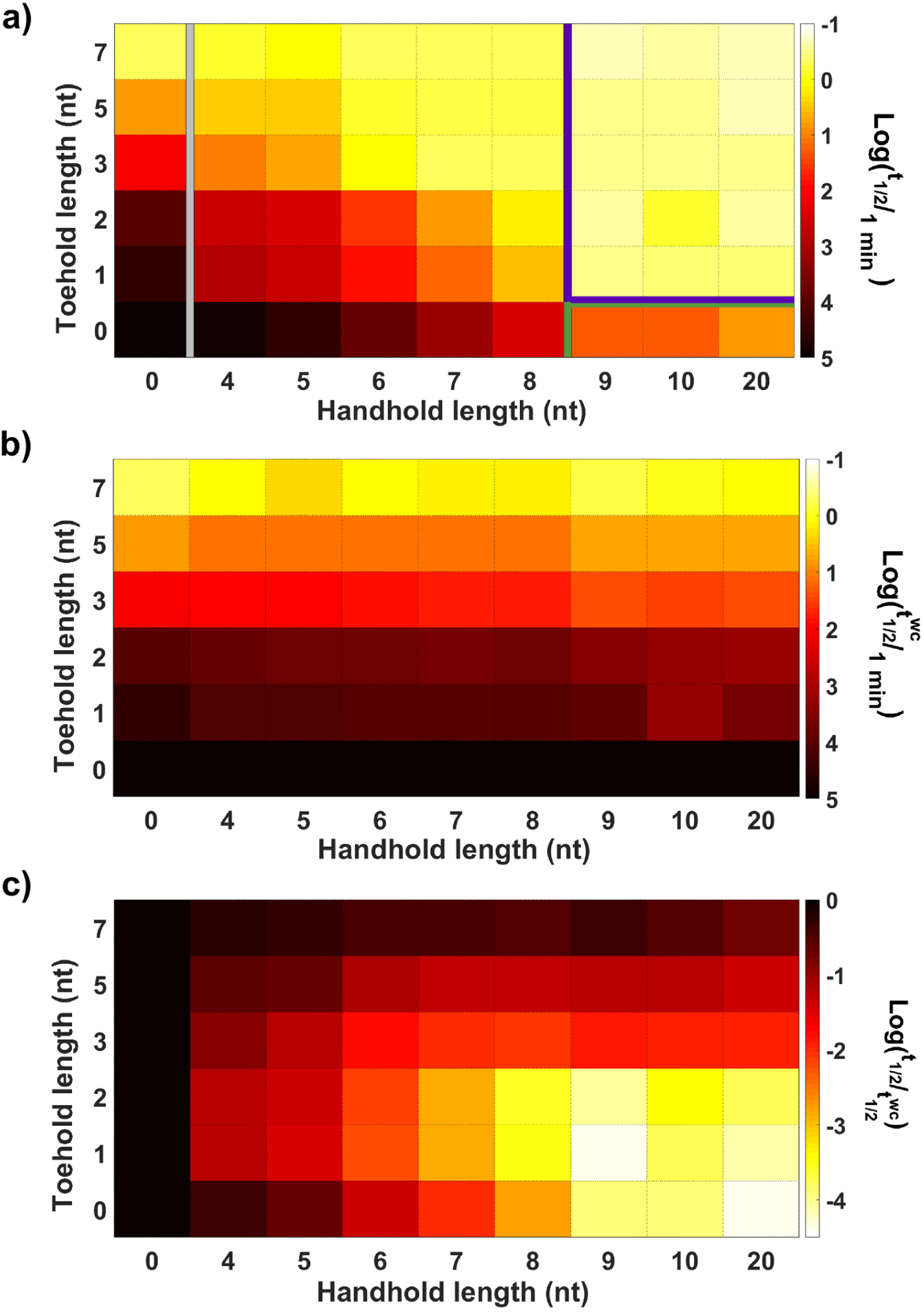
Handhold-complementarity systematically enhances strand displacement rates for short toeholds as inferred from the fits shown in Supplementary Table 15. **A) Half-lives (*t***_***1/2***_**) for HMSD**,. The analysis shows three different kinetic regimes: 2^nd^ order reactions with reversible handhold binding (from 4 to 8 nt handhold) and with irreversible handhold binding (outlined in purple); and 1^st^ order reactions where the limiting factor is the displacement once bound to the handhold (outlined in green). Reactions in the absence of handhold are outlined in grey. *t*_*1/2*_ calculated for [*I*]_0_=6 nM and [*TN*]_0_=10 nM with parameters inferred at 25°C. **B) Half-lives for *I* strands without handhold-complementarity 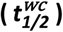.** Handhold presence in *N* produces negligible rate changes for strand displacement reactions when *I-*strands lack handhold-complementarity. **C) Comparison between *t***_***1/2***_ **and 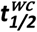.** HMSD can reduce displacement *t*_*1/2*_ up to four orders of magnitude compared to displacement triggered with *I-*strands without handhold-complementarity. This is translated as an increment of reaction rate of more than 4 order of magnitude. The reaction rate increment obtained by the addition of handhold-complementarity is sensibly higher in the regime of toeholds below 3 nt.

For HMSD to facilitate far-from-equilibrium templating, handhold binding must also be transient and release the product *IT* upon completion. To verify release, we design another reporter DNA duplex (*R*_*2*_*Q*_*2*_) to detect detachment of *IT* from *N*. Like our previous reporter, *R*_*2*_*Q*_*2*_ strands are labelled with a quenched reporter. *R*_*2*_*Q*_*2*_ has an associative toehold^**10**^ formed by two single-stranded domains brought into close proximity by a hairpin (**Fig. 3a**). As shown in **Supplementary Fig. 13**, *R*_*2*_*Q*_*2*_ only reacts at high-speed with *IT* that is free in solution. The reaction that describes the kinetics of *R*_*2*_*Q*_*2*_ is:

**Fig. 3:**
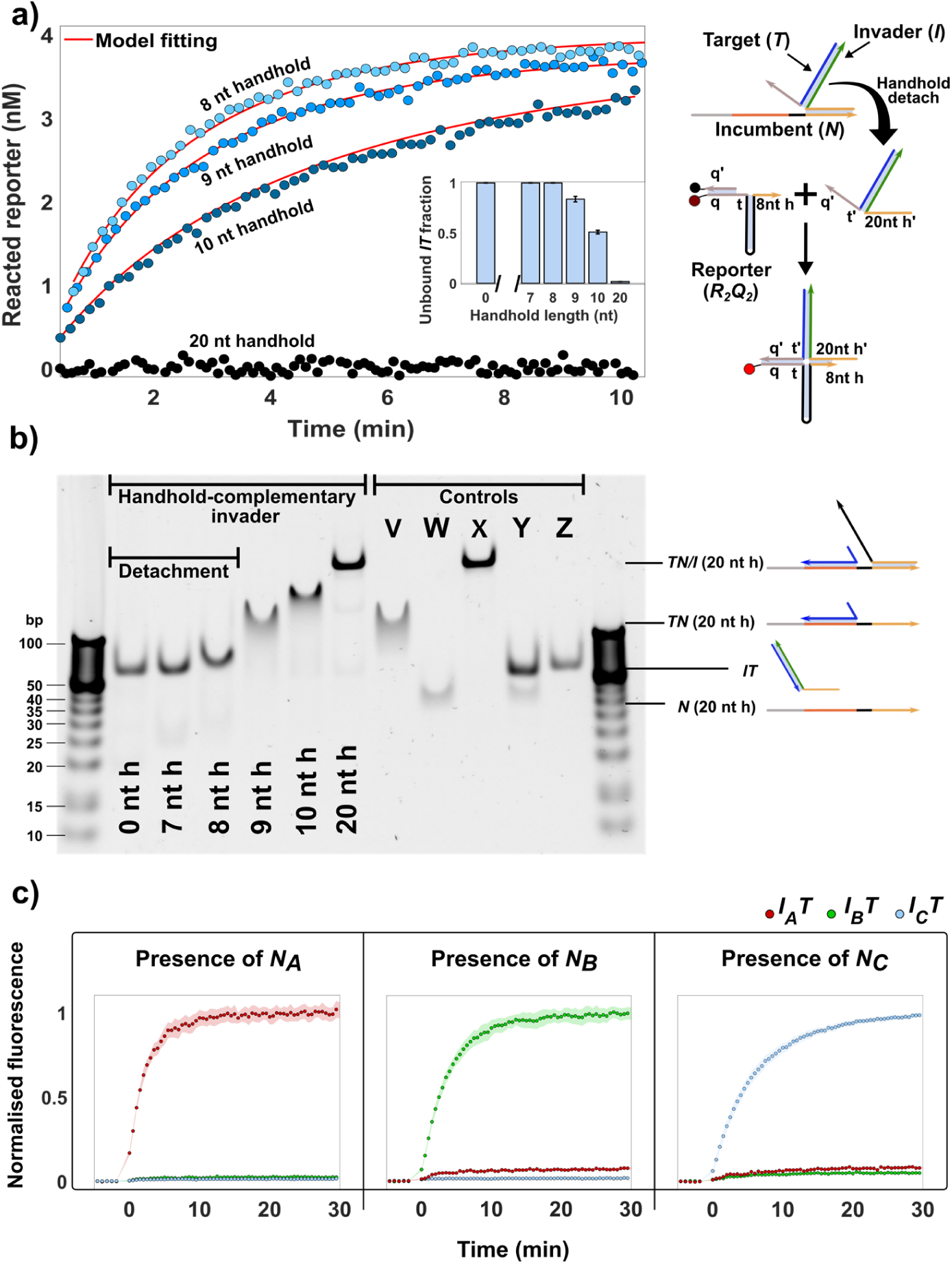
Handholds of moderate length accelerate displacement while allowing for detachment. **A) Kinetic analysis.** HMSD is triggered under the same experimental conditions as described in Fig.1 with [*I*]_0_=4 nM and a 3 nt proximal toehold. Once HMSD reaches equilibrium, reporter *R*_*2*_*Q*_*2*_ is added to react with any free *IT* by an associative toehold strand displacement. The speed of reporter triggering is negatively correlated with handhold length, saturating at 8 nt. *R*_*2*_*Q*_*2*_ kinetics are fitted as outlined in **Supplementary Note XVII** and used to infer the initially unbound fraction of *IT* product (inset). **B) Native polyacrylamide gel electrophoresis (PAGE).** 12.5 mM MgCl_2_ in 0.5x TBE at 25°C. Complete detachment after reaction is clearly identified for handholds up to 8 nt. All controls had a handhold length of 20 nt. V: *TN* duplex. W: *N-*strand. X: *TN* duplex + random strand with handhold-complementarity (results in a 3-strand complex since it cannot displace *N*). Y: *TN* + *I* without handhold-complementarity (results in detached *IT* + *N)*. Z: *IT* duplex. **C) Selective production of equally-stable complexes *I***_***A***_***T, I***_***B***_***T* and *I***_***C***_***T*.** Three reporters are used to detect the production of *I*_*x*_*T* complexes with HMSD (8 nt handhold/2 nt proximal toehold). We show that the specific production of *I*_*x*_*T* complexes is determined by the handhold sequence of the added *TN*_*x*_ complex. Standard error of the mean fluorescence (n=4) for each experiment is represented by shaded area. Conditions: [*TN*_*x*_]=10 nM, [*I*_*x*_]=25 nM for all three *I*_*x*_ strands, and 30 nM of each specific reporter, 1M NaCl in 1x TAE at 25°C.

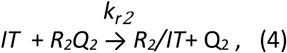

where *k*_*r2*_ is a 2^nd^ order rate constant. We obtain its value from systems with no handhold, where *IT* is fully detached ([*IT*]=[*I*]_0;_ [*N/IT*]=0). For systems with a handhold, by adding *R*_*2*_*Q*_*2*_ after HMSD reaches equilibrium, it is possible to use the reaction kinetics to extrapolate the detached amount of *IT*. We estimate the equilibrium constant:

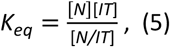

assuming that *IT* remains in a dynamic equilibrium with the *N/IT* complex, which is reasonable given that *k*_*b*_ >*k*_*r2*_. Consequently, we fit the experimental *R*_*2*_*Q*_*2*_ dynamics with the value of *K*_*eq*_ as described in *Materials and Methods.* From the fitted *K*_*eq*_, it is possible to estimate that, at experimental conditions (See *Materials and Methods*) and handhold lengths under 9 nt, over 99% of the products are in the *IT* form (**Fig. 3a)**. For handhold lengths of 9 and 10 nt, *N/IT* becomes more prevalent, as evidenced by the slowdown in reporter kinetics, and for handholds of length 20 nt, all products are sequestered, and reporter triggering is minimal. We confirm the detachment results with polyacrylamide gel electrophoresis^23^ for HMSD products with handhold lengths ranging from 0 to 20 nt. Although the gel could not be run at identical solution conditions, the results also show the substantial presence of three-stranded *N/IT* complexes only for handhold lengths over 8 nt (**Fig. 3b**).

We next harness HMSD to demonstrate the production of a far-from equilibrium ensemble of *IT* products through templating. We produce an iteration of the **Fig. 3a** experiment, but with three competing invading strands *I*_*A*_, *I*_*B*_ and *I*_*C*_, each of which is complementary to a *T* strand with a 2 nt toehold, and to different 8 nt handholds located in *N*_*A*_, *N*_*B*_ and *N*_*C*_ respectively. Free products *I*_*A*_*T, I*_*B*_*T* and *I*_*C*_*T* trigger three different reporters through an associative toehold. All three *IT* products have similar stability and their free energy is independent of the presence of other strands in solution. In **Fig. 3c** we show that the formation of any single *I*_*x*_*T* product, free in solution, is selectively templated with the appropriate handhold from a solution containing all three invaders. The bias for any one product arises without encoding that bias into the *I*_*x*_*T* binding, and so the handhold-based templating produces far-from-equilibrium complexes.

In conclusion, we have demonstrated that HMSD is not only a simple and rational method to control strand displacement kinetics, but also adds an important feature for DNA nanotechnology: templating far-from-equilibrium assemblies. Other superficially similar processes^10, 11, 17, 24^ do not involve a transient recognition interaction. HMSD paves the way to more biologically inspired DNA systems that exploit templating; the next challenge is to incorporate HMSD into a mechanism that allows a single template to repeatedly catalyse the formation of sequence-specific dimers. Eventually, HMSD has the potential to underlie engineered mechanisms for the copying of information in longer templates, as synthetic analogues of natural processes such as transcription.

## Data availability

All experimental files and the MATLAB^®^ scripts used for its analysis are available for download through the file repository Zenodo at www.zenodo.org/record/3832588.

## Author contributions

JCG and TEO conceived the project and planned the experiments. JCG performed the experiments and analysed experimental data. All authors interpreted the results and co-wrote the paper.

## Acknowledgements

This work is part of a project that has received funding from the European Research Council (ERC) under the European Union’s Horizon 2020 research and innovation programme (Grant agreement No. 851910). The authors acknowledge financial support from the Royal Society (JCG and TEO), from the Royal Academy of Engineering (RAE) via the RAE Chair in Emerging Technologies (GBS), and from the Engineering and Physical Sciences Research Council (EPSRC) (WB, grant agreement EP/P02596X/1)

Supplementary information accompanies this paper.

## DNA sequence design

DNA sequences that minimise undesired interactions during HMSD were designed with bespoke scripts using NUPACK server (http://www.nupack.org)^1^. All strands were purchased from Integrated DNA Technologies (IDT) with HPLC purification and normalized at 100 µM in LabReady^®^ buffer. All used sequences are listed by function in **Supplementary Tables 20 to 23**. (Incumbent (*N*), Target (*T*), Invader (*I*) and reporters).

## Duplex preparation

*TN* duplexes were formed by combining 200 nM of *T* with a 10% excess of *N* to ensure every *T* is annealed. For *Reporter characterisations* the *TN* anneal was prepared with a 10% excess of *T* since the strand is inert during those assays. Annealing was performed in experimental buffer (TAE 1X and 1M NaCl, pH 8.3), except for PAGE samples which were annealed in a solution of TAE 1X and 12.5 mM MgCl_2_. Strands forming reporter complexes, except *R*_*2*_*Q*_*2*_, were mixed at a concentration of 300 nM. *RQ* was annealed in experimental buffer with a 10% excess of quencher-labelled strand. *Rep*_*A*_, *Rep*_*B*_ and *Rep*_*C*_ were annealed with a 60% excess of quencher-labelled strand to minimise the crosstalk between reporter complexes. Each 100 µL of *TN* or reporter solution, except *R*_*2*_*Q*_*2*_, was annealed by heating to 95°C for 4 min and then cooled to 20°C at a rate of 1°C/min. *R*_*2*_*Q*_*2*_ reporter was annealed at a concentration of 60 nM with 10% excess of quencher-labelled strand. Each 500 µL of *R*_*2*_*Q*_*2*_ was heated up to 95°C for 5 minutes and then cooled to room temperature over 1 hour.

## Bulk fluorescence spectroscopy

Bulk fluorescence assays were carried out in a Clariostar Microplate reader (BMG LABTECH) using flat µClear bottom 96-well plates (Greiner), reading from the bottom. Kinetics were recorded, unless stated otherwise, after injecting 50 µL of the reaction trigger in 150 µL of experiment buffer containing the reactant species and the reporter complex (pump speed: 430 µL/s). The final mixture was shaken for 3 seconds (double-orbital, at 400 rpm). Injected and reacting volumes were previously preheated to the experiment temperature. The samples were contained in Eppendorf^®^ Lobind tubes and the plate reader’s injector system passivated by incubating with BSA 5% during 30 minutes to maximise concentration reproducibility during the assays^2^.

*RQ* reporter was labelled with Cy3 and FQ IowaBlack™ quencher (Excitation: 530/20 nm; Emission: 580/30 nm). *R*_*2*_*Q*_*2*_ reporter was labelled with Cy5 and RQ IowaBlack™ quencher (Excitation: 610/30 nm; Emission: 675/50 nm). Signal was averaged for 20 flashes per data point. Experiments with durations over 12 hours were averaged for 100 flashes in spiral area scan per datapoint. *Reporter characterisation* assays were measured individually with 1 flash/datapoint due to the fast rate of the reaction. Every system was assayed with at least three different reaction trigger concentrations.

## Fluorescence calibrations

Fluorescence calibrations were made for the reporter DNA duplexes used during characterisation assays (*RQ*/Cy3 and *R*_*2*_*Q*_*2*_/Cy5). Calibrations aimed to obtain the units of fluorescence produced per nM of fluorophore-labelled strand to allow quantification of reacted reporter during the assays. Calibration curves ranged from 1 to 15 nM in 200 µL and were made by triplicate from stock solutions at 100 µM. One replica included both fluorophores (Cy3 and Cy5) to rule out bleed-through fluorescence. Detailed description of the protocol and results are discussed in **Supplementary Notes IV and V and Supplementary Figures 6 to 9**.

## General protocol for characterisation experiments

HMSD systems with a combination of 9 different handhold lengths (0, 4, 5, 6, 7, 8, 9, 10 and 20 nt) and 6 different proximal toehold (0, 1, 2, 3, 5, and 7 nt) were tested at 25°C. A smaller set of systems were also characterised at 37°C (**Supplementary Table 17**). The effect of a spacer sequence located after the handhold of *N* was also tested. Spacer lengths of 0, 1, 2, 3, 5 and 8 nt were included for systems with handhold lengths of 0, 5 and 20 nt and proximal toehold lengths of 0, 2 and 7 nt. The addition of a spacer sequence did not have a clear effect on HSMD except for systems with 5 nt handholds and proximal toeholds of 0 or 2 nt, where a spacer length of 2 nt was found optimal (**Supplementary Table 16**). Based on this finding, all systems reported in the main text contain a 2 nt spacer in *N.*

Experimental protocols for the characterisation were adapted from the method employed by Srinivas et al.^3^. Each experiment consisted of the system kinetics and a set of complementary measurements. These complementary measurements quantified the fluorescence baseline and estimated the concentration of each species in the system from the fluorescence signal measured after sequentially triggering the reaction of all the species (**Supplementary Figures 1 and 2**). Intended species concentrations during the experiments, unless specified otherwise, were [*RQ*]_0_=15 nM, [*TN*]_0_=10 nM and [*I*]_0_=[4-8] nM, at a reference reaction volume of 200 µL. The purpose of considering a range of concentrations for the *I*-strand was to test the robustness of the reaction kinetics for their 2^nd^ order and pseudo-1^st^ order approximations, whilst simultaneously providing an estimate of the statistical error in the inferred reaction kinetics. A detailed description of the quantification method and the measurements during each type of characterisation assay is described in **Supplementary Notes I to III**.

## General fitting procedure

All fittings were performed with MATLAB^®^ R2019a Optimization Toolbox. In addition to the relevant rate constants, each fitting also allowed a variation of the fitted curve’s reaction initiation time (*t*_*0*_). This correction amended the initial leak of fluorescence signal produced by defective duplexes that react instantaneously with the reporter. *t*_*0*_ fitting became especially relevant for slow reactions where the initial increment of fluorescence could be equivalent to that after several hours of reaction. On the other hand, for reactions that consumed at least 90% of its reaction trigger, the estimated trigger concentration was also fitted to correct any discrepancy with the fluorescence signal of the equilibrium plateau. Detailed description of the fitting procedures is available in **Supplementary Notes XI to XVII**.

## Reporter characterisation

The value of *k*_*r*_ was obtained experimentally for every species of *N*. Experiments consisted of a solution of *RQ* triggered by a solution of *N/IT* ([*N/IT*]_0_=[4-8] nM). The injected *N/IT* solution was obtained by mixing *TN* with a proximal toehold of 7 nt with a 20% excess of *I*_*hh*_. Each individual trace was fitted to the analytical solution of the ODE describing **equation 2**:

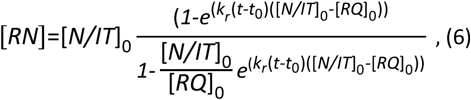

With *RN* the fluorescent species. Modifications in the size and sequence of the *N* strands produced moderate variation in *k*_*r*_. This variation in *k*_*r*_ was particularly relevant during the fitting of HMSD systems with higher rates. Therefore, subsequent fits used the *k*_*r*_ obtained for the *N* present in each fitted system to minimise the fitting error. All fitted *k*_*r*_ are collected in **Supplementary Tables 9 and 10**.

## Toehold-mediated reaction characterisation

The experimental value of *k*_*t*_ was obtained for most of the combinations of handhold and toehold lengths assayed. Otherwise, *k*_*t*_ was inferred from the mean of the *k*_*t*_ obtained for systems with the same toehold length. Experiments consisted of a solution of *RQ* and *TN* triggered by *I*_*wc*_. Each individual trace was fitted to a reduced ODE model describing **equation 3** and a TMSD 2^nd^ order reaction:

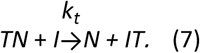

The *k*_*t*_ obtained for each system was used in subsequent fits to account for any unpredicted effect of the handhold length on *k*_*t*_. All fitted *k*_*t*_ are collected in **Supplementary Tables 15 to 17**.

## HMSD rate characterisations

A solution of *RQ* and *TN* was triggered by *I*_*hc*_. For each system, their traces were fitted simultaneously to the ODE model implied by **equations 1, 2, 3, 7** and the additional handhold binding/unbinding reactions:

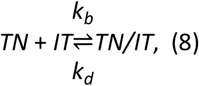

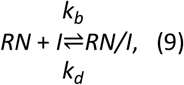

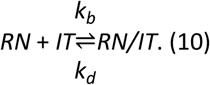

The species *TN/IT* resultant from **equation 8** is unable to undergo displacement via the handhold-bound IT complex. However, invasion of the free proximal toehold in the *TN* duplex still occurs with the rate *k*_*t*_ :

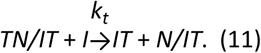

The mass-action-based ODE model of the full system (**equations 1**,**2**,**3**,**7**,**8**,**9**,**10 and 11**) is included in **Supplementary Note XI**. During the fitting, parameters *k*_*r*_ and *k*_*t*_ were fixed at the values obtained as described in *Reporter characterisation* and *Toehold-mediated reaction characterisation*, respectively. *k*_*b*_ was constrained to a certain range, determined by fittings done in systems confidently located in the 2^nd^ order regime with irreversible handhold binding (handhold=20 nt; toehold>1 nt).

For the proposed reaction mechanism, most systems operated in the 2^nd^ order regime with reversible handhold-binding, where the reaction rate was defined by relative values of *k*_*b*_, *k*_*d*_ and *k*_*m*_ (**equation 4**). This fact precluded the fitting algorithm from obtaining comparable absolute values for the rate constants across all tested systems. Constraining *k*_*b*_ to a fixed range eased this problem while still producing low error fittings for all tested systems. However, compensatory errors in the ratio 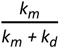 could mitigate for an inaccurate *k*_*b*_. Therefore, the effective bimolecular rate constant (**equation 4**) is considered the most reliable descriptor of system kinetics in the 2^nd^ order regime. For all tested systems, the mean values of its fitted parameters and their effective rates are collected in **Supplementary Tables 15 to 17**.

## HMSD half-life (*t*_*1/2*_) calculation

Using the rate constants obtained from *HMSD rate characterisations*, new model traces were generated for each system using the complete ODE reaction model (**Supplementary Note XI**). Traces were generated for initial conditions [*I*]_0_=6 nM, [*TN*]_0_=10 nM and [*RQ*]_0_=1 µM. The time at which 3 nM of [*I*] had reacted was considered to be the *t*_*1/2*_ of that system. The high concentration of simulated [*RQ*]_0_ made the reporter reaction effectively instantaneous, removing variability produced by *k*_*r*_. Obviating the reporter reaction also allowed calculation of the analytical expression for 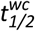 as:

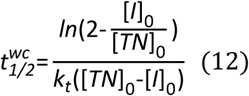

All calculated *t*_*1/2*_ and 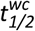 are collected in **Supplementary Tables 15 to 17**.

## Detachment assays

*I*_*hc*_ was added to a solution containing *RQ* and *TN* to produce a final mixture of *IT* and *N/IT* ([*IT*]+[*N/IT*]=[*I*]_0_=[4-8] nM). After the system reached equilibrium, 50 µL of *R*_*2*_*Q*_*2*_ was injected into the experimental solution ([*R*_*2*_*Q*_*2*_]_0_= 15 nM). During these assays, the *T* strand contained an additional domain in 3’ to trigger reporter *R*_*2*_*Q*_*2*_.The kinetics of *IT* triggering *R*_*2*_*Q*_*2*_ were measured for 10 minutes. The initial concentrations for all the strands present in the solution (*I,N* and *T*) were inferred from the signal of reporter *RQ*. The results of the controls without reporter *RQ*, shown in **Supplementary Table 19**, confirmed that the presence of *RQ* did not alter the equilibrium between *IT* and *N/IT*. For controls in the absence of *RQ*, concentrations were inferred from reporter *R*_*2*_*Q*_*2*_ signal.

To obtain the value of *k*_*r2*_, the traces of a system without handhold were fitted to the analytical solution of the ODE describing **equation 4**, analogously to *RQ* (See *Reporter characterisation* and **Supplementary Figure 20**). For systems with a handhold, each trace was fitted to the ODE describing **equation 4** with concentrations constrained by **equation 5**, to estimate the value of *K*_*eq*_. From the obtained *K*_*eq*_ the percentage of detached species was calculated for concentrations [*I*]_0_;[*N*]_0_=10nM. All fitted *K*_*eq*_ are collected in **Supplementary Table 18**.

## Polyacrylamide gel electrophoresis

The PAGE gel for detachment analysis (**Figure 4b**) was cast with 12% 37.5:1 acrylamide:bisacrylamide (Millipore^®^) and 12.5 mM MgCl_2_ (effective concentration 12 mM MgCl_2_) in TBE 0.5x and used in the same day. Samples were annealed in the presence of MgCl_2_ and saturated afterwards with *I*_*hc*_ while incubating at 25°C (as described in *Duplexes annealing*). Controls and O’range ruler 5bp ladder (Thermo Scientific™) were incubated simultaneously. Gel was pre-run for 10 min at 120 V and after that, the samples were loaded and run for 2 hours at the same voltage. 15 µL of each sample were mixed with 3 µL of orange DNA loading dye (6x) (Thermo Scientific™) and 17 µL of the mixture were loaded in each well. The gel ran in a Mini-Protean^®^ electrophoresis chamber (Bio-Rad) keeping the running buffer (0.5xTBE and 12.5 mM MgCl_2_) at 25 °C using a water bath. Gels were stained for 30 min with a GelRed^®^ 3x solution (Biotium.inc) and visualised in a FluorChem^®^ FC2 imager (Alpha Innotech). Excitation: 300 nm. Bandpass filter: 620/40 nm.

## Selective production of complexes far-from-equilibrium

The experimental system contained three reporter complexes: Rep_A_ (Identical to *R*_*2*_*Q*_*2*_) and two analogous reporters: Rep_B_ and Rep_C_. Each reporter differed in its distal toehold sequence, which enabled them to specifically recognise one of the product complexes *I*_*A*_*T, I*_*B*_*T*, or *I*_*C*_*T* when free in solution (**Supplementary Note X**). Rep_B_ was labelled with Rhodamine Red™-X and Black Hole Quencher^®^ (Excitation: 550/20 nm; Emission: 605/40nm). Rep_c_ was labelled with Alexa Fluor^®^ 488 and FQ IowaBlack™ quencher (Excitation: 488/14 nm; Emission: 535/40 nm).

Each experimental solution consisted of 190 µL of experimental buffer at 25°C, containing the three invader species with different handhold sequences (*I*_*A*_, *I*_*B*_, and *I*_*C*_) and the three reporters (Rep_A_, Rep_B_ and Rep_C_). The reaction kinetics were triggered by adding 10 µL of *TN*_*x*_ complex solution with a specific handhold (*TN*_*A*,_ *TN*_*B*_ or *TN*_*C*_). Initial concentrations were 25 nM of each species of *I*_*x*_, 30 nM of each reporter and 10 nM of *TN*_*x*_ complex. The traces for each different *TN*_*x*_ were assayed 4 times and the fluorescence of each reporter normalised with its mean maximum fluorescence during the assays.

## References

1. Chen, Y.J. et al. Programmable chemical controllers made from DNA. Nat. Nanotechnol. 8, 755–762 (2013).

2. Xin, L., Zhou, C., Duan, X. & Liu, N. A rotary plasmonic nanoclock. Nat. Commun. 10, 5394 (2019).

3. Hirose, Y. & Manley, J.L. RNA polymerase II and the integration of nuclear events. Genes Dev. 14, 1415–1429 (2000).

4. Rodnina, M.V., Gromadski, K.B., Kothe, U. & Wieden, H.J. Recognition and selection of tRNA in translation. FEBS Lett. 579, 938–942 (2005).

5. Yurke, B., Turberfield, A.J., Mills, A.P., Simmel, F.C. & Neumann, J.L. A DNA-fuelled molecular machine made of DNA. Nature 406, 605–608 (2000).

6. Srinivas, N., Parkin, J., Seelig, G., Winfree, E. & Soloveichik, D. Enzyme-free nucleic acid dynamical systems. Science 358 (2017).

7. Green, A.A. et al. Complex cellular logic computation using ribocomputing devices. Nature 548, 117–121 (2017).

8. Zhang, D.Y. & Winfree, E. Control of DNA Strand Displacement Kinetics Using Toehold Exchange. J. Am. Chem. Soc. 131, 17303–17314 (2009).

9. Srinivas, N. et al. On the biophysics and kinetics of toehold-mediated DNA strand displacement. Nucleic Acids Res. 41, 10641–10658 (2013).

10. Chen, X. Expanding the rule set of DNA circuitry with associative toehold activation. J. Am. Chem. Soc. 134, 263–271 (2012).

11. Genot, A.J., Zhang, D.Y., Bath, J. & Turberfield, A.J. Remote toehold: a mechanism for flexible control of DNA hybridization kinetics. J. Am. Chem. Soc. 133, 2177–2182 (2011).

12. Crick, F. Central Dogma of Molecular Biology. Nature 227, 561–563 (1970).

13. Tjivikua, T., Ballester, P. & Rebek, J. Self-replicating system. J. Am. Chem. Soc. 112, 1249–1250 (1990).

14. Ouldridge, T.E. & Rein Ten Wolde, P. Fundamental Costs in the Production and Destruction of Persistent Polymer Copies. Phys. Rev. Lett. 118, 158103 (2017).

15. Poulton, J.M., Ten Wolde, P.R. & Ouldridge, T.E. Nonequilibrium correlations in minimal dynamical models of polymer copying. Proc. Natl. Acad. Sci. U. S. A. 116, 1946–1951 (2019).

16. Sartori, P. & Leibler, S. Lessons from equilibrium statistical physics regarding the assembly of protein complexes. Proceedings of the National Academy of Sciences 117, 114–120 (2020).

17. Lanzmich, S.A. Replication in Early Evolution. (Ludwig-Maximilians-Universität, 2016).

18. Schulman, R., Yurke, B. & Winfree, E. Robust self-replication of combinatorial information via crystal growth and scission. Proc. Natl. Acad. Sci. U. S. A. 109, 6405–6410 (2012).

19. O’Flaherty, D.K., Zhou, L.J. & Szostak, J.W. Nonenzymatic Template-Directed Synthesis of Mixed-Sequence 3 ‘-NP-DNA up to 25 Nucleotides Long Inside Model Protocells. J. Am. Chem. Soc. 141, 10481–10488 (2019).

20. Kreysing, M., Keil, L., Lanzmich, S. & Braun, D. Heat flux across an open pore enables the continuous replication and selection of oligonucleotides towards increasing length. Nat. Chem. 7, 203–208 (2015).

21. Moreira, B.G., You, Y. & Owczarzy, R. Cy3 and Cy5 dyes attached to oligonucleotide terminus stabilize DNA duplexes: predictive thermodynamic model. Biophys. Chem. 198, 36–44 (2015).

22. SantaLucia, J., Jr. & Hicks, D. The thermodynamics of DNA structural motifs. Annu. Rev. Biophys. Biomol. Struct. 33, 415–440 (2004).

23. Stellwagen, N.C. Apparent pore size of polyacrylamide gels: Comparison of gels cast and run in Tris-acetate-EDTA and Tris-borate-EDTA buffers. Electrophoresis 19, 1542–1547 (1998).

24. Genot, A.J., Bath, J. & Turberfield, A.J. Combinatorial Displacement of DNA Strands: Application to Matrix Multiplication and Weighted Sums. Angew. Chem. Int. Ed. 52, 1189–1192 (2013).

## References

1. Wolfe, B.R., Porubsky, N.J., Zadeh, J.N., Dirks, R.M. & Pierce, N.A. Constrained Multistate Sequence Design for Nucleic Acid Reaction Pathway Engineering. J. Am. Chem. Soc. 139, 3134–3144 (2017).

2. Kanoatov, M. & Krylov, S.N. DNA Adsorption to the Reservoir Walls Causing Irreproducibility in Studies of Protein–DNA Interactions by Methods of Kinetic Capillary Electrophoresis. Anal. Chem. 83, 8041–8045 (2011).

3. Srinivas, N., Parkin, J., Seelig, G., Winfree, E. & Soloveiehile, D. Enzyme-free nucleic acid dynamical systems. Science 358 (2017).

