## Supplementary Material for "Handhold-mediated strand displacement: a nucleic acid-based mechanism for generating far-from-equilibrium assemblies through templated reactions" (from another version of this preprint: version 1, 2020-05-22)

#### Supplementary Notes

#### Supplementary Figures

### Supplementary Tables

Supplementary Table 1 | Detailed experimental procedure for reporter *RQ* characterisation. 5

Supplementary Table 2 | Detailed experimental procedure for toehold/handhold-mediated strand displacement characterisation. 5

Supplementary Table 3 | Detailed experimental procedure for the kinetic analysis of handhold detachment. 8

Supplementary Table 4 | Detailed experimental procedure to produce far-from-equilibrium complexes 11

Supplementary Table 5 | Steps of the reaction control for *RQ* 19

Supplementary Table 6 | Steps of the control for *I* without *T*-complementarity. 19

Supplementary Table 7 | Steps of the $R_2Q_2$ control 21

Supplementary Table 8 | Steps of the selectivity controls for Rep complexes 23

Supplementary Table 9 | Fitted $k_r$ at 25˚C 30

Supplementary Table 10 | Fitted $k_r$ at 37˚C 31

Supplementary Table 11 | $k_b$ fitting at 25 ˚C 33

Supplementary Table 12 | $k_{bind}$ fitting at 37 ˚C 33

Supplementary Table 13 | Estimated $\Delta G_{hb}$ and $k_d$ for each handhold at 25 ˚C 34

Supplementary Table 14 | Estimated $\Delta G_{hb}$ and $k_d$ for each handhold at 37 ˚C 35

Supplementary Table 15 | Handhold Characterisation at 25 ˚C. 36

Supplementary Table 16 | Spacer Characterisation at 25 ˚C. 38

Supplementary Table 17 | Handhold Characterisation at 37 ˚C. 39

Supplementary Table 18 | Fitting results from the kinetic analysis of handhold detachment 42

Supplementary Table 19 | Fitting results from the kinetic analysis of handhold detachment in the absence of reporter *RQ*. 43

Supplementary Table 20 |Incumbent strands (*N*) 47

Supplementary Table 21 |Target strands (*T*) 48

Supplementary Table 22 |Invader strands (*I*) 48

Supplementary Table 23 |Reporter strands 48

#### Supplementary Note I: General procedure for measuring reaction kinetics

All experiments contained at least one negative control (buffer only) and one positive control per reporter used, which allowed correction of changes in fluorescence by temperature and reaction volume. Positive controls contained a variable concentration of the reporter's fluorophore-labelled strand in experimental buffer (TAE 1x,1M NaCl) and negative controls were composed of just experimental buffer. The volume contained in the controls was changed consistently with the volume of the main experiments. The function of the controls during data analysis is further described in **Supplementary Note VI**.

The concentration for each strand species was inferred from measuring steady-state fluorescence due to the uncertainties introduced by pipetting errors. All measurements consisted of at least 10 data points. The number of measurements per experiment depends on the class of characterisation experiment, however, they all followed a similar structure that helped to quantify all the species in the experimental solution as outlined below. Specific procedures for each type of experiment are collected in **Supplementary Tables 1 and 2**. Examples of their fluorescence traces are shown in **Supplementary Figures 1 and 2.** For convenience, in all Supplementary Tables and Figures we report the quantity of each species in the reaction volume at any time as the concentration that would be present if the same amount was solvated in a reaction volume of 200 µL.

Measurement 1. Fluorescence baseline produced by residual fluorescence of *RQ*:
The experiments started by mixing 10 µL of a standard solution of *RQ* (300 nM) with experimental buffer. A measurement was made after 30 mins of incubation at the experimental temperature, when the fluorescence value of the positive controls was stabilised.

Measurement 2. Estimation of $[N]_{excess}$:
10 µL of a standard solution of *TN* (200 nM) were added to the solution, except during *Reporter characterisations*. After adding *TN*, the excess of *N* resultant from the annealing of *TN* ($[N]_{\text{excess}}$) immediately reacted with *RQ,* producing an increment of the fluorescence baseline (**See Supplementary Note XI**). The new baseline was recorded and used to infer $[N]_{\text{excess}}$.

Measurement 3. Experimental kinetics:
The next measurement tracked the kinetics after injecting 50 µL of a solution with the reaction trigger (*I* or *N/IT*) for a final reaction volume of 200 µL. Final intended reagent concentrations at 200 µL were $[RQ]_0$=15 nM, $[N]_0$=10 nM and $[I]_0$ ranging from 4 to 8 nM, unless specified otherwise. The system kinetics were measured, when possible, until the reaction reached steady state. For kinetics that took more than 2 hours to reach steady-state, several systems, with their respective controls, were ran in parallel in a single 96-well plate. The kinetics of the experiments in the plate were always measured for at least 100 hours. After it, the plate was sealed, and incubated, preserved from light, for 1 month at the experimental temperature to let the reaction reach steady state.

Measurement 4. Estimation of reaction trigger concentration from the steady-state fluorescence:
After the reaction reached equilibrium, a measurement recorded the fluorescence steady state. The total amount of reaction trigger added was inferred from the level of this steady state. In some cases, the trajectories did not reach steady state after 1 month (proximal toehold=0 nt; short handholds). For those cases, the concentration of trigger was inferred from the mean concentration of all the experiments that reached steady state in the same plate. All experiments in a plate were triggered with the same aliquot of reaction trigger, so significant variability was not expected between experiments. Experiments that reached at least 80% of the value of the highest concentration of trigger in that same plate were considered to be in steady state.

Measurement 5. Saturation and quantification of the rest of the strands:

Finally, a series of saturating additions to the experimental solution were performed. High concentrations of single strands were added to trigger unreacted substrates and infer their actual initial concentrations from the resultant change in fluorescence. For example, to quantify $[TN]_0$, all the *TN* in the solution was triggered with 10 µL of a saturating solution of $I_{hc}$ (Measurement 5 i)). $[RQ]_0$ was estimated in a similar fashion with a saturating solution of *N* (Measurement 5 ii)).

**Supplementary Table 1| Detailed experimental procedure for reporter *RQ* characterisation.**

Due to the high rate of the reaction, the four iterations of each experiment were triggered and measured individually. The triggering species was a solution of *N/IT.* Since no *TN* was added, no $[N]_{excess}$ existed, and no measurement 2 was taken.

| Steps | Addition | Purpose | Expected concentrations in 200 µL (nM) | Volume (µL) |
|---|---|---|---|---|
| **Measurement 1** | 10 µL *RQ* (300 nM)<br>140 µL Buffer | Estimate residual fluorescence baseline | *RQ*:15 | 150 |
| **Measurement 3** | [45-20] µL *N/IT* (40 nM)<br>+ [5-30] µL Buffer | Experimental kinetics | Evolve with time | 200 |
| **Measurement 4** | - | Estimate $[N]_0$ | *RQ*:[6-11]; *RN*:[9-4]; *IT*:[9-4]; | 200 |
| **Measurement 5 ii)** | 10 µL *N* (400 nM) | Estimate $[RQ]_0$ | *RN*:15; *IT*:[9-4]; *N*:[14-9] | 210 |

**Supplementary Table 2| Detailed experimental procedure for toehold/handhold-mediated strand displacement characterisation.**

The four trajectories of each reaction system and their controls were triggered by mechanical injection and measured simultaneously. To characterise the handhold-mediated strand displacement, the reaction was triggered with $I_{hc}$. To characterise the toehold rate, $I_{wc}$ was used. The species $I_{wc}$ have, instead of the handhold-complementary domain, a random sequence of the same length.

| Steps | Addition | Purpose | Expected concentrations in 200 µL (nM) | Volume (µL) |
|---|---|---|---|---|
| **Measurement 1** | 10 µL *RQ* (300 nM)<br>130 µL Buffer | Estimate residual fluorescence baseline | *RQ*:15 | 140 |
| **Measurement 2** | 10 µL *TN* (200 nM) +<br>*N* ~20 nM | Estimate $[N]_{excess}$ | *RQ*:14; *TN*:10; *RN*:~1; | 150 |
| **Measurement 3** | [45-20] µL *I* (40 nM) +<br>[5-30] µL Buffer | System kinetics | Evolve with time | 200 |
| **Measurement 4** | - | Estimate $[I]_0+[N]_{excess}$ | *RQ*:[5-10]; *TN*:[1-6]; *RN*:[10-5]; *IT*:[9-4] | 200 |
| **Measurement 5 i)** | 10 µL $I_{hc}$ (400 nM) | Estimate $[N]_0$ | *RQ*:4; *RN*:11; *IT*:10; *I*:[19-14] | 210 |
| **Measurement 5 ii)** | 10 µL *N* (400 nM) | Estimate $[RQ]_0$ | *RN*:15; *IT*:10; *I*:[19-14]; *N*:16 | 220 |

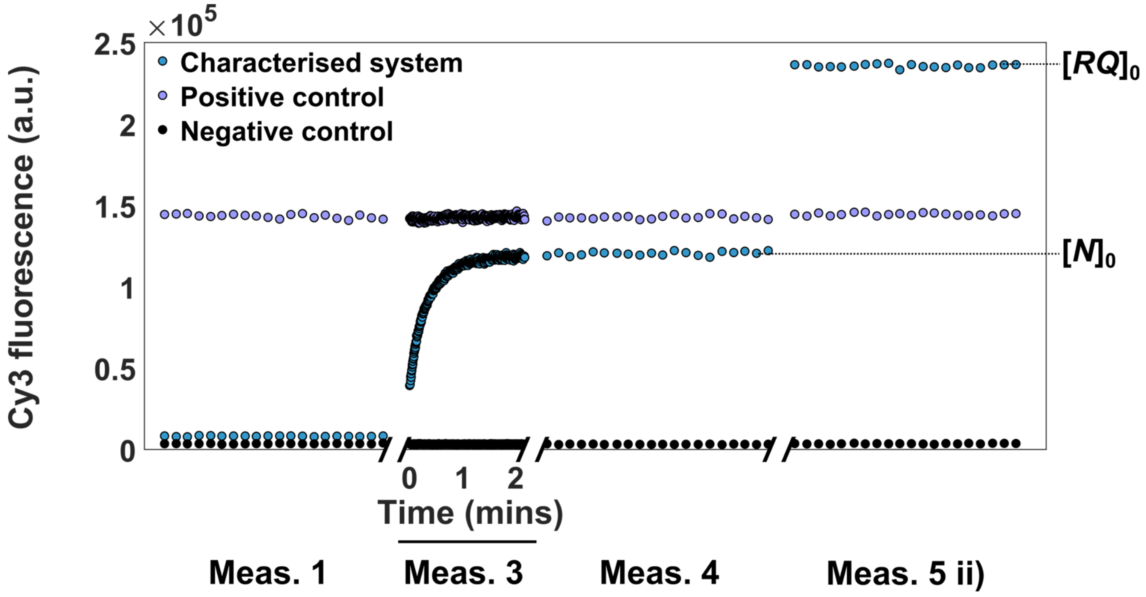

**Supplementary Figure 1| Example of reporter complex *RQ* reporter characterisation.**
Raw data from the triggering reaction of *RQ* with an *N/IT* complex with 5 nt handhold, 8 nt spacer and 7 nt toehold at 25˚C (5/8/7_25). Measurement 1 contains the baseline of the experiment produced by 15 nM of quenched *RQ* . In measurement 3, a variable amount of trigger *N/IT* was added, between 9 and 4 nM (8 nM in this case). After the reaction finished, the total amount of trigger *N/IT* was estimated from measurement 4. After that, 20 nM of *N* was added to trigger the rest of *RQ* and quantify it in measurement 5 ii). All concentrations are defined for a reaction volume of 200 µL. Positive control consisted of more than 10 nM of the Cy3-labelled reporter strand. Negative control consisted of only reaction buffer.

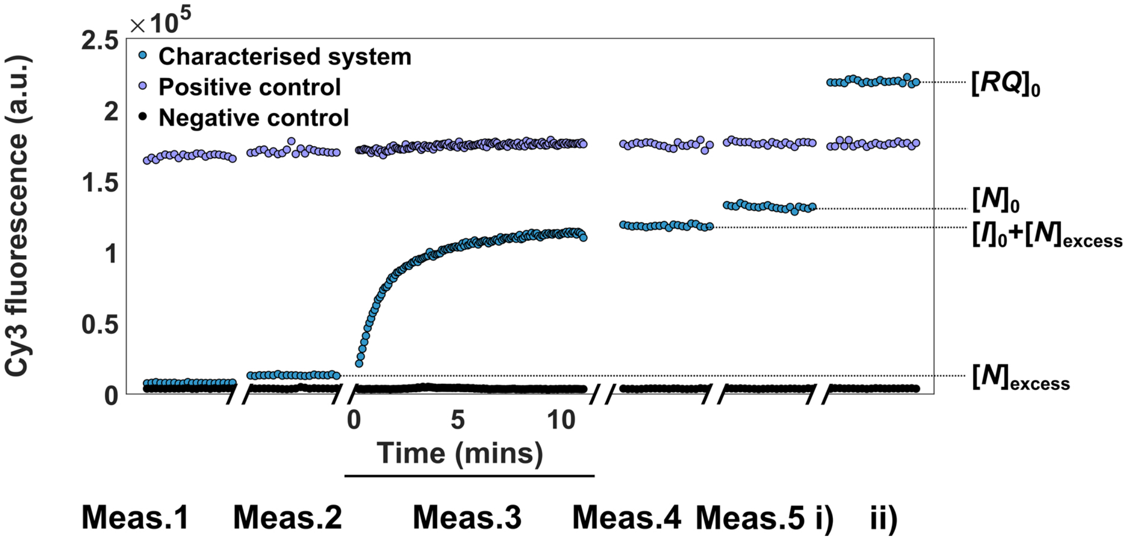

**Supplementary Figure 2| Example of a handhold-mediated strand displacement characterisation experiment.**

Raw data from the reaction system with 7 nt handhold, 2 nt spacer and 5 nt toehold at 25˚C (7/2/5_25) triggered with $I_{hc}$. Measurement 1 contains the baseline of the experiment produced by 15 nM of quenched *RQ*. Then, 10 nM of *TN* was added. $[N]_{excess}$ was inferred from the annealing increase the baseline, which was recorded in measurement 2. In measurement 3, a variable amount of trigger $I_{hc}$ was added, between 9 and 4 nM (8 nM in this case). After the reaction finished, measurement 4 was used to estimate the total amount of trigger $I_{hc}$. Measurement 5 i) saturated the solution with 20 nM of $I_{hc}$ to estimate the total amount of *N* in the solution. After that 20 nM of *N* was added to trigger the rest of *RQ* and quantify it in measurement 5 ii). All concentrations are defined for a reaction volume of 200 µL. Positive control consisted of more than 10 nM of the Cy3-labelled reporter strand. Negative control consisted of only reaction buffer.

#### Supplementary Note II: Experimental procedure for the kinetic analysis of handhold detachment

The design of detachment experiments had to be modified to minimise $R_2Q_2$ leak reactions (**Supplementary Figure 3**). The triggering mechanism for the reporter $R_2Q_2$ is composed of an associative toehold with an 8-nt distal toehold and a 2-nt proximal toehold. The distal toehold is complementary to the handhold-complementary domain of $I_{hc}$ and is unable to bind $R_2Q_2$ until $IT$ detaches. However, the proximal toehold is complementary with a 3' additional domain in $T$. This 2-nt toehold-mediated displacement reaction can always take place when $R_2Q_2$ and $T$ are together in the solution.

To reduce the time that $R_2Q_2$ and the $T$ strand were together, $R_2Q_2$ was added as the trigger of the reaction. Steps of the experiments and an example of the fluorescence traces are show in **Supplementary Table 3** and **Supplementary Figure 4.** The modification required the addition of a new control to the experiment. In addition to the negative control and the positive controls for red and green fluorescence, a 4$^{th}$ control was added to quantify the residual fluorescence resultant after quenching of $R_2Q_2$. This "quenching control" was composed of 15 nM of the reporter complex $R_2Q_2$ from the same aliquot used in the actual experiments. This control was eventually saturated with $IT$ complex to measure the total fluorescence and calculate the efficiency of quenching for $R_2Q_2$, which was used to calculate the fluorescence baseline for the experimental trajectories.

Control without *RQ* reporter complex:

The signal of reporter $RQ$ was used to infer the concentrations of all the species in the solution. The green signal shows less noise and it also allows the estimation of $[N]_{\text{excess}}$, which is relevant for the calculation of $K_{eq}$ (eq.15). To demonstrate that $RQ$ did not modify $K_{eq}$, a subset of the reactions (7,8,9 and 10 nt handhold) was repeated without including $RQ$ in the solution. In this case, the concentration of each species was inferred only from the fluorescence signal produced by $R_2Q_2$, so the quantification was more inexact and species like $N_{excess}$ could not be quantified (it was assumed to be 0). Results from this control, collected in **Supplementary Table 19**, showed that rate saturation still occurred for handholds shorter than 9 nt. The obtained $K_{eq}$ slightly differed, but that effect could be attributed to the limitations during the quantification of the assay species.

**Supplementary Table 3| Detailed experimental procedure for the kinetic analysis of handhold detachment.**

The four trajectories of each reaction system and their controls were triggered by mechanical injection and measured simultaneously. All the tested $TN$ contained 3 nt proximal toeholds and a 2 nt spacer.

| Conditions | Addition | Purpose | Expected concentrations in 200 µL (nM) | Volume (µL) |
|---|---|---|---|---|
| **Measurement 1** | 10 µL $RQ$ (300 nM)<br>130 µL Buffer | Estimate residual fluorescence baseline | $RQ$:15 | 90 |
| **Measurement 2** | 10 µL $TN$ (200 nM) +<br>$N$ ~20 Nm | Estimate $[N]_{\text{excess}}$ | $RQ$:14; $TN$:10; $RN$:~1; | 100 |
| **Measurement 3** | [45-20] µL $I_{hc}$ (40 nM) +<br>[5-30] µL Buffer | Trigger $TN$ and estimate $[I]_0+[N]_{\text{excess}}$ | $RQ$:[5-10]; $TN$:[1-6]; $RN$:[10-5]; $IT$:[9-4] | 150 |
| **Measurement 4** | 50 µL $R_2Q_2$ (60 nM). Reporter is also added to the quenched control. | $R_2Q_2$ kinetics | Evolve with time | 200 |
| **Measurement 5** | - | Estimate $[IT]^{\text{Total}}$ | $RQ$:[5-10]; $TN$:[1-6]; $RN$:[10-5];<br>$R_2IT$:[9-4]; $R_2Q_2$:[6-11] | 200 |
| **Measurement 6** | 10 µL $I_{hc}$ (400 nM) | Estimate $[N]_0$ | $RQ$:4; $RN$:11; $R_2IT$:10; $R_2Q_2$:5; $I$:[19-14] | 210 |
| **Measurement 7** | 10 µL $T$ (400 nM) | Estimate $[R_2Q_2]_0$ | $RQ$:4; $RN$:11; $IT$~[10]; $R_2IT$:15 | 220 |

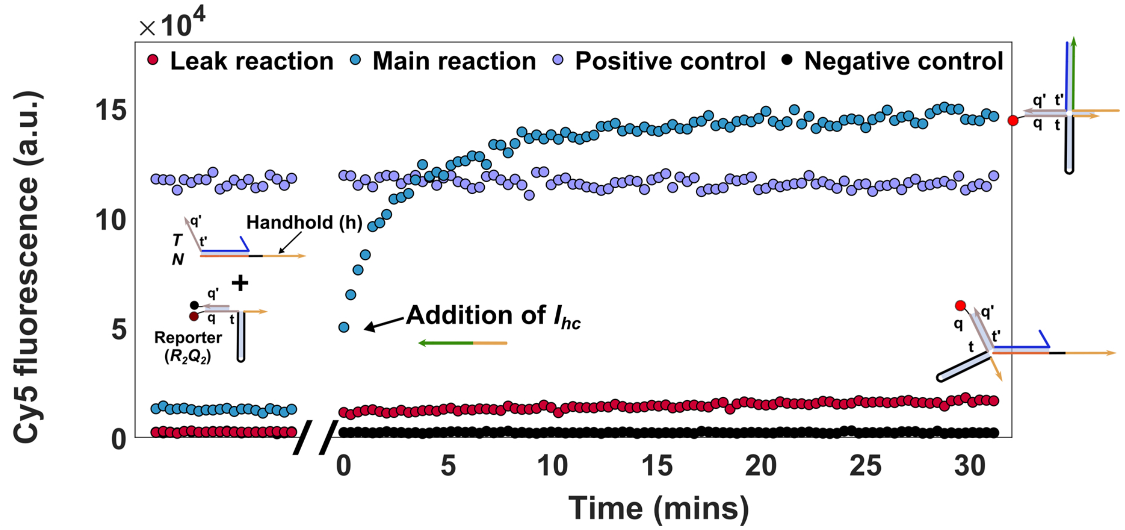

**Supplementary Figure 3| Reaction mechanisms of *$R_2Q_2$***

Raw data from *$R_2Q_2$* characterisation experiments. **Main reaction)** Intended reaction mechanism for *$R_2Q_2$*. A solution of 15 nM of *$R_2Q_2$* is triggered by 10 nM of *N/IT*.. The detached *IT* triggered an associative toehold reaction with a rate of 7.14±0.23.$10^5$ $M^{-1}s^{-1}$ ($k_{r2}$ **Supplementary Figure 20**). **Leak reaction)** Leak produced after adding 15 nM of *$R_2Q_2$* to a solution with 10 nM *TN.* The mere presence of *$R_2Q_2$* and *T* in the same solution triggered a 2 nt toehold-mediated strand displacement with an estimated rate of 4.66±0.89.$10^2$ $M^{-1}.s^{-1}$ (Rate inferred from the mean of the assayed 2 nt toehold-mediated strand displacement reactions at 25˚ C). All concentrations are defined for a reaction volume of 200 µL. Positive control consisted of more than 10 nM of the Cy5-labelled reporter strand. Negative control consisted of only reaction buffer.

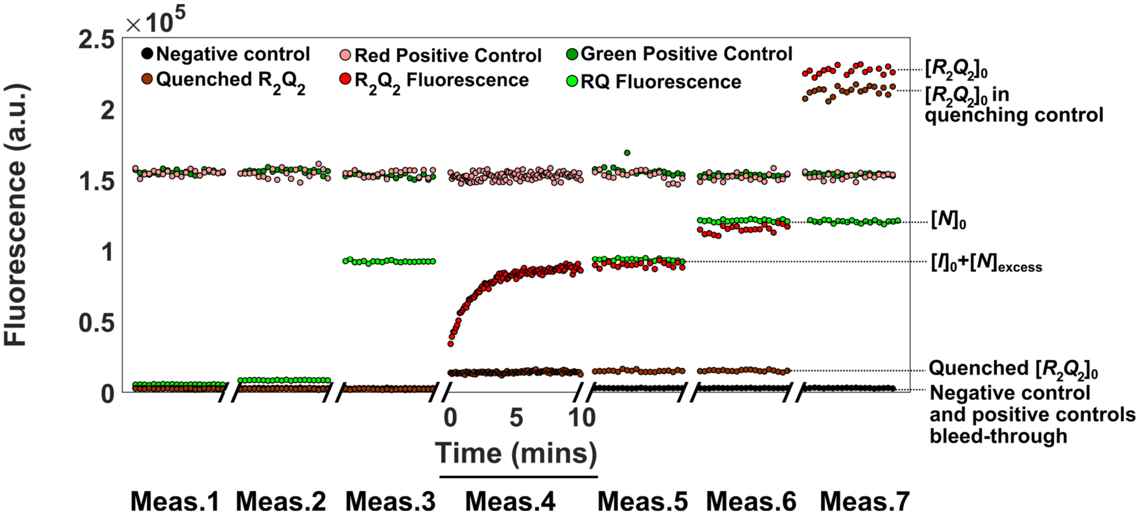

**Supplementary Figure 4| Example of the kinetic analysis of handhold detachment**

Raw data from the reaction system with 7 nt handhold, 2 nt spacer and 3 nt toehold at 25˚C (7/2/3_25) triggered with $I_{hc}$. *T* strands in these experiments always contain an extra domain at 3' that trigger reporter $R_2Q_2$. Measurement 1 contains the baseline of the experiment produced by 15 nM of quenched *RQ*. Then, 10 nM of *TN* was added. $[N]_{\text{excess}}$ was inferred from the annealing increase the baseline, which was recorded in measurement 2. After that, a variable amount of trigger $I_{hc}$ was added, between 9 and 4 nM (8 nM in this case), and after the reaction finished, measurement 3 estimated $[I]_0$, which is equal to the total amount of produce *IT.* During measurement 4, 15 nM of $R_2Q_2$ was added and only red fluorescence was recorded. From this measurement, the signal of the quenched $R_2Q_2$ control was also measured. Measurement 5 recorded again the baseline of green and red fluorescence after the reaction finished. Measurement 6 saturated the solution with 20 nM of $I_{hc}$ to estimate the total amount of *N* in the solution. After that, 20 nM of *T* was added to trigger the rest of $R_2Q_2$ and quantify it in measurement 7. Quenched control was triggered with 20 nM of *IT.* All concentrations are defined for a reaction volume of 200 µL. Green fluorescence was pre-eminently used for species quantification due to its higher signal to noise ratio and red fluorescence to follow the actual reaction kinetics. Positive controls consisted of more than 10 nM of the labelled reporter strands. Negative control consisted of only reaction buffer.

#### Supplementary Note III: Selective production of complexes far-from-equilibrium

The objective of the experiment was to observe the speed difference between the three competing displacement reactions. To minimise the effect produced by any difference in concentration, all *I*-strands and reporter complexes were present at much higher concentrations than the reaction trigger (10 nM $TN_x$). Each set of three experiments was assayed simultaneously using the same aliquots of $I_x$ strands and reporter complexes to ensure consistency between the different conditions. At the end of each experiment, all wells were triggered by adding *T* strand in order to check that all $I_x$ and reporter complexes were present. Steps of the experiments and an example of the fluorescence traces are shown in **Supplementary Table 4** and **Supplementary Figure 5.** Fluorescence in this class of experiments were not calibrated as in previous ones because they did not need to be fitted. Gain was set to arbitrary values that does not saturate for 30 nM of each reporter strand. No bleed-through fluorescence was detected between channels during the experiments.

**Supplementary Table 4| Detailed experimental procedure to produce far-from-equilibrium complexes**

The three $TN_x$ conditions and their controls were triggered by mechanical injection and measured simultaneously. All the tested *TN* were triggered with a 2 nt proximal toehold and contained an 8 nt handhold and a 2 nt spacer.

| Conditions | Addition | Purpose | Expected concentrations in 200 µL (nM) | Volume (µL) |
|---|---|---|---|---|
| **Measurement 1** | 20 µL of each Rep (300 nM) + 130 µL Mix of *I* (38.46 nM each) | Estimate residual fluorescence baseline | $Rep_A$:30; $Rep_B$:30; $Rep_C$:30; $I_A$:25; $I_B$:25; $I_C$:25; | 190 |
| **Measurement 2** | 10 µL $TN_X$ (200 nM) | System kinetics | Evolve with time | 200 |
| **Measurement 3** | - | Give max. fluorescence value for each Rep | The *I* and Rep with the same handhold complementarity as $N_X$ will have reacted. | 200 |
| **Measurement 4** | 20 µL *T* (1000 nM) | Trigger Rep to test that all species were present | $Rep_X/T$:~25 ; Rest of $Rep_x$ could react with *T* free in solution. | 220 |

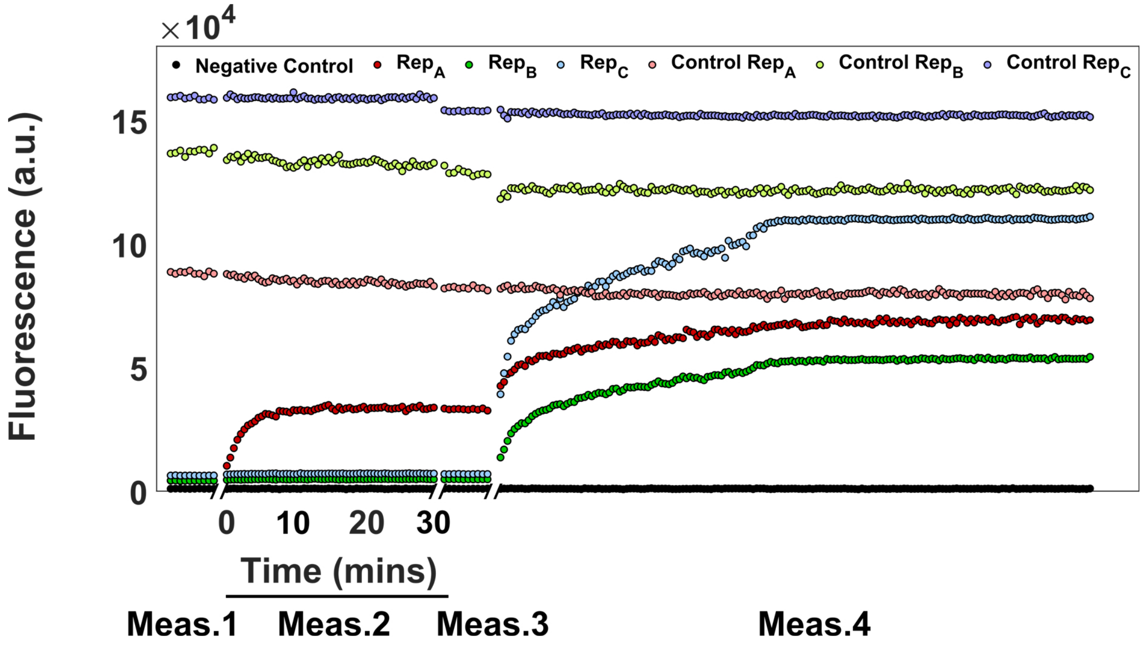

**Supplementary Figure 5| Example of raw experimental data from the production of complex $I_AT$**
The experiment consisted of three competing reaction systems with a different 8 nt handhold, 2 nt spacer and 2 nt toehold at 25˚C (8/2/2_25). Measurement 1 contains the baseline of the experiment produced by 30 nM of each quenched reporter and 25 nM of each $I_x$. Then, 10 nM of $TN_A$ were added, and the kinetics recorded in measurement 2. After the reaction finished, measurement 3 estimated the total amount of produced *IT.* The fluorescence signal obtained for $Rep_A$ was used then to normalise the red fluorescence of all the experiments. After the experiment, in measurement 4, the wells were saturated with 20 nM of *T* to trigger all the reporter complexes. All concentrations are defined for a reaction volume of 200 µL. Positive control consisted of more than 30 nM of the labelled reporter strands. Negative control consisted of only reaction buffer.

#### Supplementary Note IV: Experimental protocol for fluorescence calibration curves

Two fresh aliquots of 1 µM of the fluorophore-labelled reporter strand were created from a stock solution of 100 µM. The aliquots contained 10 µL of the stock solution in 990 µL of TAE 1x buffer (pH=8.3). From each 1 µM aliquot, two more were created with a final concentration of 200 µM, producing a total of four 200 µM aliquots. Four replicas of the calibration curve were pipetted from the 200 µM aliquots, for a range of 1 to 15 µM and a final volume of 160 µL. The 200 µM aliquot used for each point was chosen randomly. The volume was raised up to 200 µL with 40 µL of a solution of 5M NaCl in TAE 1x, for a final reaction buffer of 1M NaCl in TAE 1x. The plate was incubated at the assay temperature for 30 minutes until the fluorescence signal stabilized. Each calibration point was obtained from the median of 10 data points. For each replica of the calibration curve, calibration points were treated as outliers when deviating more than 2 nM from its expected value in a preliminary linear regression.

The photomultiplier gain was set so that its dynamic range was 1.25x the highest fluorescence value in the calibration experiments. Two replicas for the calibration curve of Cy5 contained concentration values up to 16 nM. No bleed-through fluorescence between Cy5 and Cy3 was observed during these calibration experiments.

#### Supplementary Note V: Fitting procedure for fluorescence calibration curves

The value of a negative control consisting of only buffer was subtracted from all the points in the calibration curves. All calibration curves were fitted with a weighted linear regression to a model of the form f(x)=ax. The residuals were weighted with the inverse of the fluorescence reading to account for the Poisson distribution of the photomultiplier counts[1]. Regression coefficient value is reported with 95% confidence bounds. Fitting results and their residuals are shown in **Supplementary Figures 6 to 9.**

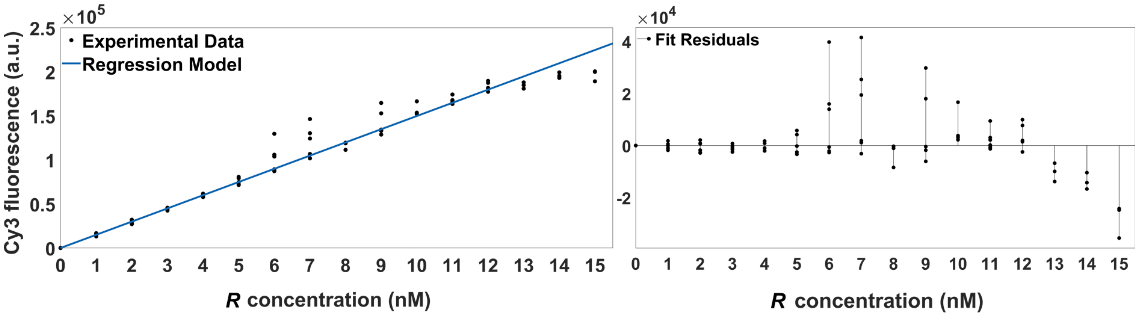

**Supplementary Figure 6| RQ reporter complex (Cy3) calibration curve at 25˚C. Gain 2251**

Used to extrapolate concentrations for all the characterisation experiments at 25˚C. Left: Regression line. Right: Fit residuals. a = 1.498e+04 (1.469e+04, 1.527e+04)**.** Goodness of fit: SSE: 6.915e+04. R-square: 0.9915. Adjusted R-square: 0.9915. RMSE: 27.87.

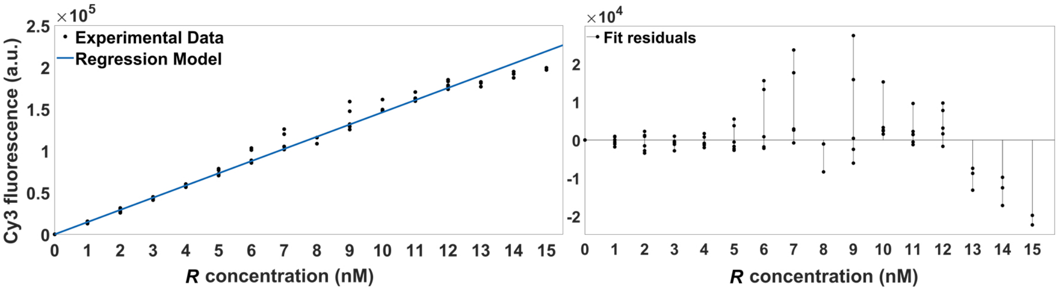

**Supplementary Figure 7| RQ reporter complex (Cy3) calibration curve at 37˚C. Gain 2336**

Used to extrapolate concentrations for all the characterisation experiments at 37˚C. Left: Regression line. Right: Fit residuals. a=1.461e+04 (1.44e+04, 1.481e+04)**.** Goodness of fit: SSE: 3.386e+04. R-square: 0.9955. Adjusted R-square: 0.9955. RMSE: 19.5.

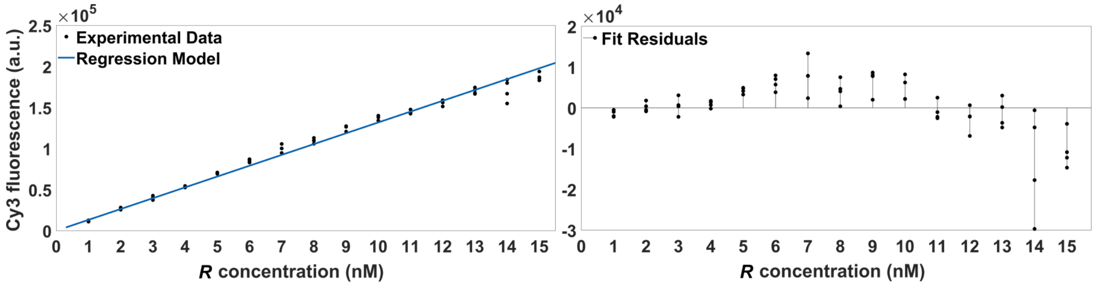

**Supplementary Figure 8|RQ reporter complex (Cy3) calibration curve at 25 degrees. Gain 2230**
Used to extrapolate concentrations for detachment assays. Gain was reduced during this calibration to avoid the frequency of detector saturation events. Left: Regression line. Right: Fit residuals**.** a = 1.319e+04 (1.298e+04, 1.339e+04)**.** Goodness of fit: SSE: 2.189e+04. R-square: 0.9923. Adjusted R-square: 0.9923. RMSE: 19.26.

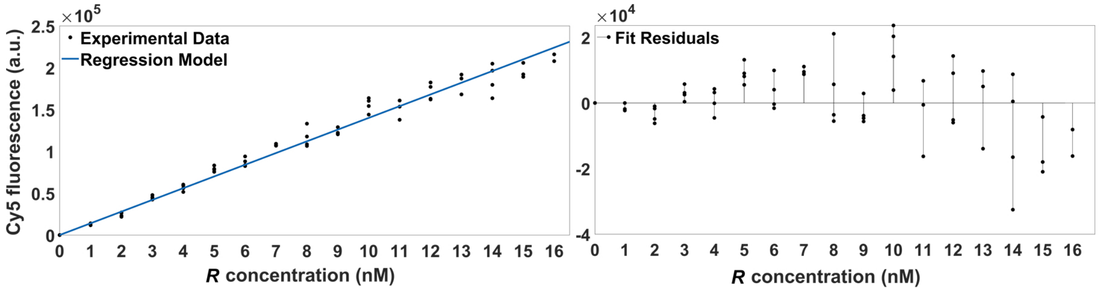

**Supplementary Figure 9| $R_2Q_2$ reporter complex (Cy5) calibration curve at 25 degrees. Gain 2816**
Used to extrapolate concentrations for detachment assays. Left: Regression line. Right: Fit residuals**.** a=1.4e+04 (1.371e+04, 1.43e+04)**.** Goodness of fit: SSE: 4.667e+04. R-square: 0.993. Adjusted R-square: 0.993. RMSE: 27.22.

#### Supplementary Note VI: Fluorescence data processing

To compare calibration lines with the fluorescence signal produced during the experiments, the experimental fluorescence signal was corrected as follows:

Step 1. Removing background fluorescence produced by experimental buffer:
The value of the negative control at each time point was subtracted from the fluorescence value of all wells.

Step 2. Merging all measurements in a single file:
All measurements, except the actual kinetics, were converted in a single data point per well, with its fluorescence value given by the median of all the data points in that set. Then, all the single data points corresponding to the same experiment were merged into a file that also contained the complete kinetics of the experiment.

Step 3. Calculation of the fluorescence variation in the positive control signal:
During longer experiments, the plate could be taken out of the plate reader between measurements or liquids at slightly different temperatures could be injected. These temperature fluctuations could produce distortions in the fluorescence signal, especially for assays at 37˚C. The fluorescence value of the positive control for each data point was used to track such variation of fluorescence signal due to temperature changes during the experiments. The positive control value during the 1$^{st}$ measurement was used as the reference value for the experiment since all wells were at the expected experimental temperature and any photobleaching was assumed to be negligible.

Step 4- Correction of all trajectories with the positive control ratio:
All data points were multiplied by the ratio between the reference control value and the value of the positive control at that time. For experiments running in parallel, a single ratio value at that time was obtained by calculating the mean of the ratio of all positive controls in the plate. The results were corrected traces for each experiment and a completely straight positive control trace (**Supplementary Figure 10**). This approach was applied because the variation of fluorescence produced by temperature changes is proportional to the fluorescent signal[2]

Step 4.1- For quick measurements (*Reporter Characterisation* kinetics):
The correction described in step 4 would be detrimental for quick measurements, where all the fluorescence traces had a high amount of noise, and signal fluorescence artifacts were more common. In addition, since all the kinetics were measured individually, positive controls could not be measured in parallel with them. For these cases, all the datapoints in the kinetics section were multiplied by the ratio between the reference control value and control value obtained after the end of the kinetics in measurement 4 (**Supplementary Note I**)

Step 5- Correction for residual fluorescence:
To infer the concentration of fluorescence species from comparing the fluorescent signal from experiment to the calibration curves, the background due to the residual fluorescence of RQ complexes needed to be subtracted. The background was subtracted from the signal $F_i$ at data point $i$ using the formula:

$$F_i^{*} = F_{max} \frac{F_i - F_1}{(F_{max} - F_1)} \quad (1).$$

Here, $F_i^{*}$ is the data point $i$ with the corrected background fluorescence, $F_1$ the residual fluorescence background value during measurement 1 and $F_{max}$ the final fluorescence value from which the amount of *RQ* was inferred in measurement 6 (**Supplementary Note I**). The method assumes that any difference of fluorescence with the negative control prior to injection of reactant species to the reporter-containing solution is due to unquenched *RQ,* being proportional to $[RQ]_0$. The fluorescence was corrected in such a way that $F_1$ became 0 and $F_{max}$ did not change.

Step 6- Transformation of fluorescence arbitrary units to nM:

Once all the corrections are applied, all the values of the data points are transformed to nM by dividing them by the slope of their corresponding calibration curve (**Supplementary Note V**).

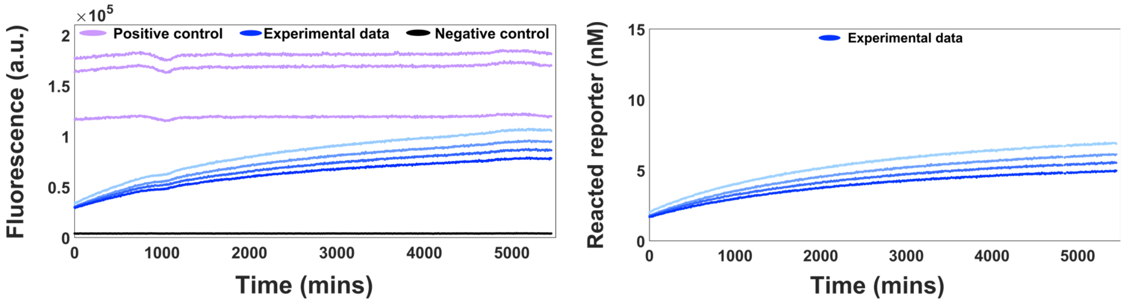

**Supplementary Figure 10| Example of fluorescence data processing**

Four traces from reaction system with 20 nt handhold, 5 nt spacer and 2 nt proximal toehold (20/5/2_25), triggered with [4-7] nM of $I_{wc}$. **Left)** Raw data. Irregularities during the experiment can produce bumps in the fluorescence that tampers the fitting process. These fluctuations appeared in all trajectories and controls contained in the plate. This example contained three different positive controls, with different Cy3-labelled strand concentrations, that presented the same proportional variations in fluorescence. **Right)** The same data after applying the fluorescence processing pipeline. Fluorescence units were transformed to nM and the bumps were corrected. All concentrations are defined for a reaction volume of 200 µL.

#### Supplementary Note VII:Additional fluorescence processing

Out of range fluorescence:

The photomultiplier gain was set so that its dynamic range was 1.25x the fluorescence signal produced by 15 nM of Cy3. However, during some assays, the total amount of *RQ* was slightly over that concentration, getting closer to 90% of the range of the photomultiplier. This resulted in a fluorescence signal over the photomultiplier detection limit when measuring the fluorescence produced by $[RQ]_0$. To correct these out of range data points, the gain was slightly decreased until no saturation events appeared.

After that, the values in the saturated and the low-gain measurements were compared. The ratio between the values in each measurement was calculated for the positive controls and the samples that did not saturate. Assuming that linearity is maintained at the two different gains, the value of the saturated data points was estimated by multiplying the datapoint at low gain by the mean of the obtained ratios. Any error introduced in the concentration of *RQ* using this approximation is considered negligible since small variations in the concentration of *RQ* have a small impact in the fitting procedure for most of the systems. All files modified this way are listed in the file Original_files/Raw_files_modifications.txt found in www.zenodo.org/record/3832588.

Out of range fluorescence during kinetics:

When out-of-range outliers appeared during kinetics, that sample was discarded. For experiments longer than 2 hours, the datapoint with the outlier was removed from all samples.

Estimation of residual fluorescence during kinetic analysis of handhold detachment:

Since during detachment assays $R_2Q_2$ was used to trigger the reaction, the residual fluorescence for each experiment could not be measured. Thus, the quenched control of each experiment was used to calculate the residual fluorescence of $R_2Q_2$ for each trace. The quenching efficiency for $R_2Q_2$ was estimated from the quenched control by calculating the ratio between its residual fluorescence control and its fluorescence value once fully triggered. By multiplying the obtained quenching efficiency by the maximum fluorescence of each trace, the value of residual red fluorescence was estimated for each sample. The obtained value of residual fluorescence of $R_2Q_2$ is used to correct the fluorescence traces with **Supplementary equation 1**, in the same way as with *RQ*.

#### Supplementary Note VIII: Reaction controls for *RQ*

The purpose of this control was to demonstrate that no leak reaction occurred between *TN* and *RQ.* A plate with 18 reactions was set. The 18 wells contained a solution of the *RQ* reporter and *TN* with a 7 nt toehold. 6 different systems were tested in triplicate: 0, 5 and 20 nt long handholds with spacers of 0 and 8 nt. The solution was incubated for more than 24 hours without producing any fluorescent signal. After that, three different concentrations of $I_{hc}$ (4,6 and 8 nM) were added to each system to demonstrate that the produced fluorescence signal was proportional to the $I_{hc}$ added. Steps of the reaction control for *RQ* and the obtained results are shown in **Supplementary Table 5** and **Supplementary Figure 11.** An additional control was assayed to test if the reporter could not be triggered by handhold complementarity alone. The concentrations and systems in the additional control were the same as the main control, but in this case, the added *I* strand only had complementarity with the handhold domain of the *N* and not with *T* (**Invader-random 3', See Supplementary Table 22)**. Fluorescence was measured for more than 30 hours without any increase of fluorescence observed. Steps of the control using *I* without *T* complementarity and the obtained results are shown in **Supplementary Table 6** and **Supplementary Figure 12.**

**Supplementary Table 5| Steps of the reaction control for *RQ***

All the trajectories in the plate and their controls were triggered by mechanical injection and measured simultaneously. All the tested *TN* contained a 7 nt proximal toehold.

| Steps | Addition | Description | Purpose | Volume (µL) |
|---|---|---|---|---|
| **Measurement 1** | 10 µL *RQ* (300 nM) + 10 µL *TN* (200 nM) + 130 µL Buffer | Kinetics during 16.5 hours of the *RQ* basal fluorescence | Demonstrate that the presence of *TN* does not trigger *RQ* | 150 |
| **Measurement 2** | - | Measurement after ~24 hours | Demonstrate signal has not changed | 150 |
| **Measurement 3** | [40/30/20] µL $I_{hc}$ (40 nM) + [10/20/30] µL Buffer | Trigger *TN* | Demonstrate that *I* can trigger the reaction, and check proportionality of fluorescence signal. | 200 |
| **Measurement 4** | 10 µL *I* (400 nM) | Trigger all *TN* | Demonstrate triggering of total *TN* | 210 |

**Supplementary Table 6| Steps of the control for *I* without *T*-complementarity**

All the trajectories in the plate and their controls were triggered by mechanical injection and measured simultaneously. All the tested *TN* contained a 7 nt proximal toehold.

| Steps | Addition | Description | Purpose | Volume (µL) |
|---|---|---|---|---|
| **Measurement 1** | 10 µL *RQ* (300 nM) + 130 µL Buffer | Measure baseline | Remove the residual fluorescence | 140 |
| **Measurement 2** | 10 µL *TN* (200 nM ) | Measure new baseline | Check the signal produced by $[N]_{excess}$ | 150 |
| **Measurement 3** | 50 µL *I* (40 nM) | Add *I* without *T* complementarity. | Demonstrate that handhold interaction does not trigger the reporter. | 200 |
| **Measurement 4** | - | Measure after one day | Demonstrate signal has not changed | 200 |

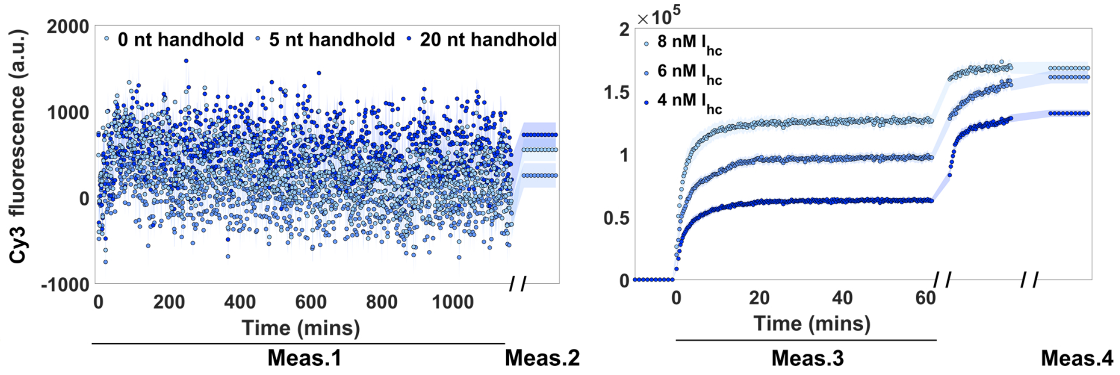

**Supplementary Figure 11| Reporter *RQ* control, demonstrating absence of leak when mixed with unreacted TN**

Processed fluorescence data of 6 different systems tested in triplicate: 0, 5 and 20 nt long handholds with spacers of 0 and 8 nt, and a proximal toehold of 7 nt. Mean of the signal produced for each handhold length with its standard error (shaded). The baseline of fluorescence produced by $[N]_{excess}$ was removed to clearly show the scale of the fluorescence signal noise. Left) After the kinetics of measure 1, measure 2 is taken after approximately one day. No significant increase in the fluorescence signal could be observed. Right) For each replica of each system, a different concentration of $I_{hc}$ were added: 8, 6 and 4 nM. The produced signal is proportional to the concentration of $I_{hc}$, which allows the quantification of the strand. All concentrations are defined for a reaction volume of 200 µL.

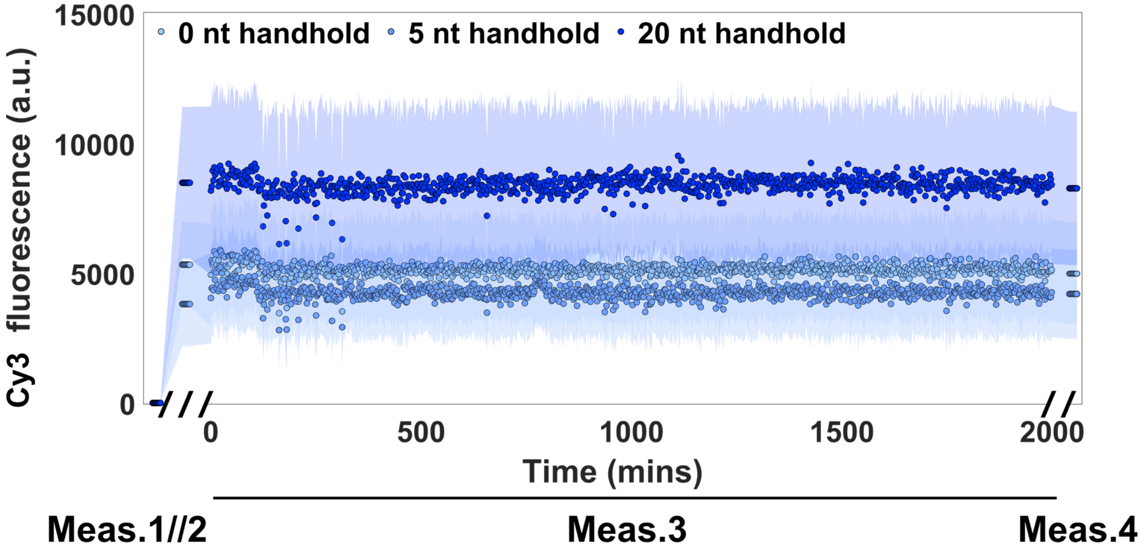

**Supplementary Figure 12| Reporter *RQ* control using *I* without *T*-complementary, demonstrating absence of reaction when using a *I* that is only handhold complementary**

Processed fluorescence data of 6 different systems tested in triplicate: 0, 5 and 20 nt long handholds with spacers of 0 and 8 nt, and a proximal toehold of 7 nt. Mean of the signal produced for each handhold length with its standard error (shaded) for n=9. Measure 1 set the baseline produced by 15 nM of quenched *RQ* and in measure 2, a new baseline produced by the $[N]_{excess}$ after adding 10 nM *TN*. In measurement 3, 10 nM of *I* without *T* complementarity is added and measured for at least 30 hours to observe no increase in the fluorescence signal. Measurement 4 is taken after approximately one day and no increase is observed either. All concentrations are defined for a reaction volume of 200 µL.

#### Supplementary Note IX: Reaction controls for $R_2Q_2$

Five different conditions were assayed in duplicate, testing different qualities of $R_2Q_2$. The used *T* contained a 3 nt proximal toehold and a 5' domain to trigger the reporter $R_2Q_2$. Condition 1 demonstrated that the reporter can be quickly triggered by the addition of the complex *IT.* During the experiments of condition 2, $R_2Q_2$ was quickly triggered when a *TN* with a short handhold (6 nt) reacted with $I_{hc}$. Conditions 3 and 4 showed that the leak reaction of $R_2Q_2$ with *TN* (reported in **Supplementary Note II**) is independent of handhold length. In addition, condition 4 demonstrated that systems with 20 nt handholds cannot react quickly since *IT* cannot detach from the handhold. Finally, condition 5 showed that $R_2Q_2$ does not change its fluorescence signal during 3 hours in the presence of only $I_{hc}$. Concentrations of $R_2Q_2$ greatly differed during these experiments due to an unoptimized annealing process of $R_2Q_2$. However, the qualitative effect of each condition can be observed. Steps for each condition and the obtained results are shown in **Supplementary Table 7** and **Supplementary Figure 13.**

**Supplementary Table 7| Steps of the $R_2Q_2$ control**

All the trajectories in the plate and their controls were triggered by mechanical injection and measured simultaneously. All the tested *TN* contained a 3 nt proximal toehold. Each column of the table contains the species added to the conditions before each measurement.

| Conditions | Measurement 1 | Measurement2 | Measurement 3 | Measurement 4 |
|---|---|---|---|---|
| **Condition 1** | 75 µL $R_2Q_2$ (40 nM) + 75 µL Buffer | 10 µL *IT* (200 nM) + 40 µL Buffer | 10µL $I_{hc}$ (400 nM) | 10µL *IT* (400 nM) |
| **Condition 2** | 75 µL $R_2Q_2$ (40 nM) + + 75 µL Buffer | 10 µL *TN*(6/2/3) (200 nM ) + 40 µL $I_{hc}$ (40 nM) | 10µL $I_{hc}$ (400 nM) | 10µL *IT* (400 nM) |
| **Condition 3** | 75 µL $R_2Q_2$ (40 nM) + + 75 µL Buffer | 10 µL *TN*(0/2/3) (200 nM ) + 40 µL $I_{hc}$ (40 nM) | 10µL $I_{hc}$ (400 nM) | 10µL *IT* (400 nM) |
| **Condition 4** | 75 µL $R_2Q_2$ (40 nM) + + 75 µL Buffer | 10 µL *TN*(20/2/3) (200 nM) + 40 µL $I_{hc}$ (40 nM) | 10µL $I_{hc}$ (400 nM) | 10µL *IT* (400 nM) |
| **Condition 5** | 75 µL $R_2Q_2$ (40 nM) + + 75 µL Buffer | 50 µL $I_{hc}$ (40 nM ) | 10µL $I_{hc}$ (400 nM) | 10µL Buffer |

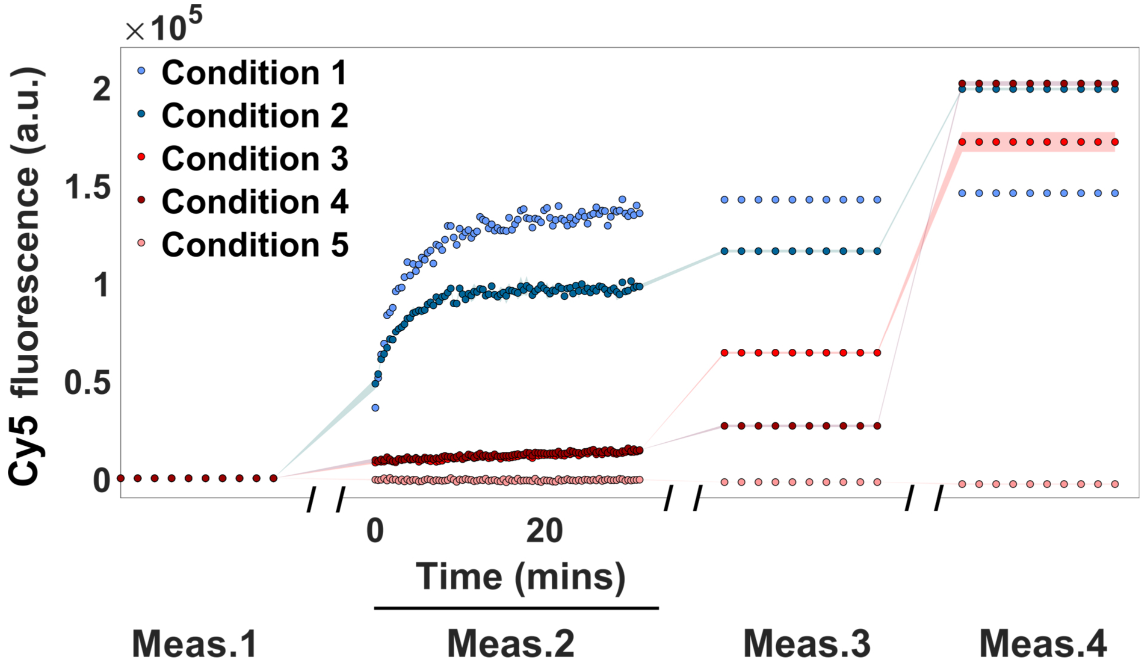

**Supplementary Figure 13| Reporter *$R_2Q_2$* controls**

Processed fluorescence data of five different conditions tested in duplicate (except condition 1). Mean of the signal produced for each condition with its standard error (shaded). The purpose of each condition is described in **Supplementary Note VIII**. Measure 1 set the baseline produced by 15 nM of quenched $R_2Q_2$ and measure 2 contains the kinetics of every condition (Described in **Supplementary Table 7**). Measurement 3 was taken after adding 20 nM $I_{hc}$ to all conditions. During measurement 3, condition 3 did not reach saturation due to the slow reaction rate of the 3 nt proximal toehold reaction. In the case of condition 4, it did not reach saturation because the produced *IT* could not detach from the 20 nt handhold and trigger the reporter. Measurement 4 was taken after adding 20 nM *IT* to all conditions, except condition 5. All concentrations are defined for a reaction volume of 200 µL.

#### Supplementary Note X: Specificity controls for $Rep_A$, $Rep_B$ and $Rep_C$

For the selective production of complexes far-from-equilibrium, not only must the handhold do its part, but the reporters that monitor the reactions ($Rep_A$, $Rep_B$ and $Rep_C$) must be able to discern between the $IT_X$ complexes. The purpose of these controls was to confirm that the designed reporters react only with the $IT_x$ with which they had handhold complementarity. The handholds sequences used during the experiments were: the first 8 nt of sequence used during the HMSD characterisations for $Rep_A$, a similar sequence with the same GC content for $Rep_B$, and the sequence complementary to $I_{wc}$ for $Rep_C$ (**Supplementary Table 23**).

Since the design of the reporters was the same as $R_2Q_2$ ($Rep_A$ being the same reporter) a slow leak reaction was expected in the presence of strand *T*. This leak was slower for $Rep_B$ and $Rep_C$, since these reporters contained a non-*T*-complementary 2 nt clamp at the end of the displaced domain that helped to maintain quencher and fluorescence strands hybridised. For $Rep_A$ this clamp was not needed since the additional unspecific interaction between Cy5 and the quencher was strong enough to stabilize the hybridisation of $Rep_A$ strands[3].

The controls tested the rate at which the reporters reacted with each *TN* complex and if any leak could be produced by the interaction of the $I_x$ strands alone with the reporter. Results showed that during the time that takes the reporters to consume 10 nM of complementary $TN_x$, the amount of non-complementary $TN_x$ consumed is less than 10%. The additions during each step of the control experiments are described in **Supplementary Table 8**. Results for $Rep_A$, $Rep_B$ and $Rep_C$ are shown in **Supplementary Figures 14, 15 and 16,** respectively**.**

**Supplementary Table 8| Steps of the selectivity controls for Rep complexes**

All the trajectories in the plate and their controls were triggered by mechanical injection and measured simultaneously. The used *T* contained a 3 nt proximal toehold and a 5' domain to trigger the reporter $R_2Q_2$. Each column of the table contains the species added to the conditions before each measurement.

| Conditions | Measurement 1 | Measurement 2 | Measurement 3 | Measurement 4 |
|---|---|---|---|---|
| $I_x + TN_x$ | 20 µL Rep (300 nM) + 5 µL $I_x$ (1µM) + 165 µL Buffer | 10 µL $TN_X$ (200 nM) | 20 µL *T* (1 µM) | 5 µL *I*-complementary to the reporter (1 µM) |
| $I_A + I_B + I_C$ | 20 µL *Rep* (300 nM) + 5 µL $I_A$ (1µM) + 5 µL $I_B$ (1µM) + 5 µL $I_C$ (1µM) + 155 µL Buffer | 10 µL Buffer | 20 µL *T* (1 µM) | 5 µL *I*-complementary to the reporter (1 µM) |

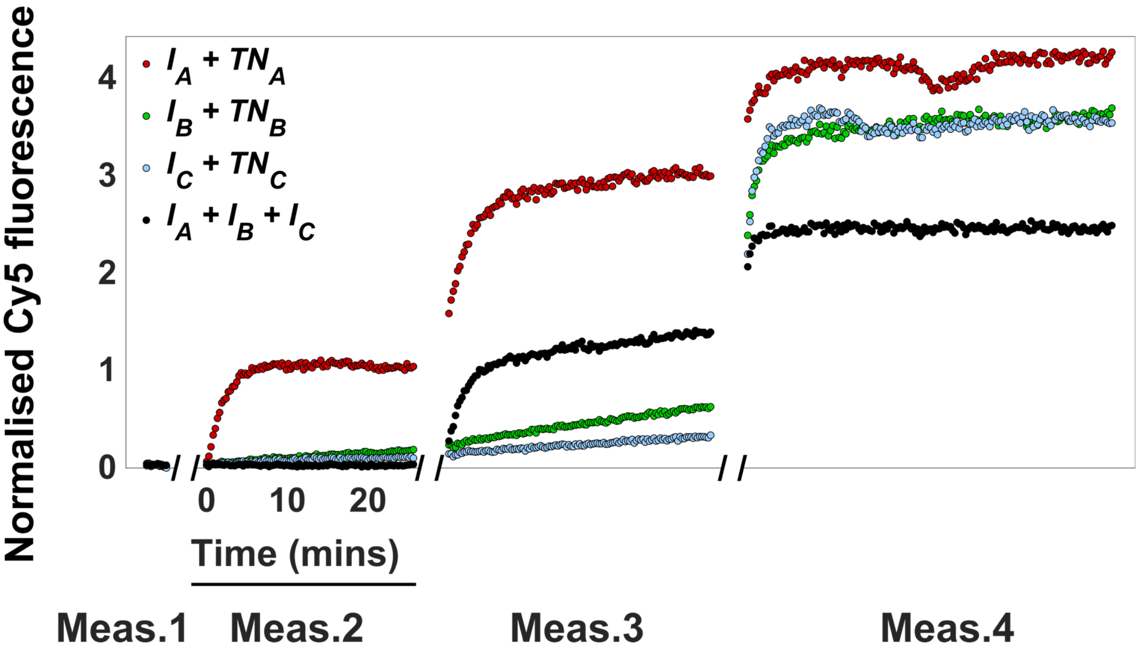

**Supplementary Figure 14| Control to demonstrate Rep$_A$ specificity for $I_AT$**

Processed fluorescence data normalised to the max value of $I_A + TN_A$ condition during measurement 2. Measurement 1 set the baseline produced by 30 nM of quenched Rep$_A$ and 25 nM of $I_x$ (75 nM in total in condition $I_A$+$I_B$+$I_C$). Measurement 2 contains the kinetics of every condition after adding 10 nM of the specified $TN_x$ or just buffer for condition $I_A$+$I_B$+$I_C$. Condition $I_A$+$TN_A$ is the only one capable of reacting at high speed. Measurement 3 was taken after saturating each condition with 100 nM of $T$, so only the wells containing $I_A$ managed to produce fluorescence at an increased rate. In measurement 4, all wells were saturated with 40 nM of $I_A$ to trigger all Rep$_A$. Rep$_A$, in comparison with the other two reporters, had more leakiness, especially with $TN_B$ since handhold A and B have a similar sequence. However, in competition with the other two reporters, the leakiness was greatly reduced. All concentrations are defined for a reaction volume of 200 µL.

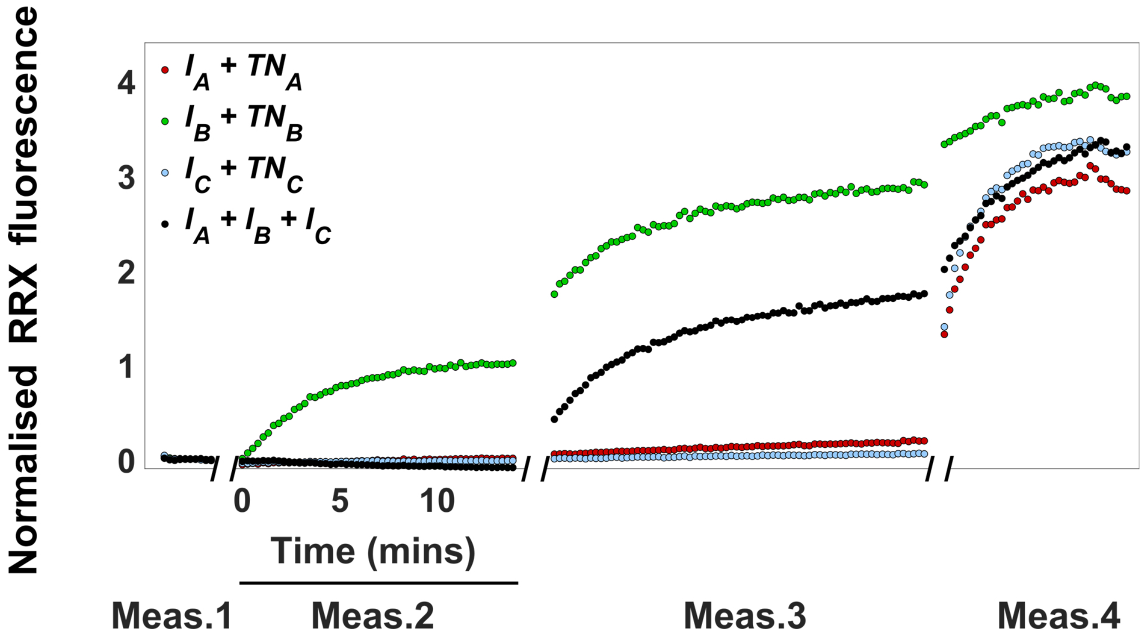

**Supplementary Figure 15| Control to demonstrate Rep$_B$ specificity for $I_BT$**

Processed fluorescence data normalised to the max value of $I_B$ +$TN_B$ condition during measurement 2. Measurement 1 set the baseline produced by 30 nM of quenched Rep$_B$ and 25 nM of $I_x$ (75 nM in total in condition $I_A$+$I_B$+$I_C$). Measurement 2 contains the kinetics of every condition after adding 10 nM of the specified $TN_x$ or just buffer for condition $I_A$+$I_B$+$I_C$. Condition $I_B$+$TN_B$ is the only one capable of reacting at high speed. Measurement 3 was taken after saturating each condition with 100 nM of *T*, so only the wells containing $I_B$ managed to produce fluorescence at an increased rate. In measurement 4, all wells were saturated with 40 nM of $I_B$ to trigger all Rep$_B$. Rep$_B$ showed less leakiness than Rep$_A$, but the similarity between handholds A and B still resulted in a quicker leak reaction for $I_A$ than for $I_C$. All concentrations are defined for a reaction volume of 200 µL.

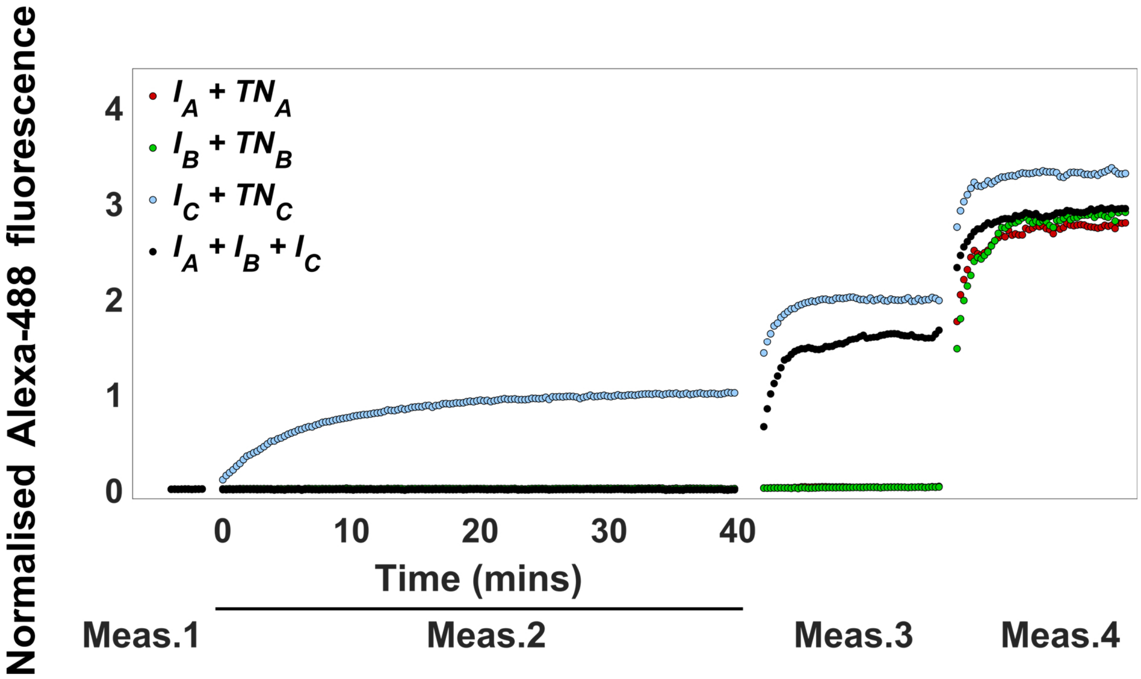

**Supplementary Figure 16| Control to demonstrate $Rep_C$ specificity for *$I_CT$***

Processed fluorescence data normalised to the max value of $I_C$+$TN_C$ condition during measurement 2. Measurement 1 set the baseline produced by 30 nM of quenched $Rep_C$ and 25 nM of $I_x$ (75 nM in total in condition $I_A$+$I_B$+$I_C$). Measurement 2 contains the kinetics of every condition after adding 10 nM of the specified $TN_x$ or just buffer for condition $I_A$+$I_B$+$I_C$. Condition $I_C$+$TN_C$ is the only one capable of reacting at high speed. Measurement 3 was taken after saturating each condition with 100 nM of *T*, so only the wells containing $I_C$ managed to produce fluorescence at an increased rate. In measurement 4, all wells were saturated with 40 nM of $I_C$ to trigger all $Rep_C$. $Rep_C$ did showed no detectable leakiness. All concentrations are defined for a reaction volume of 200 µL.

#### Supplementary Note XI: Implementation of handhold-mediated strand displacement reaction network as an ODE system

The handhold-mediated strand displacement chemical reaction network is composed of the following reactions:

Binding of *I* to *TN*:

$$TN + I \underset{k_d}{\overset{k_b}{\rightleftharpoons}} TN/I\ . \quad (2)$$

*N* displacement by *I*:

$$TN/I \xrightarrow{k_m} N/IT\ . \quad (3)$$

Competing TMSD:

$$TN + I \xrightarrow{k_t} N + IT\ . \quad (4)$$

Blocking and unblocking of *TN*'s handhold by free in solution *IT*:

$$TN + IT \underset{k_d}{\overset{k_b}{\rightleftharpoons}} TN/IT\ . \quad (5)$$

*TN* with blocked handhold reacting via TMSD:

$$TN/IT + I \xrightarrow{k_t} IT + N/IT\ . \quad (6)$$

*RQ* triggering by TMSD.

$$N/IT + RQ \xrightarrow{k_r} RN/IT + Q\ . \quad (7)$$

Produced quencher strand (*Q*) is inert.

Free in solution *IT* attaching and detaching from the handhold in *RN*

$$RN + IT \underset{k_d}{\overset{k_b}{\rightleftharpoons}} RN/IT\ . \quad (8)$$

Free in solution *I* attaching and detaching from the handhold in *RN*

$$RN + I \underset{k_d}{\overset{k_b}{\rightleftharpoons}} RN/I\ . \quad (9)$$

*RN/I* triggering *TN* and *TN/IT* by toehold interaction.

$$TN + RN/I \xrightarrow{k_t} N + RN/IT\ , \quad (10)$$

$$TN/IT + RN/I \xrightarrow{k_t} RN/IT + N/IT\ . \quad (11)$$

The ODE model does not include inert species (*Q*), or those which concentration is negligible for all reaction systems (*N*). Counting the *N* molecule produced by direct TMSD (**Supplementary equation 4**) as a distinct species greatly increases the complexity of the ODE model that results from this reaction network. Since $k_r$ and $k_b$ are always large, we assume for simplicity that direct TMSD instead produces the complex $N/IT$, which swiftly binds to the reporter (**Supplementary equations 7 and 10**). Subsequent detachment of *IT* from the triggered reporter complex *RN* is then accounted for by the backwards reaction in **Supplementary equation 8**. This approximation is likely to be worst when both the toehold and handhold are long. In this limit, toehold-based displacements will produce *N* relatively rapidly, and this *N* can act to sequester unreacted invaders via its handhold, either before or after triggering the reporter. In our approximation, the *N* species will instead remain bound to the reacted invader-target complex, and cannot perform sequestration of unreacted invader. However, regions of long proximal toeholds are not the primary focus of this work, and any discrepancies introduced are likely small on the scale of over four orders of magnitude considered here. In particular, these sequestering species only appear over the course of the reaction, and most invaders will stably bind to a handhold early in the process.

No base-pair-level intermediates were considered explicitly for any of the branch migration reactions. The resultant ODE model was composed of the mass-action law based differential equations:

$$\frac{d[TN/I]}{dt} = k_b[TN][I] - k_d[TN/I] - k_m[TN/I], \quad (12)$$

$$\frac{d[TN/IT]}{dt} = k_b[TN][IT] - k_d[TN/IT] - k_t[TN/IT][I], \quad (13)$$

$$\frac{d[N/IT]}{dt} = k_m[TN/I] + k_t[TN][I] + k_t[TN/IT][I] - k_r[N/IT][RQ] + k_t[RN/I][TN] + k_t[RN/I][TN/IT], \quad (14)$$

$$\frac{d[IT]}{dt} = k_t[TN/IT][I] + k_d[TN/IT] + k_d[RN/IT] - k_b[TN][IT] - k_b[RN][IT], \quad (15)$$

$$\frac{d[RN]}{dt} = k_d[RN/I] + k_d[RN/IT] - k_b[RN][I] - k_b[RN][IT] + k_t[RN/I][TN], \quad (16)$$

$$\frac{d[RN/I]}{dt} = k_b[RN][I] + k_b[RN][IT] - k_d[RN/I] - k_d[RN/IT] - k_t[RN/I][TN] - k_t[RN/I][TN/IT], \quad (17)$$

$$\frac{d[RN/IT]}{dt} = k_r[N/IT][RQ] + k_b[RN][IT] - k_d[RN/IT] + k_t[RN/I][TN/IT]. \quad (18)$$

With conservation laws for the initial species:

$[I] = [I]_0 - [TN/I] - [TN/IT] - [N/IT] - [IT] - [RN/I] - [RN/IT]$, (19)

$[TN] = [TN]_0 - [TN/I] - 2[TN/IT] - [N/IT] - [IT] - [RN/IT]$, (20)

$[RQ] = [RQ]_0 - [RN] - [RN/I] - [RN/IT]$, (21)

And initial conditions: $[TN/I]_0$, $[TN/IT]_0$, $[N/IT]_0$, $[IT]_0$, $[RN/I]_0$, $[RN/IT]_0 = 0$ and $[RN] = [N]_{excess}$, being $[N]_{excess}$ the excess of *N* resultant from the inexact stoichiometry when annealing *TN* with a 10% excess of *N*. $[N]_{excess}$, $[I]_0$, $[N]_0 = [TN]_0 + [N]_0$ and $[RQ]_0$ were inferred from the fluorescence signal produced during characterisation experiments as described in **Supplementary Note I**.

#### Supplementary Note XII: General fitting procedure

All fluorescence data were first corrected as described in **Supplementary Note VI** and the baseline produced by $[N]_{excess}$ subtracted. All fitting procedures included, in addition to their rate constants, the reaction trigger concentration and $t_0$ for each trajectory.

The initial guesses for the reaction trigger concentrations were inferred from the fluorescence value of measurement 4. The last value of the recorded kinetics in measurement 3 was compared to its value in measurement 4 to determine the percentage of completion of the reaction during recording in measurement 3. For kinetics that did not reach at least 90% completion, the concentration of reaction trigger was only allowed to fluctuate ±10% from its initial guess.

The initial guess for $t_0$ ($t_{0\_init}$) was obtained from the cutting point of a regression line with the x-axis, except during *reporter RQ fitting* (**Supplementary Note XIII**). The regression line was calculated by fitting the values of the kinetics that were under 50% completion, where the trajectories were almost linear.

Fittings were performed with the MATLAB® function *fminsearch*, based on the simplex search method by Lagarias et al.[4] The decimal logarithm of the error function was minimised for all the fluorescence traces of a given system:

$$log_{10}(mean\ weighted\ error) = log_{10}\frac{\sum_i \sum_t \frac{(f_{exp}(i,t)-f_{model}(i,t))^2}{f_{model}(i,t)}}{n}\ , (22)$$

where $f_{exp}(i,t)$ is the inferred concentration at time $t$ of the trace $i$, and $f_{model}(i,t)$ is the sum of concentrations of all fluorescent species predicted by the model at time $t$ at the reagent conditions of the trace $i$. The error at each point was weighted by $f_{model}(i,t)$ to account for the Poisson distribution of the error for the readings of the photomultiplier as done by Zhang et al.[1] . When more than one trace was fitted simultaneously, the mean total weighted error was calculated by dividing the total error by the number of traces (*n)* that were fitted simultaneously.

All MATLAB fitting scripts are available for download at www.zenodo.org/record/3832588.

#### Supplementary Note XIII: Reporter *RQ* fitting

The experimental value of $k_r$ was obtained for every species of *N* used during the characterisation experiments. Trajectories were fitted individually to **equation 6** in the main text. Since all reactions were completed in the span of a minute, the effect of the initial leak on $t_0$ was not significant. Thus, the time of *N/IT* injection was considered $t_{0\_init}$ .

The data was fitted in a three-step iteration. During the 1st round of fittings, the initial guess $k_{r\_init}$ was inferred from the time at which the reaction reached 90% completion:

$$k_{r\_init} = \frac{ln(10 - 9\frac{[N]_0}{[RQ]_0})}{t_{90\%}([RQ]_0 - [N]_0)} , \qquad (23)$$

with $t_{90\%}$ being the time at which 90% of $[I]_0$ reacted, and $[N]_0$ the total amount of strand *N* injected in the form of *N/IT.* During the 2nd round of fittings, all the trajectories were fitted using as $k_{r\_init}$ the mean of the $k_r$ obtained for all the conditions at that temperature during the first round ($k_{r\_init}$ at 25˚C: $5.10^6$ $s^{-1}M^{-1}$ ; $k_{r\_init}$ at 37˚C: $4.8.10^6$ $s^{-1}M^{-1}$). For the 3rd round, all traces were fitted using as initial guesses their results of the second round for $k_r$ , $t_0$ and $[N]_0$. All fitted $k_r$ at 25 ˚C and 37 ˚C are collected in **Supplementary Tables 9 and 10**, respectively

**Supplementary Table 9| Fitted $k_r$ at 25˚C**

Data obtained using procedures described in **Supplementary Notes I and XIII.** The reactant strands used were all *N* strands "Incumbent x hh/x sp" (**Supplementary Table 20**) with "Target (7 th)" (**Supplementary Table 21**) and "Invader ($I_{hc}$)" (**Supplementary Table 22**). Reporter strands were "Reporter 1 (*R*)" and "Reporter 1 quencher (*Q*)" (**Supplementary Table 23**). Mean of all fitted $k_r$ = $5.068.10^6\,s^{-1}M^{-1}$ ; Standard deviation = $0.505.10^6 s^{-1}M^{-1}$ ; Standard mean error (SEM) = $0.052.10^6 s^{-1}M^{-1}$.

| *Condition (handhold/spacer)* | *Individual experiments $k_r.10^6$ ($s^{-1}M^{-1}$)* | | | | *Mean $k_r.10^6$ ± SEM ($s^{-1}M^{-1}$)* |
|---|---|---|---|---|---|
| **0/0** | 5.46 | 5.42 | 5.55 | 5.60 | 5.506±0.041 |
| **0/1** | 5.41 | 6.11 | 5.54 | 6.02 | 5.77±0.17 |
| **0/2** | 5.18 | 5.18 | 5.24 | 5.09 | 5.17±0.03 |
| **0/3** | 5.45 | 4.99 | 5.08 | 5.04 | 5.14±0.1 |
| **0/5** | 5.42 | 5.63 | 5.63 | 5.79 | 5.617±0.076 |
| **0/8** | 5.51 | 5.29 | 5.43 | 5.21 | 5.359±0.069 |
| **4/2** | 5.47 | 5.25 | 5.44 | 4.08 | 5.06±0.33 |
| **5/0** | 5.32 | 5.06 | 5.63 | 6.11 | 5.55±0.22 |
| **5/1** | 3.95 | 4.21 | 4.27 | 4.58 | 4.25±0.13 |
| **5/2** | 4.66 | 4.35 | 4.33 | 5.68 | 4.76±0.32 |
| **5/3** | 5.42 | 5.69 | 4.49 | 5.38 | 5.24±0.26 |
| **5/5** | 5.63 | 5.61 | 5.98 | 5.69 | 5.728±0.086 |
| **5/8** | 5.08 | 4.63 | 5.03 | 4.65 | 4.85±0.12 |
| **6/2** | 4.46 | 4.91 | 4.53 | 4.52 | 4.61±0.1 |
| **7/2** | 5.00 | 4.85 | 4.94 | 5.10 | 4.97±0.51 |
| **8/2** | 4.82 | 4.55 | 4.72 | 5.05 | 4.78±0.1 |
| **9/2** | 4.60 | 4.54 | 4.71 | 4.83 | 4..67±0.064 |
| **10/2** | 4.80 | 4.83 | 6.01 | 4.80 | 5.11±0.30 |
| **20/0** | 3.97 | 4.13 | 3.77 | 4.14 | 4.004±0.087 |
| **20/1** | 5.24 | 5.06 | 5.45 | 5.31 | 5.265±0.081 |
| **20/2** | 4.87 | 4.96 | 5.11 | 4.75 | 4.922±0.078 |
| **20/3** | 4.91 | 5.01 | 5.00 | 4.80 | 4.931±0.049 |
| **20/5** | 5.13 | 5.06 | 5.47 | 5.58 | 5.31±0.13 |
| **20/8** | 5.12 | 5.12 | 5.12 | 4.9 | 5.08±0.041 |

**Supplementary Table 10| Fitted $k_r$ at 37˚C**

Data obtained using procedures described in **Supplementary Notes I and XIII.** The reactant strands used were all *N* strands “Incumbent x hh/x sp” except the ones with spacer 1,3,5 nt (**Supplementary Table 20**) with “Target (7 th)” (**Supplementary Table 21**) and “Invader ($I_{hc}$)” (**Supplementary Table 22**). Reporter strands were “Reporter 1 (*R*)” and “Reporter 1 quencher (*Q*)” (**Supplementary Table 23**). Legend: Red highlight: $k_r$ values that were not obtained from direct experimental evidence. These values were estimated from the mean of all the obtained $k_r$ at 37˚C. Mean of all $k_r$ = 4.52.10$^6$s$^{-1}$M$^{-1}$ ; Standard deviation = 0.53.10$^6$s$^{-1}$M$^{-1}$ ; Standard mean error (SEM) = 0.11.10$^6$s$^{-1}$M$^{-1}$

| *Condition (handhold/spacer)* | *Individual experiments $k_r$.10$^6$ (s$^{-1}$M$^{-1}$)* | | | | *Mean $k_r$.10$^6$ ± SEM (s$^{-1}$M$^{-1}$)* |
|---|---|---|---|---|---|
| **0/0** | 4.38 | 4.61 | 4.92 | 5.15 | 4.76±0.17 |
| **0/8** | 4.66 | 4.87 | 4.93 | 5.27 | 4.94±0.13 |
| **4/2** | - | - | - | - | 4.52±0.11 |
| **5/0** | 4.65 | 4.57 | 4.84 | 4.64 | 4.677±0.057 |
| **5/2** | - | - | - | - | 4.52±0.11 |
| **5/8** | 4.07 | 4.26 | 4.54 | 4.88 | 4.44±0.18 |
| **6/2** | - | - | - | - | 4.52±0.11 |
| **7/2** | - | - | - | - | 4.52±0.11 |
| **8/2** | - | - | - | - | 4.52±0.11 |
| **9/2** | - | - | - | - | 4.52±0.11 |
| **10/2** | - | - | - | - | 4.52±0.11 |
| **20/0** | 3.29 | 3.52 | 3.76 | 3.54 | 3.525±0.096 |
| **20/2** | - | - | - | - | 4.52±0.11 |
| **20/8** | 4.63 | 4.84 | 4.76 | 4.95 | 4.794±0.069 |

#### Supplementary Note XIV: TMSD fitting

To obtain the value of $k_t$ the trajectories of each system were fitted individually to a simplified ODE model consisting of **Supplementary equations 14 and 18,** assuming $[TN/IT]$, $k_b$, $k_d$ and $k_m$ =0.

$$\frac{d[N]}{dt} = k_t[TN][I] - k_r[N][RQ]\,, \quad (24)$$

$$\frac{d[RN]}{dt} = k_r[N][RQ]\,. \quad (25)$$

The data was fitted iteratively in three steps. During the 1st round of fittings, $k_{t\_init}$ was obtained using a modification of **Supplementary equation 23**, exchanging $[N]_0$ for $[I]_0$, and $[RQ]_0$ for $[TN]_0$. For kinetics that did not reach 90% of completion, $k_{t\_init}$ was obtained from the generalized form of **Supplementary equation 23**:

$$k_{t\_init} = \frac{ln\left(\frac{[I]_0}{[I]_{end}} + \frac{[I]_0}{[TN]_0}\left(1 - \frac{[I]_0}{[I]_{end}}\right)\right)}{t_{end}[TN]_0 - [I]_0}\,, \qquad (26)$$

with $[I]_{end}$ being the maximum $[I]$ value recorded during the kinetics and $t_{end}$ the final time of the measurement (corrected with $t_{0\_init}$). During the 2nd round of fittings, each system fluorescence traces were fitted using as $k_{t\_init}$ the median of all the $k_t$ obtained for that system during the 1st round. For the 3rd round, all trajectories were fitted using as initial guesses its results of the second round for $k_t$ , $t_0$ and $[I]_0$.

Reactions triggered with $I_{hc}$, but in the absence of handholds in $N$ were also fitted following this procedure to confirm the value of $k_t$ obtained with $I_{wc}$. All fitted $k_t$ are collected in **Supplementary Tables 15 to 17.**

#### Supplementary Note XV: $k_b$ estimation

For reaction systems located in the 2nd order regime with irreversible handhold-binding (handhold=20 nt; toehold>1 nt), $k_d$ is considered negligible and $k_m$ very large, and thus the reaction rate is solely determined by $k_b$ (**equation 3**). Thus, the fluorescence traces of system confidently located in the 2nd order regime (20 nt handhold; >1 nt toehold) were individually fitted to the complete ODE model described in **Supplementary Note XI** to obtain an estimation of $k_b$.

During the fitting, $k_m$ was fixed to an arbitrary high value ($10^8$ $s^{-1}$) and $k_d$ to $1.10^{-22}$ $s^{-1}$. The data was fitted in a two-step iteration. During the 1st round of fittings, $k_{b\_init}$ was arbitrarily set to $10^7$ $s^{-1}M^{-1}$. For the 2nd round, all trajectories were fitted using as initial guesses its results of the 1st round for $k_b$ , $t_0$ and $[I]_0$.

For the subsequent fittings, the value of $k_b$ at each temperature was constrained to a range of twice the standard deviation of $k_b$ from these fittings, centred on its mean value. All $k_b$ and their ranges at 25˚C and 37˚C are collected in **Supplementary Tables 11 and 12**, respectively

**Supplementary Table 11| $k_b$ fitting at 25 ˚C**

Data obtained using procedures described in **Supplementary Notes II and XV.** The reactant strands used were all *N* strands "Incumbent 20 hh/x sp" (**Supplementary Table 20**) with *T* strands "Target (x th)" from 2 to 7 nt (**Supplementary Table 21**) and "Invader ($I_{hc}$)" (**Supplementary Table 22**). Reporter strands were "Reporter 1 (*R*)" and "Reporter 1 quencher (*Q*)" (**Supplementary Table 23**). Mean of all $k_b$ = $5.14.10^6 s^{-1}M^{-1}$ ; Standard deviation = $1.27.10^6 s^{-1}M^{-1}$ ; Standard mean error (SEM) = $0.17.10^6 s^{-1}M^{-1}$.

| *Condition (handhold/spacer/toehold)* | *Individual experiments $k_b$ $10^6$ ($s^{-1}M^{-1}$)* | | | | *Mean $k_b$ $10^6$ ± SEM ($s^{-1}M^{-1}$)* |
|---|---|---|---|---|---|
| **20/0/7** | 6.41 | 6.31 | 5.87 | 7.87 | **6.62±0.44** |
| **20/0/2** | 3.47 | 3.49 | 3.21 | 3.51 | **3.421±0.071** |
| **20/1/7** | 7.68 | 6.87 | 6.92 | 6.83 | **7.07±0.2** |
| **20/1/2** | 5.44 | 5.22 | 5.77 | 5.24 | **5.42±0.13** |
| **20/2/7** | 5.74 | 5.74 | 5.44 | 5.95 | **5.72±0.1** |
| **20/2/5** | 7.60 | 7.66 | 7.88 | 6.92 | **7.52±0.21** |
| **20/2/3** | 4.91 | 4.86 | 4.87 | 4.47 | **4.78±0.1** |
| **20/2/2** | 5.55 | 5.28 | 5.52 | 5.29 | **5.411±0.071** |
| **20/3/7** | 6.22 | 4.89 | 5.00 | 4.52 | **5.16±0.37** |
| **20/3/2** | 4.90 | 4.69 | 4.59 | 4.46 | **4.659±0.092** |
| **20/5/7** | 3.87 | 3.55 | 3.61 | 3.40 | **3.609±0.098** |
| **20/5/2** | 5.17 | 4.23 | 4.39 | 3.98 | **4.44±0.26** |
| **20/8/7** | 4.12 | 3.99 | 4.19 | 4.12 | **4.105±0.044** |
| **20/8/2** | 4.47 | 3.82 | 3.84 | 4.08 | **4.05±0.15** |

**Supplementary Table 12| $k_{bind}$ fitting at 37 ˚C**

Data obtained using procedures described in **Supplementary Notes II and XV.** The reactant strands used were all *N* strands "Incumbent 20 hh/x sp" (**Supplementary Table 20**) with *T* strands "Target (x th)" from 2 to 7 nt (**Supplementary Table 21**) and "Invader ($I_{hc}$)" (**Supplementary Table 22**). Reporter strands were "Reporter 1 (*R*)" and "Reporter 1 quencher (*Q*)" (**Supplementary Table 23**). Legend: Red: Rates with a standard error bigger than the 10% of the mean. Corresponds with long experimental conditions (toehold = 0 nt) or systems at the boundary between different kinetic regimes. Mean of all $k_b$ = $1.10^7 s^{-1}M^{-1}$ ; Standard deviation = $0.45.10^7 s^{-1}M^{-1}$ ; Standard mean error (SEM) = $0.11.10^7 s^{-1}M^{-1}$.

| *Condition (handhold/spacer)* | *Individual experiments $k_b$ $10^6$ ($s^{-1}M^{-1}$)* | | | | *Mean $k_b$ $10^6$ ± SEM ($s^{-1}M^{-1}$)* |
|---|---|---|---|---|---|
| **20/0/7** | 13.73 | 15.82 | 17.81 | 17.84 | **16.3±0.98** |
| **20/0/2** | 10.47 | 10.66 | 10.18 | 11.25 | **10.64±0.23** |
| **20/8/7** | 6.09 | 6.68 | 2.90 | 3.07 | **4.69±0.99** |
| **20/8/2** | 7.94 | 8.98 | 9.23 | 8.59 | **8.68±0.28** |

#### Supplementary Note XVI: HMSD fitting

The parameters $k_d$, $k_m$, $k_b$, $[I]_0$ and $t_0$ were fitted simultaneously for each set of HMSD trajectories. All the trajectories for each handhold, toehold and spacer lengths were fit to a single set of values for $k_d$, $k_m$ and $k_b$. Constraints for the value of $k_b$ and the value for the rest of the constants of the model were obtained experimentally (**See Supplementary Notes XIII, XIV and XV**). The data was fitted iteratively in three steps. During the 1st round of fittings, $k_{d\_init}$ was calculated with an estimation of handhold-binding free energy for each handhold and spacer combination[5], using the equilibrium equation:

$$k_{d\_init} = \frac{k_b}{e^{-\left(\frac{\Delta G_{hb}}{RT}\right)}}, \qquad (27)$$

where $R$ is the gas constant (1.987 cal mol$^{-1}$ K$^{-1}$), $T$ is the experimental temperature and $\Delta G_{hb}$ is the estimation of the handhold-binding Gibbs free energy in calories per mol[5]. All the estimated $\Delta G_{hb}$ and $k_{d\_init}$ at 25˚C and 37˚C are collected in **Supplementary Tables 13 and 14**, respectively. The value of $k_d$ was constrained during the fitting to a range $\pm\sqrt{10}\ k_{d\_init}$. $k_{m\_init}$ was changed iteratively in the range of [$10^{-6}$ – $10^{4}$] one order of magnitude each time and the best fitting was stored. For the 2nd round, the trajectories were fitted without constraining $k_d$, using the best result of the 1st fitting round as initial guesses for $k_d$, $k_b$, $k_m$, $t_0$ and $[I]_0$. The 3rd fitting round used the results of the 2nd fitting round as initial guesses. Results from this fitting procedure and their calculated reaction rates and half-lives at 25˚C and 37˚C are collected in **Supplementary Tables 15, 16 and 17.**

To estimate an error for the obtained rates for each system, each trajectory was individually fitted using the result of the fitting process as initial guesses for $k_d$, $k_b$, $k_m$, $t_0$ and $[I]_0$. For reactions with 2nd order kinetics with reversible handhold binding, we report the standard error for the obtained 2nd order rate $k$ (**equation 3**), since we assume compensatory errors between the ratio $\frac{k_m}{k_m + k_d}$ and $k_b$. For reactions with 2nd order kinetics with irreversible handhold binding, we report the standard error for the obtained $k_b$ and for reactions with 1st order kinetics, we report the standard error for the obtained $k_m$, because they are the reaction constant determining the reaction rate in each condition.

**Supplementary Table 13| Estimated $\Delta G_{hb}$ and $k_d$ for each handhold at 25 ˚C**

Data obtained as described in **Supplementary Notes II and XVI.** The reactant strands used were all *N* strands "Incumbent x hh/x sp" (**Supplementary Table 20**) with all *T* strands "Target (x th)" (**Supplementary Table 21**) and "Invader ($I_{hc}$)" (**Supplementary Table 22**). Reporter strands were "Reporter 1 (*R*)" and "Reporter 1 quencher (*Q*)" (**Supplementary Table 23**).

| ***Condition (handhold/spacer)*** | **$\Delta G_{hb}$ (*Kcal M$^{-1}$*)** | ***$k_{d\_init}$ ($s^{-1}$)*** | ***Mean fitted $k_d$ for each condition ($s^{-1}$)*** | | | | | | ***Mean $k_d$ ± SEM ($s^{-1}$)*** |
|---|---|---|---|---|---|---|---|---|---|
| **4/2** | -7.24 | 2.48E+01 | 7.82E+00 | 1.29E+01 | 1.13E+01 | 3.07E+01 | 4.03E+01 | 8.43E+00 | 1.86±0.55E+1 |
| **5/0** | -7.26 | 2.40E+01 | 1.34E+01 | - | - | 2.11E+01 | - | 7.67E+00 | 1.41±0.39E+1 |
| **5/1** | -7.96 | 7.35E+00 | 2.11E+00 | - | - | 7.80E+02 | - | 2.57E+00 | 2.6±2.6E+2 |
| **5/2** | -8.03 | 6.53E+00 | 8.27E+00 | 1.70E+01 | 3.19E+00 | 1.96E+01 | 9.81E+00 | 2.46E+00 | 1.01±0.29E+1 |
| **5/3** | -7.96 | 7.35E+00 | 2.63E+00 | - | - | 9.16E+00 | - | 9.56E+00 | 7.1±2.2E+0 |
| **5/5** | -7.74 | 1.07E+01 | 6.00E+00 | - | - | 7.62E+00 | - | 1.37E-01 | 4.6±2.3E+0 |
| **5/8** | -7.96 | 7.35E+00 | 2.59E+00 | - | - | 4.70E+00 | - | 2.79E+00 | 3.36±0.67E+0 |
| **6/2** | -10.22 | 1.62E-01 | 5.14E-01 | 1.56E-01 | 2.87E-02 | 2.07E-01 | 8.92E-01 | 7.95E-02 | 3.1±1.4E-1 |
| **7/2** | -11.18 | 3.21E-02 | 1.63E-02 | 5.90E-02 | 9.77E-02 | 4.72E-02 | 2.07E-03 | 8.26E-02 | 5.1±1.5E-2 |
| **8/2** | -12.39 | 4.16E-03 | 1.60E-02 | 9.74E-03 | 7.65E-02 | 1.23E-01 | 1.76E-02 | 2.63E-04 | 4.1±2E-2 |
| **9/2** | -15.04 | 4.75E-05 | 8.88E-14 | 2.37E-11 | 9.01E-09 | 6.09E-09 | 3.73E-09 | 4.08E-06 | 6.8±6.8E-7 |
| **10/2** | -15.83 | 1.25E-05 | 2.89E-12 | 5.80E-03 | 6.13E-03 | 3.79E-03 | 4.90E-04 | 1.52E-03 | 3±1.1E-3 |
| **20/0** | -29.72 | 8.22E-16 | 1.04E-20 | - | - | 6.59E-16 | - | 3.81E-16 | 3.5±1.9E-16 |
| **20/1** | -30.42 | 2.52E-16 | 3.01E-16 | - | - | 3.72E-16 | - | 1.17E-16 | 2.63±0.76E-16 |
| **20/2** | -30.5 | 2.20E-16 | 2.39E-16 | 6.68E-16 | 1.46E-16 | 5.41E-16 | 8.03E-17 | 8.14E-17 | 2.9±1E-16 |
| **20/3** | -30.42 | 2.52E-16 | 9.14E-17 | - | - | 4.10E-16 | - | 1.27E-16 | 2.1±1E-16 |
| **20/5** | -30.21 | 3.59E-16 | 1.69E-16 | - | - | 2.58E-16 | - | 1.67E-16 | 1.98±0.3E-16 |
| **20/8** | -30.42 | 2.52E-16 | 2.65E-16 | - | - | 5.73E-16 | - | 2.17E-16 | 3.5±1.1E-16 |
| | ***Toehold lengths*** | | ***7 nt*** | ***5 nt*** | ***3 nt*** | ***2 nt*** | ***1 nt*** | ***0 nt*** | |

**Supplementary Table 14| Estimated $\Delta G_{hb}$ and $k_d$ for each handhold at 37 ˚C**

Data obtained as described in **Supplementary Notes II and XVI.** The reactant strands used were all *N* strands "Incumbent x hh/x sp" except the ones with spacer 1,3 and 5 nt (**Supplementary Table 20**) with "Target (x th)" with 0, 2 and 8 nt (**Supplementary Table 21**) and "Invader ($I_{hc}$)" (**Supplementary Table 22**). Reporter strands were "Reporter 1 (*R*)" and "Reporter 1 quencher (*Q*)" (**Supplementary Table 23**).

| ***Condition (handhold/spacer)*** | **ΔGhb** *(Kcal M$^{-1}$)* | ***$k_{d\_init}$*** *($s^{-1}$)* | | ***Fitted $k_d$*** *($s^{-1}$)* | | ***Mean $k_d$ ± SEM*** *($s^{-1}$)* |
|---|---|---|---|---|---|---|
| **4/2** | -6.02 | 5.72E+02 | - | - | 1.95E+02 | 1.95E+02 |
| **5/0** | -6.26 | 3.88E+02 | 3.37E+02 | 1.82E+02 | 1.28E+02 | 2.16±0.51E+02 |
| **5/2** | -6.87 | 1.44E+02 | - | - | 5.01E+01 | 5.01E+01 |
| **5/8** | -6.84 | 1.51E+02 | 1.19E+02 | 3.59E+01 | 4.79E+01 | 6.8±2.1E+01 |
| **6/2** | -8.53 | 9.75E+00 | - | - | 9.13E+00 | 9.13E+00 |
| **7/2** | -9.45 | 2.19E+00 | - | - | 3.8E+00 | 3.8E+00 |
| **8/2** | -10.22 | 6.28E-01 | - | - | 1.05E+00 | 1.05E+00 |
| **9/2** | -12.49 | 1.59E-02 | - | - | 1.37E-02 | 1.37E-02 |
| **10/2** | -13.34 | 3.98E-03 | - | - | 1.53E-03 | 1.53E-03 |
| **20/0** | -24.65 | 4.26E-11 | 4.36E-10 | 1.04E-10 | 1.2E-11 | 1.8±1.1E-10 |
| **20/2** | -25.26 | 1.58E-11 | - | - | 1.6E-11 | 1.6E-11 |
| **20/8** | -25.23 | 1.66E-11 | 6.98E-11 | 1.52E-06 | 8.7E-12 | 5.1±4.14E-07 |
| | ***Toehold lengths*** | | ***7 nt*** | ***2 nt*** | ***0 nt*** | |

**Supplementary Table 15| Handhold Characterisation at 25 ˚C**

Data used for **Figure 2** in the main text. All the relevant characterisation constants are accompanied by the standard error of the mean of individual fittings (**Supplementary Note XVI**). Data obtained as described in **Supplementary Notes II, XIV (to obtain $k_t$) and XVI.** The reactant strands used were all *N* strands "Incumbent x hh/2 sp" (**Supplementary Table 20**) with all "Target (x th)" (**Supplementary Table 21**) and *I* strands "Invader ($I_{hc}$)", and to obtain the value of $k_t$ "Invader random-5' ($I_{wc}$)" (**Supplementary Table 22**). Reporter strands were "Reporter 1 (*R*)" and "Reporter 1 quencher (*Q*)" (**Supplementary Table 23**). Legend: **Bold:** Relevant reaction rate for the system. Yellow highlight: 1st order reaction rates. Green: $k_b$ with a negligible standard error of the mean. Artefact produced by the boundaries imposed to $k_b$ during its fitting. Red: Rates with a standard error bigger than the 10% of the mean. Corresponds to long experimental conditions (toehold=0 nt) or systems at the boundary between different kinetic regimes. Red highlight: $k_t$ values that were not obtained from direct experimental evidence but from the mean of all the systems with the same toehold length. Experiments omitted to reduce the number of extremely slow experiments. Standard error of the mean for each parameter was calculated from the values obtained in individual fittings.

| ***System hh/sp/toe*** | $k_m$ *($s^{-1}$)* | $k_d$ *($s^{-1}$)* | $k_b$ *($s^{-1}$ $M^{-1}$)* | $k_t$ *($s^{-1}$ $M^{-1}$)* | ***2nd order rate (k) (eq.3)*** *($s^{-1}$ $M^{-1}$)* | $t_{1/2}^{wc}$ *(min)* | $t_{1/2}$ *(min)* |
|---|---|---|---|---|---|---|---|
| **0/2/7** | 0 | 0 | 0 | **2.942±0.085E+6** | 2.942±0.085E+6 | 5.45E-01 | 5.45E-01 |
| **0/2/5** | 0 | 0 | 0 | **2.101±0.037E+5** | 2.101±0.037E+5 | 6.56E+00 | 6.56E+00 |
| **0/2/3** | 0 | 0 | 0 | **1.267±0.025E+4** | 1.267±0.025E+4 | 1.08E+02 | 1.08E+02 |
| **0/2/2** | 0 | 0 | 0 | **1.65±0.15E+2** | 1.65±0.15E+2 | 9.63E+03 | 9.63E+03 |
| **0/2/1** | 0 | 0 | 0 | **8.59±0.26E+1** | 8.59±0.26E+1 | 2.84E+04 | 2.84E+04 |
| **4/2/7** | 1.12E+0 | 8.19E+00 | 6.41E+06 | 1.053±0.007E+6 | **1.823±0.052E+6** | 1.77E+00 | 7.80E-01 |
| **4/2/5** | 9.52E-1 | 1.26E+01 | 5.24E+06 | 1.136±0.009E+5 | **4.82±0.11E+5** | 1.23E+01 | 2.93E+00 |
| **4/2/3** | 3.37E-1 | 1.32E+01 | 4.51E+06 | 1.599±0.011E+4 | **1.28±0.017E+5** | 8.77E+01 | 1.10E+01 |
| **4/2/2** | 1.34E-2 | 2.42E+01 | 5.72E+06 | 1.884±0.015E+2 | **3.357±0.066E+3** | 7.44E+03 | 4.19E+02 |
| **4/2/1** | 1.13E-2 | 2.79E+01 | 4.62E+06 | 1.097±0.045E+2 | **1.984±0.076E+3** | 1.28E+04 | 7.08E+02 |
| **4/2/0** | 1.78E-5 | 7.55E+00 | 4.76E+06 | 8.3±1.3E+0 | **1.95±0.41E+1** | 1.70E+05 | 7.31E+04 |
| **5/2/7** | 1.79E+0 | 8.57E+00 | 3.87E+06 | 6.123±0.093E+5 | **1.282±0.025E+6** | 2.29E+00 | 1.10E+00 |
| **5/2/5** | 4.18E-2 | 3.23E-01 | 3.87E+06 | 8.69±0.16E+4 | **4.958±0.068E+5** | 1.61E+01 | 2.85E+00 |
| **5/2/3** | 4.27E-1 | 8.46E+00 | 5.24E+06 | 1.674±0.008E+4 | **2.723±0.043E+5** | 1.823±0 .052E+6 | 5.29E+00 |
| **5/2/2** | 9.66E-3 | 7.28E+00 | 3.87E+06 | 2.354±0.048E+2 | **5.348±0.027E+3** | 4.82±0. 11E+5 | 2.63E+02 |
| **5/2/1** | 4.08E-3 | 8.54E+00 | 6.41E+06 | 1.116±0.006E+2 | **3.18±0.1E+3** | 1.26E+04 | 4.45E+02 |
| **5/2/0** | 1.49E-5 | 2.11E+00 | 5.33E+06 | 9.36±0.62E+0 | **4.607±0.064E+1** | 1.52E+05 | 3.13E+04 |
| **6/2/7** | 6.62E-1 | 5.14E-01 | 3.93E+6 | 1.072±0.021E+6 | **3.285±0.06E+6** | 1.31E+00 | 4.55E-01 |
| **6/2/5** | 1.55E-1 | 1.56E-01 | 3.76E+6 | 1.066±0.011E+5 | **1.98±0.21E+6** | 1.32E+01 | 8.45E-01 |
| **6/2/3** | 3.75E-2 | 2.87E-02 | 3.76E+6 | 2.135±0.015E+4 | **2.152±0.078E+6** | 6.57E+01 | 1.12E+00 |
| **6/2/2** | 2.70E-3 | 2.07E-01 | 3.77E+6 | 2.641±0.029E+2 | **4.87±0.12E+4** | 5.31E+03 | 3.66E+01 |
| **6/2/1** | 4.92E-3 | 8.92E-01 | 4.04E+6 | 1.335±0.01E+2 | **2.227±0.037E+4** | 1.05E+04 | 6.74E+01 |
| **6/2/0** | 7.18E-6 | 7.95E-02 | 4.53E+6 | 9.36±0.62E+0 | **4.19±0.18E+2** | 1.52E+05 | 6.03E+03 |
| **7/2/7** | 8.16E-2 | 1.63E-02 | 3.76E+6 | 9.99±0.17E+5 | **4.13±0.13E+6** | 1.40E+00 | 5.17E-01 |
| **7/2/5** | 1.92E-1 | 5.90E-02 | 3.76E+6 | 1.146±0.009E+5 | **2.991±0.031E+6** | 1.22E+01 | 5.71E-01 |
| **7/2/3** | 1.23E-1 | 9.77E-02 | 6.29E+6 | 2.433±0.032E+4 | **3.53±0.11E+6** | 5.76E+01 | 5.55E-01 |
| **7/2/2** | 5.63E-3 | 4.72E-02 | 3.76E+6 | 2.987±0.045E+2 | **4±1.3E+5** | 4.69E+03 | 6.87E+00 |
| **7/2/1** | 7.75E-4 | 2.07E-03 | 6.28E+6 | 1.247±0.006E+2 | **1.71±0.11E+6** | 1.12E+04 | 1.74E+01 |
| **7/2/0** | 4.09E-5 | 8.26E-02 | 3.76E+6 | 9.36±0.62E+0 | **1.87±0.12E+3** | 1.52E+05 | 1.24E+03 |
| **8/2/7** | 7.27E-2 | 1.60E-02 | 3.76E+6 | 8.548±0.09E+5 | **3.94±0.14E+6** | 1.64E+00 | 5.59E-01 |
| **8/2/5** | 7.79E-2 | 9.74E-03 | 3.76E+6 | 1.11±0.007E+5 | **3.45±0.1E+6** | 1.26E+01 | 6.41E-01 |
| **8/2/3** | 4.65E+0 | 7.65E-02 | 3.81E+6 | 2.281±0.018E+4 | **3.768±0.047E+6** | 6.15E+01 | 3.88E-01 |
| **8/2/2** | 3.78E-2 | 1.23E-01 | 6.29E+6 | 2.52±0.032E+2 | **1.478±0.044E+6** | 5.56E+03 | 1.47E+00 |
| **8/2/1** | 5.58E-3 | 1.76E-02 | 6.29E+6 | 1.28±0.017E+2 | **1.51±0.39E+6** | 1.10E+04 | 3.68E+00 |
| **8/2/0** | 4.84E-5 | 2.63E-04 | 6.29E+6 | 9.36±0.62E+0 | **9.78±0.83E+5** | 1.52E+05 | 2.70E+02 |
| **9/2/7** | 2.01E+0 | 8.88E-14 | 4.04±0.16E+6 | 2.384±0.069E+6 | **6.42±0.3E+6** | 5.88E-01 | 2.27E-01 |
| **9/2/5** | 3.90E+0 | 2.37E-11 | 3.828±0.048E+6 | 2.529±0.038E+5 | **4.08±0.056E+6** | 5.54E+00 | 3.51E-01 |
| **9/2/3** | 1.99E+0 | 9.01E-09 | 4.92±0.1E+6 | 6.137±0.024E+4 | **4.99±0.1E+6** | 2.28E+01 | 2.93E-01 |
| **9/2/2** | 4.01E+0 | 6.09E-09 | 5.29±0.34E+6 | 4.66±0.89E+2 | **5.29±0.34E+6** | 3.03E+03 | 2.73E-01 |
| **9/2/1** | 6.57E+1 | 3.73E-09 | 4.405±0.084E+6 | 1.69±0.36E+2 | **4.406±0.084E+6** | 8.28E+03 | 3.22E-01 |
| **9/2/0** | **6.007±0.021E-4** | 4.08E-06 | 6.29E+6 | 9.36±0.62E+0 | 6.29E+6 | 1.52E+05 | 1.96E+01 |

| | | | | | | | |
|---|---|---|---|---|---|---|---|
| **10/2/7** | 1.26E+0 | 2.89E-12 | 3.76E+6 | 1.586±0.049E+6 | **5.347±0.098E+6** | 8.84E-01 | 2.75E-01 |
| **10/2/5** | 6.83E+1 | 5.80E-03 | 3.95±0.18E+6 | 2.417±0.034E+5 | **4.2±0.19E+6** | 5.80E+00 | 3.39E-01 |
| **10/2/3** | 1.83E+0 | 6.13E-03 | 4.41±0.14E+6 | 5.054±0.009E+4 | **4.44±0.14E+6** | 2.77E+01 | 3.29E-01 |
| **10/2/2** | 3.85E-2 | 3.79E-03 | 4.5±0.1E+6 | 6.97±0.058E+2 | **4.1±0.1E+6** | 2.01E+03 | 8.00E-01 |
| **10/2/1** | 8.53E+1 | 4.90E-04 | 3.76E+6 | 3.584±0.02E+2 | **3.76E+6** | 3.91E+03 | 3.76E-01 |
| **10/2/0** | **8.85±0.25E-4** | 1.52E-03 | 3.76E+6 | 9.36±0.62E+0 | 3.76E+6 | 1.52E+05 | 1.54E+01 |
| **20/2/7** | 1.09E+1 | 2.39E-16 | 5.7±0.11E+6 | 1.277±0.024E+6 | **6.98±0.16E+6** | 1.10E+00 | 2.06E-01 |
| **20/2/5** | 5.16E+1 | 6.68E-16 | 6.29E+6 | 2.85±0.41E+5 | **6.579±0.071E+6** | 4.92E+00 | 2.17E-01 |
| **20/2/3** | 2.08E+0 | 1.46E-16 | 4.75±0.11E+6 | 5.201±0.045E+4 | **4.8±0.11E+6** | 2.70E+01 | 3.04E-01 |
| **20/2/2** | 4.68E+0 | 5.41E-16 | 5.412±0.072E+6 | 8.64±0.1E+2 | **5.413±0.072E+6** | 1.63E+03 | 2.66E-01 |
| **20/2/1** | 2.34E+0 | 8.03E-17 | 3.761±0.012E+6 | 3.03±0.16E+2 | **3.762±0.012E+6** | 4.63E+03 | 3.83E-01 |
| **20/2/0** | **2.038±0.015E-3** | 8.14E-17 | 6.29E+6 | 9.36±0.62E+0 | 6.29E+6 | 1.52E+05 | 6.09E+00 |

**Supplementary Table 16| Spacer Characterisation at 25 ˚C**

Data used to infer the effect of a spacer sequence between the handhold and the displaced domain. All the relevant characterisation constants are accompanied by the standard error of the mean of individual fittings (**Supplementary Note XVI**). Data obtained as described in **Supplementary Notes II, XIV (to obtain $k_t$) and XVI.** The reactant strands used were all *N* strands "Incumbent x hh/x sp" excluding 2 nt spacers (**Supplementary Table 20**) with *T* strands "Target (x th)" with toeholds of 0, 2 and 8 nt (**Supplementary Table 21**) and *I* strands "Invader ($I_{hc}$)", and to obtain the value of $k_t$ "Invader random-5' ($I_{wc}$)" (**Supplementary Table 22**). Reporter strands were "Reporter 1 (*R*)" and "Reporter 1 quencher (*Q*)" (**Supplementary Table 23**). Legend: **Bold:** Relevant reaction rate for the system. Yellow highlight: 1st order reaction rates. Green: $k_b$ with a negligible standard error. Artefact produced by the boundaries imposed to $k_b$ during its fitting. Red: Rates with a standard error bigger than the 10% of the mean. Corresponds to long experimental conditions (toehold=0 nt) or systems at the boundary between different kinetic regimes. Red highlight: $k_t$ values that were not obtained from direct experimental evidence but from the mean of all the systems with the same toehold length. Experiments omitted to reduce the number of extremely slow experiments. Standard error of the mean for each parameter was calculated from the values obtained in individual fittings.

| *System hh/sp/toe* | $k_m$ ($s^{-1}$) | $k_d$ ($s^{-1}$) | $k_b$ ($s^{-1}$ $M^{-1}$) | $k_t$ ($s^{-1}$ $M^{-1}$) | *2nd order rate (k) (eq.3)* ($s^{-1}$ $M^{-1}$) | $t_{1/2}^{wc}$ (min) | $t_{1/2}$ (min) |
|---|---|---|---|---|---|---|---|
| **0/0/7** | 0 | 0 | 0 | **4.39±0.12E+6** | 4.39±0.12E+6 | 2.79E-01 | 2.82E-01 |
| **0/0/2** | 0 | 0 | 0 | **1.99±0.1E+2** | 1.99±0.1E+2 | 7.52E+03 | 7.51E+03 |
| **0/1/7** | 0 | 0 | 0 | **3.43±0.12E+6** | 3.43±0.12E+6 | 4.20E-01 | 4.23E-01 |
| **0/1/2** | 0 | 0 | 0 | **1.79±0.029E+2** | 1.79±0.029E+2 | 7.48E+03 | 7.47E+03 |
| **0/1/0** | 0 | 0 | 0 | **1.041±0.067E+1** | 1.041±0.067E+1 | 3.78E+05 | - |
| **0/3/7** | 0 | 0 | 0 | **2.274±0.057E+6** | 2.274±0.057E+6 | 5.74E-01 | 5.77E-01 |
| **0/3/2** | 0 | 0 | 0 | **1.37±0.12E+2** | 1.37±0.12E+2 | 9.08E+03 | 9.07E+03 |
| **0/5/7** | 0 | 0 | 0 | **5.503±0.095E+5** | 5.503±0.095E+5 | 2.34E+00 | 2.35E+00 |
| **0/5/2** | 0 | 0 | 0 | **1.407±0.094E+2** | 1.407±0.094E+2 | 9.16E+03 | 9.15E+03 |
| **0/8/7** | 0 | 0 | 0 | **3.25±0.11E+5** | 3.25±0.11E+5 | 3.83E+00 | 3.83E+00 |
| **0/8/2** | 0 | 0 | 0 | **1.093±0.068E+2** | 1.093±0.068E+2 | 1.32E+04 | 1.32E+04 |
| **5/0/7** | 1.46E-1 | 1.34E+01 | 6.25E+6 | 3.09±0.068E+5 | **3.76±0.17E+5** | 4.54E+00 | 3.76E+00 |
| **5/0/2** | 1.98E-2 | 2.11E+01 | 6.29E+6 | 3.112±0.043E+2 | **6.21±0.096E+3** | 4.55E+03 | 2.27E+02 |
| **5/0/0** | 1.31E-4 | 7.67E+00 | 5.73E+6 | 9.36±0.62E+0 | **1.069±0.048E+2** | 1.52E+05 | 1.33E+04 |
| **5/1/7** | 8.91E-2 | 2.11E+00 | 3.76E+6 | 4.764±0.07E+5 | **6.29±0.17E+5** | 2.94E+00 | 2.29E+00 |
| **5/1/2** | 2.28E-1 | 7.80E+02 | 5.12E+6 | 2.758±0.028E+2 | **1.774±0.023E+3** | 5.04E+03 | 7.90E+02 |
| **5/1/0** | 1.82E-6 | 2.57E+00 | 4.51E+6 | 9.36±0.62E+0 | **1.24±0.13E+1** | 1.52E+05 | 1.16E+05 |
| **5/3/7** | 3.27E-1 | 2.63E+00 | 4.16E+6 | 9.13±0.16E+5 | **1.373±0.054E+6** | 1.54E+00 | 1.05E+00 |
| **5/3/2** | 1.19E-3 | 9.16E+00 | 4.03E+6 | 3.855±0.029E+2 | **9.12±0.11E+2** | 3.62E+03 | 1.55E+03 |
| **5/3/0** | 5.96E-7 | 9.56E+00 | 3.76E+6 | 9.36±0.62E+0 | **9.5±1.4E+0** | 1.52E+05 | 1.49E+05 |
| **5/5/7** | 2.12E-1 | 6E+00 | 3.76E+6 | 3.64±0.14E+5 | **4.92±0.19E+5** | 3.85E+00 | 2.88E+00 |
| **5/5/2** | 1.80E-4 | 7.62E+00 | 6.22E+6 | 2.66±0.025E+2 | **4.14±0.11E+2** | 5.27E+03 | 3.43E+03 |
| **5/5/0** | 1.76E-9 | 0.14E+00 | 6.16E+6 | 9.36±0.62E+0 | **9.33±0.58E+0** | 1.52E+05 | 1.86E+05 |
| **5/8/7** | 2.07E-1 | 2.59E+00 | 5.40E+6 | 4.23±0.11E+5 | **8.22±0.49E+5** | 3.32E+00 | 1.76E+00 |
| **5/8/2** | 1.78E-4 | 4.7E+00 | 6.29E+6 | 1.46±0.12E+2 | **4.05±0.23E+2** | 8.43E+03 | 3.53E+03 |
| **5/8/0** | 6.19E-7 | 2.79E+00 | 6.18E+6 | 9.36±0.62E+0 | **1.062±0.058E+1** | 1.52E+05 | 1.36E+05 |
| **20/0/7** | 1.46E+1 | 1.04E-20 | 6.29±0.11E+6 | 6.284±0.085E+5 | **6.92±0.12E+6** | 2.23E+00 | 2.08E-01 |
| **20/0/2** | 1.03E+1 | 6.59E-16 | 3.76E+6 | 1.43±0.044E+3 | **3.76E+6** | 1.02E+03 | 3.78E-01 |
| **20/0/0** | **4.015±0.033E-4** | 3.81E-16 | 6.29E+6 | 9.36±0.62E+0 | 6.29E+6 | 1.52E+05 | 2.92E+01 |
| **20/1/7** | 2.67E+0 | 3.01E-16 | 6.29E+6 | 1.34±0.02E+6 | **7.634±0.02E+6** | 1.05E+00 | 1.92E-01 |
| **20/1/2** | 2.02E+0 | 3.72E-16 | 5.41±0.13E+6 | 1.436±0.012E+3 | **5.41±0.13E+6** | 9.81E+02 | 2.71E-01 |
| **20/1/0** | **2.62±0.11E-3** | 1.17E-16 | 6.29E+6 | 9.36±0.62E+0 | 6.29E+6 | 1.52E+05 | 4.84E+00 |
| **20/3/7** | 1.39E+0 | 9.14E-17 | 4.8±0.43E+6 | 1.417±0.063E+6 | **6.21±0.49E+6** | 9.89E-01 | 2.38E-01 |
| **20/3/2** | 1.66E+1 | 4.1E-16 | 4.672±0.089E+6 | 1.36±0.014E+3 | **4.673±0.089E+6** | 1.03E+03 | 3.04E-01 |
| **20/3/0** | **1.28±0.02E-3** | 1.27E-16 | 5.46E+6 | 9.36±0.62E+0 | 5.46E+6 | 1.52E+05 | 9.50E+00 |
| **20/5/7** | 5.48E-1 | 1.69E-16 | 3.761±0.032E+6 | 5.291±0.033E+5 | **4.29±0.035E+6** | 2.65E+00 | 3.58E-01 |
| **20/5/2** | 1.27E-1 | 2.58E-16 | 4.52±0.26E+6 | 7.387±0.033E+2 | **4.53±0.26E+6** | 1.90E+03 | 4.62E-01 |
| **20/5/0** | **6.191±0.028E-4** | 1.67E-16 | 3.96E+6 | 9.41±0.73E+0 | 3.96E+6 | 1.52E+05 | 1.93E+01 |
| **20/8/7** | 1.74E+1 | 2.65E-16 | 4.047±0.045E+6 | 7.752±0.044E+5 | **4.822±0.049E+6** | 1.81E+00 | 2.95E-01 |
| **20/8/2** | 1.68E-1 | 5.73E-16 | 4.2±0.13E+6 | 6.429±0.045E+2 | **4.21±0.13E+6** | 2.21E+03 | 4.49E-01 |
| **20/8/0** | **5.686±0.03E-4** | 2.17E-16 | 3.76E+6 | 9.36±0.62E+0 | 3.76E+6 | 1.52E+05 | 2.10E+01 |

**Supplementary Table 17| Handhold Characterisation at 37 ˚C**

All the relevant characterisation constants are accompanied by the standard error of the mean between all the obtained fluorescence traces for that reaction system. Data obtained as described in **Supplementary Notes II, XIV (to obtain $k_t$) and XVI.** The reactant strands used were *N* strands "Incumbent x hh/x sp" with spacers 0, 2 and 8 (**Supplementary Table 20**) with *T* strands "Target (x th)" with toeholds of 0, 2 and 7 nt (**Supplementary Table 21**) and *I* strands "Invader ($I_{hc}$)", and to obtain the value of $k_t$ "Invader random-5' ($I_{wc}$)" (**Supplementary Table 22**). Reporter strands were "Reporter 1 (*R*)" and "Reporter 1 quencher (*Q*)" (**Supplementary Table 23**). Legend: **Bold:** Relevant reaction rate for the system. Yellow highlight: 1st order reaction rates. Red: Rates with a standard error bigger than the 10% of the mean. Corresponds with long experimental conditions (toehold=0 nt) or systems at the boundary between different kinetic regimes. Red highlight: $k_t$ values that were not obtained from direct experimental evidence. Their value was estimated from the $k_t$ for a toehold=0 nt at 25˚C. Standard error of the mean for each parameter was calculated from the values obtained in individual fittings.

| ***System hh/sp/toe*** | $k_m$ *($s^{-1}$)* | $k_d$ *($s^{-1}$)* | $k_b$ *($s^{-1}$ $M^{-1}$)* | $k_t$ *($s^{-1}$ $M^{-1}$)* | ***2nd order rate (k) (eq.3)*** *($s^{-1}$ $M^{-1}$)* | $t^{wc}_{1/2}$ *(min)* | $t_{1/2}$ *(min)* |
|---|---|---|---|---|---|---|---|
| **0/0/7** | 0 | 0 | 0 | **5.14±0.11E+6** | 5.14±0.11E+6 | 2.38E-01 | 2.41E-1 |
| **0/0/2** | 0 | 0 | 0 | **4.33±0.2E+2** | 4.24±0.19E+2 | 3.63E+03 | 3.63E+3 |
| **0/8/7** | 0 | 0 | 0 | **4.27±0.053E+5** | 4.27±0.053E+5 | 2.53E+00 | 2.53E+0 |
| **0/8/2** | 0 | 0 | 0 | **2.71±0.16E+2** | 2.65±0.16E+2 | 5.43E+03 | 5.42E+3 |
| **5/0/7** | 2.3E+0 | 3.37E+02 | 8.34E+06 | 3.746±0.048E+5 | **4.312±0.127E+5** | 3.74E+00 | 3.26E+0 |
| **5/0/2** | 3.61E-2 | 1.82E+02 | 7.66E+06 | 4.32±0.17E+2 | **1.955±0.187E+3** | 3.31E+03 | 7.18E+2 |
| **5/0/0** | 1.22E-3 | 1.28E+02 | 5.73E+06 | 9.36±0.62E+0 | **6.419±0.073E+1** | 1.52E+05 | 2.18E+4 |
| **5/8/7** | 1.82E+0 | 1.19E+02 | 1.04E+07 | 4.915±0.08E+5 | **6.472±0.133E+5** | 2.85E+00 | 2.17E+0 |
| **5/8/2** | 2.85E-4 | 3.59E+01 | 5.57E+06 | 3.4±0.19E+2 | **3.841±0.284E+2** | 4.21E+03 | 3.66E+3 |
| **5/8/0** | 4.36E-5 | 4.80E+01 | 7.97E+06 | 9.36±0.62E+0 | **1.661±0.123E+1** | 1.52E+05 | 8.45E+4 |
| **20/0/7** | 1.24E+1 | 4.36E-10 | **1.459±0.003E+7** | 6.424±0.083E+5 | **1.5229±0.003E+7** | 2.18E+00 | 9.82E-2 |
| **20/0/2** | 7.93E+0 | 1.04E-10 | **1.054±0.025E+7** | 5.33±1E+2 | **1.054±0.025E+7** | 2.76E+03 | 1.40E-1 |
| **20/0/0** | **6.456±0.081E-4** | 1.18E-11 | 1.19E+07 | 9.36±0.62E+0 | 1.19E+7 | 1.52E+05 | 1.81E+1 |
| **20/8/7** | 6.37E+0 | 6.98E-11 | **5.57±0.27E+6** | 6.187±0.078E+5 | **6.19±0.28E+6** | 2.27E+00 | 2.32E-1 |
| **20/8/2** | 1.31E+1 | 1.52E-06 | **8.59±0.75E+6** | 3.77±0.4E+2 | **8.59±0.75E+6** | 3.79E+03 | 1.68E-1 |
| **20/8/0** | **1.323±0.008E-3** | 8.75E-12 | 1.44E+07 | 9.36±0.62E+0 | **1.44E+7** | 1.52E+05 | 8.9E+0 |
| **4/2/0** | 3.58E-4 | 1.95E+02 | 8.67E+06 | 9.36±0.62E+0 | **2.527±0.104E+1** | 5.58E+04 | 5.55E+4 |
| **5/2/0** | 1.39E-4 | 5.01E+01 | 7.66E+06 | 9.36±0.62E+0 | **3.798±0.11E+1** | 3.71E+04 | 3.70E+4 |
| **6/2/0** | 1.41E-4 | 9.13E+00 | 6.41E+06 | 9.36±0.62E+0 | **1.08±0.11E+2** | 1.31E+04 | 1.31E+4 |
| **7/2/0** | 1.78E-4 | 3.80E+00 | 7.07E+06 | 9.36±0.62E+0 | **3.395±0.099E+2** | 4.35E+03 | 4.25E+3 |
| **8/2/0** | 2.80E-4 | 1.06E+00 | 5.57E+06 | 9.36±0.62E+0 | **1.486±0.031E+3** | 1.02E+03 | 1.02E+3 |
| **9/2/0** | **1.85±0.17E-4** | 1.37E-02 | 9.02E+06 | 9.36±0.62E+0 | 1.2E+5 | 8.20E+01 | 9.73E+1 |
| **10/2/0** | **5.65±0.36E-4** | 1.53E-03 | 9.60E+06 | 9.36±0.62E+0 | 2.59E+6 | 2.27E+01 | 2.27E+1 |
| **20/2/0** | **1.78±0.28E-3** | 1.60E-11 | 5.57E+06 | 9.36±0.62E+0 | 5.57E+6 | 7.02E+00 | 6.96E+0 |

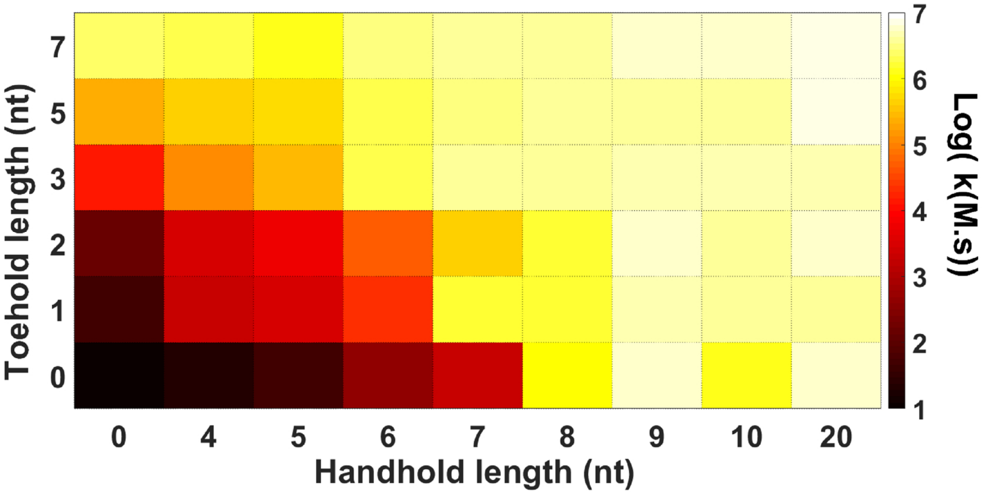

**Supplementary Figure 17| Inferred 2nd order Rates for handhold-mediated strand displacement.**
*k* values collected in **Supplementary Table 15.**

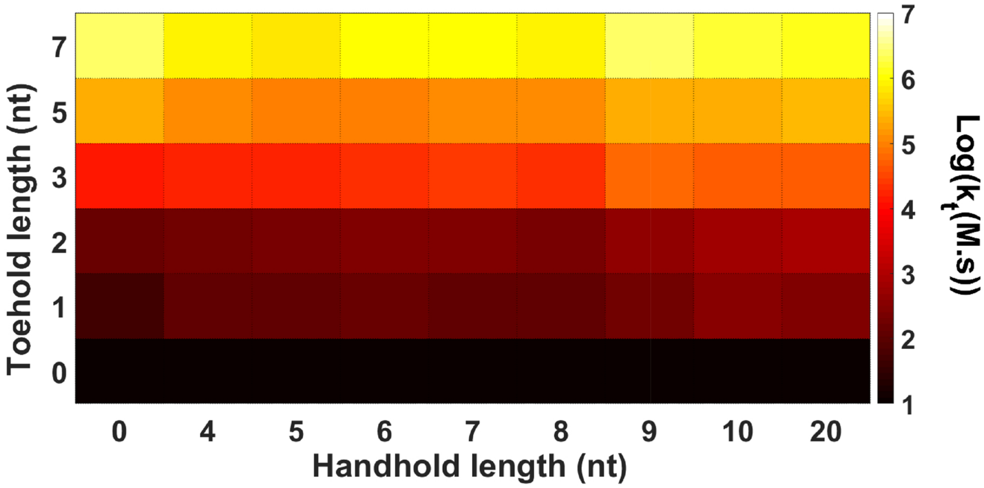

**Supplementary Figure 18| Rates for *I*-strands without handhold complementarity.**
The reaction rates are the ones of a toehold mediated strand displacement ($k_t$, **Supplementary Table 15**). Handhold presence in *N* produces negligible rate changes for strand displacement reactions when *I*-strands lack handhold-complementarity.

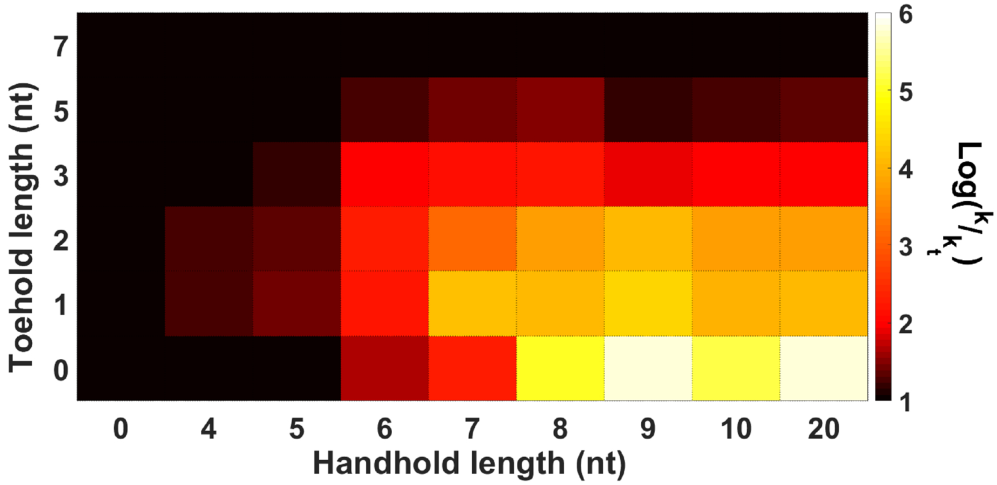

**Supplementary Figure 19| Comparison between *k* and $k_t$**
Our inferred second order rate constants suggest that HMSD can increase displacement rate by up to 5 orders of magnitude compared to displacement triggered with *I*-strands without handhold-complementarity. The reaction rate increment obtained by the addition of handhold-complementarity is sensibly higher in the regime of toeholds below 3 nt.

#### Supplementary Note XVII: Fitting of the kinetic analysis of handhold detachment

$R_2Q_2$ controls shown in **Supplementary Note IX** demonstrated that the presence of strand *T* in the solution triggered the associative toehold at a slow rate, except when free in solution as a product *IT*. Due to the high rate of reaction of $R_2Q_2$ with the product *IT*, the slow-rate leak reactions were ignored when fitting the kinetics of $R_2Q_2$ .The obtained trajectories were fitted to obtain the value of $K_{eq}$ (**equation 5**). Each trajectory was fit individually to the ODE describing **equation 4**:

$$\frac{d[R_2IT]}{dt} = k_{r2}[R_2Q][IT]\,, \quad (28)$$

With constraints:

$$[R_2Q_2]=[R_2Q_2]_0-[R_2IT]\,, \quad (29)$$

$$[IT]=\frac{-(K_{eq}+[N]_0+[R_2IT]-[I]_0)+\sqrt{([N]_0+K_{eq}+[R_2IT]-[I]_0)^2-4K_{eq}([R_2IT]-[I]_0)}}{2}\,. \quad (30)$$

**Supplementary equation 30** describes free *IT* assuming an instantaneous equilibrium between *N/IT* and *IT+N* (equation 5). The detailed derivation of **Supplementary equation 30** from the **equation 5** of the main text is included in the **Supplementary Note XIX**. The values of $k_{r2}$ were obtained from individually fitting Supplementary equations 24 and 25 to the results for a system in the absence of handhold (**Supplementary Figure 20**)

For each system, its fluorescence traces were fitted individually to obtain the value of $K_{eq}$, $[I]_0$ and $t_0$. The value for $K_{eq\_init}$ was set arbitrarily to $5.10^{-9}$ nM. For the obtained $K_{eq}$, the percentage of expected detached [*IT*] in the equilibrium was calculated with **Supplementary equation 30** for $[N]_0$;$[I]_0$=10nM and $[R_2IT]$=0 nM. Fitted $K_{eq}$ and the expected percentage of detached [*IT*] are collected in **Supplementary Table 18.** Results for the control without *RQ* reporter are collected in **Supplementary Table 19.**

**Supplementary Table 18| Fitting results from the kinetic analysis of handhold detachment**

Data obtained as described in **Supplementary Notes III and XVII.** The reactant strands used were all *N* strands "Incumbent x hh/2 sp" (**Supplementary Table 20**) with "Target(3 th)-$R_2Q_2$ trigger" (**Supplementary Table 21**) and "Invader ($I_{hc}$)" (**Supplementary Table 22**). Reporter strands were "Reporter 1 (*R*)", "Reporter 1 quencher (*Q*)", "Rep$_A$ ($R_2$)" and "Rep$_A$ quencher ($Q_2$)"(**Supplementary Table 23**).

| ***Condition (handhold/spacer)*** | ***$K_{eq}$ (M)*** | ***% of detached displacement products*** |
|---|---|---|
| **4/2** | 1.27E-1 | 99.99 |
| **5/2** | 1.19E-1 | 99.99 |
| **6/2** | 9E-1 | 99.99 |
| **7/2** | 1.01E-1 | 99.99 |
| **8/2** | 2.1E-1 | 99.99 |
| **9/2** | 5.6E-8 | 83.3±2.8 |
| **10/2** | 3.945E-9 | 45.6±1.7 |
| **20/2** | ~0 | 0 |

**Supplementary Table 19| Fitting results from the kinetic analysis of handhold detachment in the absence of reporter *RQ***

Data obtained as described in **Supplementary Notes III and XVII.** The reactant strands used were *N* strands "Incumbent x hh/2 sp" with handholds 7, 8, 9 and 10 nt (**Supplementary Table 20**) with "Target(3 th)-$R_2Q_2$ trigger" (**Supplementary Table 21**) and "Invader ($I_{hc}$)" (**Supplementary Table 22**). Reporter strands were "$Rep_A$ ($R_2$)" and "$Rep_A$ quencher ($Q_2$)"(**Supplementary Table 23).** Standard error of the mean for each parameter was calculated from the values obtained in individual fittings.

| *Condition (handhold/spacer)* | *$K_{eq}$ ± SEM (M)* | *% of detached displacement products* |
|---|---|---|
| **7/2** | 1.01E-1 | 99.99 |
| **8/2** | 1.77E-1 | 99.99 |
| **9/2** | 2.381±0.02E-8 | 75.84±0.14 |
| **10/2** | 5.22±0.23E-9 | 50.68±0.82 |

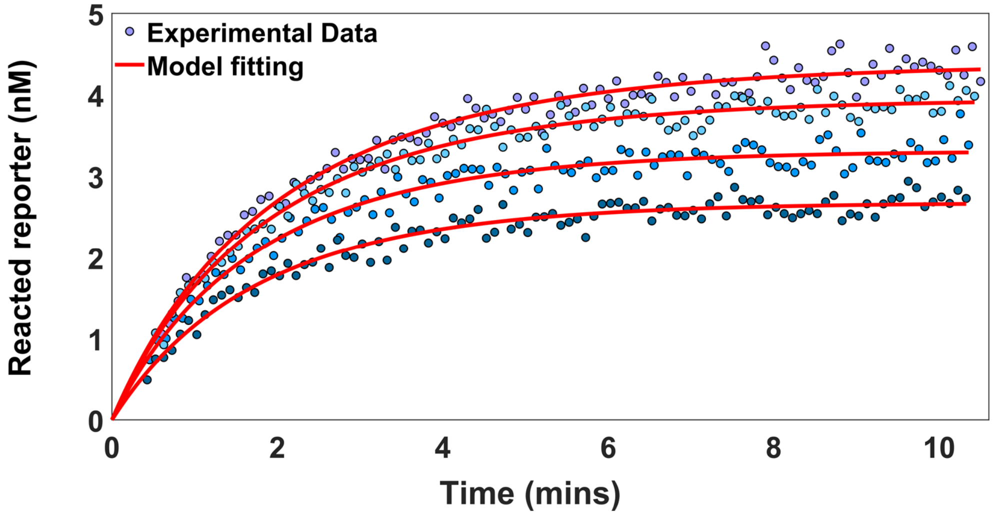

**Supplementary Figure 20| Triggering of reporter $R_2Q_2$ in the absence of handhold.**

Processed fluorescence data of the addition of $R_2Q_2$ (Measurement 4) on a solution with a system without handhold, 2 nt spacer, and a 3 nt toehold that has reacted with $I_{hc}$. Each trajectory was fitted individually with **equations 28 and 29** to obtain the value of $k_{r2}$. In the absence of handhold all *IT* was assumed to be detached. The obtained values for $k_{r2}$ were 6.51, 7.14, 7.58 and 7.33.10$^5$ M$^{-1}$s$^{-1}$, which produces a mean of 7.14±0.23.10$^5$ M$^{-1}$s$^{-1}$. All concentrations are defined for a reaction volume of 200 µL.

#### Supplementary Note XVIII: Derivation of HMSD bimolecular rate (equation 3)

To calculate the effective rate of *IT* production, we omit the reporter reaction and assume the species *TN/I* to be in quasi-steady state, since $k_d$ and $k_m$ are assumed to be large:

$$\frac{d[TN/I]}{dt} = k_b[TN][I] - k_d[TN/I] - k_m[TN/I] = 0$$

$$[TN/I] = \frac{k_b}{k_d + k_m}\,[TN][I] \quad (31)$$

So, the rate of production of *IT* for handhold mediated strand displacement is defined :

$$\frac{d[IT]}{dt} = k_m\,[TN/I] = k_m\,\frac{k_b}{k_d+k_m}\,[TN][I] \quad (32)$$

Toehold-mediated strand displacement also contributes to the reaction with a rate equal to $k_t[TN][I]$. By adding the contribution of handhold and toehold mediated displacements we obtain the final rate:

$$\frac{d[IT]}{dt} = \left(k_m\,\frac{k_b}{k_d+k_m} + k_t\right)[TN][I] = k\,[TN][I] \quad (33)$$

#### Supplementary Note XIX: Derivation of Supplementary equation 30

From the equilibrium equation (**equation 5**), it is possible to obtain a boundary for the value of [*IT*] depending on the concentrations of the species in the solution. The first step is to substitute $[N]=[N]_0-[N/IT]$ and $[N/IT]=[I]_0-[IT]-[R_2IT]$:

$$K_{eq}=\frac{[N][IT]}{[N/IT]}=\frac{([N]_0+[IT]+[R_2IT]-[I]_0)[IT]}{[I]_0-[IT]-[R_2IT]},\quad (34)$$

$$([I]^0-[IT]-[R_2IT])K_{eq}=([N]_0+[IT]+[R_2IT]-[I]_0)[IT]\ ,$$

$$K_{eq}([I]^0-[R_2IT])-K_{eq}[IT]=[IT]^2+([N]_0+[R_2IT]-[I]_0)[IT]\ ,$$

$$0=[IT]^2+(K_{eq}+[N]_0+[R_2IT]-[I]_0)[IT]+K_{eq}([R_2IT]-[I]_0)\ (35)\ ,$$

[*IT*] (**Supplementary equation 30**) is the positive solution of the quadratic equation:

$$[IT]=\frac{-(K_{eq}+[N]_0+[R_2IT]-[I]_0)+\sqrt{(K_{eq}+[N]_0+[R_2IT]-[I]_0)^2-4K_{eq}([R_2IT]-[I]_0)}}{2}$$

#### Supplementary Note XX:Sequences

All sequences used during the experiments are collected in **Supplementary Table 20 to 23.** Strands are named by their handhold (hh), spacer (sp) and toehold (th) sequences length. The combination of these strands produces the complexes referred during the supplementary as "X/X/X_Temperature". For example, the combination of Incumbent(5hh/2sp) with Target(7nt) reacting at 25˚C, is referred as the system 5/2/7_25. The domains of each strand are coded in the same colour as in **Figures 1 and 3,** specifically:

Grey: *R* binding domain in *N* strands and binding domain between *R* and *Q* strands.

Orange: *N* strands domain. Displaced domain during the strand displacement. Same sequence as green.

Green: *I* strands domain. Same sequence as orange.

Blue: *T* strands domain. Complementary to orange/green

Gold: Handhold domain of variable length in *N* strands. Variable 8 nt sequence in reporters. Handhold-complementary sequence in *I* strands.

Light blue: Toehold domain. Always same sequence, but variable length in *T* strands. 7 nt for *I* strands and 2 nt for $I_{A/B/C}$ strands

Black: Spacer domains between orange and gold domains in *N.* Hairpins, poly-T spacers and clamps in $Rep_{A/B/C}$ strands.

Purple: $Rep_{A/B/C}$ binding domain in *N* strands and binding domain between $Rep_{A/B/C}$ and their quenchers.

**Supplementary Table 20 | Incumbent strands (*N*)**

| | Sequence | Length (nt) |
|---|---|---|
| **Incumbent (0 hh/0 sp)** | 5′ – CCC TAT TAC CCT TGC CGA CCC TAC CTA CCT ACC CTG CG – 3′ | 38 |
| **Incumbent (0 hh/1 sp)** | 5′ – CCC TAT TAC CCT TGC CGA CCC TAC CTA CCT ACC CTG CGA – 3′ | 39 |
| **Incumbent (0 hh/2 sp)** | 5′ – CCC TAT TAC CCT TGC CGA CCC TAC CTA CCT ACC CTG CGA T – 3′ | 40 |
| **Incumbent (0 hh/3 sp)** | 5′ – CCC TAT TAC CCT TGC CGA CCC TAC CTA CCT ACC CTG CGA TA – 3′ | 41 |
| **Incumbent (0 hh/5 sp)** | 5′ – CCC TAT TAC CCT TGC CGA CCC TAC CTA CCT ACC CTG CGA TAC C – 3′ | 43 |
| **Incumbent (0 hh/8 sp)** | 5′ – CCC TAT TAC CCT TGC CGA CCC TAC CTA CCT ACC CTG CGA TAC CTA A – 3′ | 46 |
| **Incumbent (4 hh/2 sp)** | 5′ – CCC TAT TAC CCT TGC CGA CCC TAC CTA CCT ACC CTG CGA TGA TG – 3′ | 44 |
| **Incumbent (5 hh/0 sp)** | 5′ – CCC TAT TAC CCT TGC CGA CCC TAC CTA CCT ACC CTG CGG ATG T – 3′ | 43 |
| **Incumbent (5 hh/1 sp)** | 5′ – CCC TAT TAC CCT TGC CGA CCC TAC CTA CCT ACC CTG CGA GAT GT – 3′ | 44 |
| **Incumbent (5 hh/2 sp)** | 5′ – CCC TAT TAC CCT TGC CGA CCC TAC CTA CCT ACC CTG CGA TGA TGT – 3′ | 45 |
| **Incumbent (5 hh/3 sp)** | 5′ – CCC TAT TAC CCT TGC CGA CCC TAC CTA CCT ACC CTG CGA TAG ATG T – 3′ | 46 |
| **Incumbent (5 hh/5 sp)** | 5′ – CCC TAT TAC CCT TGC CGA CCC TAC CTA CCT ACC CTG CGA TAC CGA TGT – 3′ | 48 |
| **Incumbent (5 hh/8 sp)** | 5′ – CCC TAT TAC CCT TGC CGA CCC TAC CTA CCT ACC CTG CGA TAC CTA AGA TGT – 3′ | 51 |
| **Incumbent (6 hh/2 sp)** | 5′ – CCC TAT TAC CCT TGC CGA CCC TAC CTA CCT ACC CTG CGA TGA TGT G – 3′ | 46 |
| **Incumbent (7 hh/2 sp)** | 5′ – CCC TAT TAC CCT TGC CGA CCC TAC CTA CCT ACC CTG CGA TGA TGT GA – 3′ | 47 |
| **Incumbent (8 hh/2 sp)** | 5′ – CCC TAT TAC CCT TGC CGA CCC TAC CTA CCT ACC CTG CGA TGA TGT GAA – 3′ | 48 |
| **Incumbent (9 hh/2 sp)** | 5′ – CCC TAT TAC CCT TGC CGA CCC TAC CTA CCT ACC CTG CGA TGA TGT GAA G – 3′ | 49 |
| **Incumbent (10 hh/2 sp)** | 5′ – CCC TAT TAC CCT TGC CGA CCC TAC CTA CCT ACC CTG CGA TGA TGT GAA GT – 3′ | 50 |
| **Incumbent (20 hh/0 sp)** | 5′ – CCC TAT TAC CCT TGC CGA CCC TAC CTA CCT ACC CTG CGG ATG TGA AGT GAA TGA TGA G – 3′ | 58 |
| **Incumbent (20 hh/1 sp)** | 5′ – CCC TAT TAC CCT TGC CGA CCC TAC CTA CCT ACC CTG CGA GAT GTG AAG TGA ATG ATG AG – 3′ | 59 |
| **Incumbent (20 hh/2 sp)** | 5′ – CCC TAT TAC CCT TGC CGA CCC TAC CTA CCT ACC CTG CGA TGA TGT GAA GTG AAT GAT GAG – 3′ | 60 |
| **Incumbent (20 hh/3 sp)** | 5′ – CCC TAT TAC CCT TGC CGA CCC TAC CTA CCT ACC CTG CGA TAG ATG TGA AGT GAA TGA TGA G – 3′ | 61 |
| **Incumbent (20 hh/5 sp)** | 5′ – CCC TAT TAC CCT TGC CGA CCC TAC CTA CCT ACC CTG CGA TAC CGA TGT GAA GTG AAT GAT GAG – 3′ | 63 |
| **Incumbent (20 hh/8 sp)** | 5′ – CCC TAT TAC CCT TGC CGA CCC TAC CTA CCT ACC CTG CGA TAC CTA AGA TGT GAA GTG AAT GAT GAG – 3′ | 66 |
| **Template A (8 hh/2 sp)** | 5′ – CCG ACC CTA CCT ACC TAC CCT GCG ATG ATG TGA A – 3′ | 34 |
| **Template B (8 hh/2 sp)** | 5′ – CCG ACC CTA CCT ACC TAC CCT GCG ATG ATA GCA T – 3′ | 34 |
| **Template C (8 hh/2 sp)** | 5′ – CCG ACC CTA CCT ACC TAC CCT GCG ATT TAG GTT T – 3′ | 34 |

#### Supplementary Table 21 | Target strands (*T*)

| | Sequence | Length (nt) |
|---|---|---|
| Target (0 th) | 5′ – CGC AGG GTA GGT AGG TAG GGT CGG TAA CTT ATC G – 3′ | 34 |
| Target (1 th) | 5′ – GCG CAG GGT AGG TAG GTA GGG TCG GTA ACT TAT CG – 3′ | 35 |
| Target (2 th) | 5′ – TGC GCA GGG TAG GTA GGT AGG GTC GGT AAC TTA TCG – 3′ | 36 |
| Target (3 th) | 5′ – GTG CGC AGG GTA GGT AGG TAG GGT CGG TAA CTT ATC G – 3′ | 37 |
| Target (5 th) | 5′ – AAG TGC GCA GGG TAG GTA GGT AGG GTC GGT AAC TTA TCG – 3′ | 39 |
| Target (7 th) | 5′ – GGA AGT GCG CAG GGT AGG TAG GTA GGG TCG GTA ACT TAT CG – 3′ | 41 |
| Target (3 th) -$R_2Q_2$ trigger | 5′ – GTG CGC AGG GTA GGT AGG TAG GGT CGG CTC AAC ATA CAT CG – 3′ | 41 |

#### Supplementary Table 22 | Invader strands (*I*)

| | Supplementary Note XXI: Sequence | Length (nt) |
|---|---|---|
| Invader ($I_{hc}$) | 5′ – CTC ATC ATT CAC TTC ACA TCC CGA CCC TAC CTA CCT ACC CTG CGC ACT TCC – 3′ | 51 |
| Invader-random 5′ ($I_{wc}$) | 5′ – ACC TAT AAC CTG AAA CCT AAC CGA CCC TAC CTA CCT ACC CTG CGC ACT TCC – 3′ | 51 |
| Invader-random 3′ | 5′ – CTC ATC ATT CAC TTC ACA TCA ACT CCC ACT CAA CTC ACC TCA ACT CAA CCC – 3′ | 51 |
| Invader A (2 th/ 8 hh) | 5′ – TTA TGC TCC CGA CCC TAC CTA CCT ACC CTG CGC A – 3′ | 34 |
| Invader B (2 th/ 8 hh) | 5′ – ATG CTA TCC CGA CCC TAC CTA CCT ACC CTG CGC A – 3′ | 34 |
| Invader C (2 th/ 8 hh) | 5′ – AAA CCT AAC CGA CCC TAC CTA CCT ACC CTG CGC A – 3′ | 34 |

#### Supplementary Table 23 | Reporter strands

| | Sequence | Length (nt) |
|---|---|---|
| Reporter 1 (*R*) | 5′ – AGG GTC GGC AAG GGT AAT AGG GAC ttt-Cy3 – 3′ | 27 |
| Reporter 1 quencher (*Q*) | 5′ – IowaBlackFQ-ttt GTC CCT ATT ACC CTT GCC – 3′ | 21 |
| $Rep_A$ ($R_2$) | 5′ – Cy5-ttt CGA TGT ATG TTG AGG CGT GCC GTT TTC GGC ACG CGA TGT GAA – 3′ | 45 |
| $Rep_A$ quencher ($Q_2$) | 5′ – CAA CAT ACA TCG ttt-IowaBlackRQ – 3′ | 15 |
| $Rep_B$ fluorescence | 5′ – RhodamineRed™X-ttt GTC GAT GTA TGT TGA GGC GTG CCG TTT TCG GCA CGC GAT AGC AT – 3′ | 47 |
| $Rep_B$ quencher | 5′ – CAA CAT ACA TCG AC ttt-BlackHoleQuencher2 | 17 |
| $Rep_C$ fluorescence | 5′ – AlexaFluor®488-ttt GTC GAT GTA TGT TGA GGC GTG CCG TTT TCG GCA CGC TTA GGT TT – 3′ | 47 |
| $Rep_C$ quencher | 5′ – CAA CAT ACA TCG AC ttt-IowaBlackFQ – 3′ | 17 |

1. Zhang, D.Y. & Winfree, E. Control of DNA Strand Displacement Kinetics Using Toehold Exchange. *J. Am. Chem. Soc.* **131**, 17303-17314 (2009).
2. Liu, W.-T., Wu, J.-H., Li, E.S.-Y. & Selamat, E.S. Emission characteristics of fluorescent labels with respect to temperature changes and subsequent effects on DNA microchip studies. *Appl. Environ. Microbiol.* **71**, 6453-6457 (2005).
3. Moreira, B.G., You, Y., Behlke, M.A. & Owczarzy, R. Effects of fluorescent dyes, quenchers, and dangling ends on DNA duplex stability. *Biochem. Biophys. Res. Commun.* **327**, 473-484 (2005).
4. Lagarias, J.C., Reeds, J.A., Wright, M.H. & Wright, P.E. Convergence properties of the Nelder-Mead simplex method in low dimensions. *SIAM J. Optim.* **9**, 112-147 (1998).
5. SantaLucia, J., Jr. & Hicks, D. The thermodynamics of DNA structural motifs. *Annu. Rev. Biophys. Biomol. Struct.* **33**, 415-440 (2004).
